# Competition with *Pseudomonas aeruginosa* induces *Staphylococcus aureus* in an antibiotic-tolerant viable but non culturable state

**DOI:** 10.1101/2025.04.30.651255

**Authors:** Urszula Łapińska, Nicolas Oswaldo Gomez, Gayathri Chandran, Thomas Tunstall, Sophia Zborowsky, Giulia Tolle, Anthony D. Verderosa, María García-Castillo, Paul A. O’Neill, Audrey Farbos, Aaron R. Jeffries, Andrew J. Young, Rafael Canton, Pierluigi Caboni, Mark A. T. Blaskovich, Adilia Warris, Krasimira Tsaneva-Atanasova, Stefano Pagliara, The ERADIAMR consortium

## Abstract

Interactions between different species of pathogenic microbes often affect clinical outcome by altering the virulence or antibiotic resistance of individual microbes. By investigating the interactions between *Staphylococcus aureus* and *Pseudomonas aeruginosa* at the population, single-cell and molecular level we discovered that a sub-population of *S. aureus* enters in a viable non-culturable state that it is not detected via standard microbiology assays. In the presence of *P. aeruginosa*, *S. aureus* adopts a survival lifestyle similar to previously described intracellular *S. aureus* persisters, downregulating nitrogen metabolism and amino acid biosynthesis, while upregulating protein maturation processes. Entrance in a viable but non culturable state is the primary survival strategy of *S. aureus* in response to vancomycin treatment, whereas only a minority of the *S. aureus* population survive ciprofloxacin treatment while in a viable but non culturable state. These bacterial interactions may shape the evolution of resistance traits of co-infecting pathogens. Manipulating these interspecies adaptations could provide new opportunities for early therapeutic interventions.

## Introduction

Interspecies microbial interactions have shaped the population structure of microbial communities in ecosystems and their biodiversity (*1*). Microbes can engage in mutualistic, neutral or antagonistic interactions, when sharing the same environment (*2*, *3*). In nutrient poor environments lacking essential nutrients, including carbon, nitrogen and iron sources, scavenging is a widely used microbial strategy among bacteria. In these settings bacteria compete for resources (*4*) also via antibiosis, that involves the synthesis of antimicrobial molecules that inhibit growth or kill other bacterial species (*3*).

Interactions between co-colonizing bacteria may influence the development and persistence of co-infections by altering pathogen virulence (*5*) and susceptibility to antibiotics (*6*, *7*), ultimately affecting the health of host humans, animals and plants (*8*). Nevertheless, antibiotic therapies are often designed against individual pathogens without consideration of how interspecies interactions may alter each pathogen susceptibility to antibiotic therapy (*6*). Clinical trials have often reported a lack of association between clinical response to antibiotic therapy and *in vitro* susceptibility testing (i.e. the measurement of an antibiotic minimum inhibitory concentration (MIC) (*9*)), also because these tests are generally carried out against individual microbes (*10*) besides other important factors (*11*)(*12*). Therefore, antibiotic therapy often fails to eradicate co-infecting pathogens (*6*), especially if such pathogens are already intrinsically resistant to key clinical antibiotics (*13*, *14*), thus further exacerbating the current antimicrobial resistance (AMR) crisis (*15*).

Two of the most important bacterial pathogens of humans, the gram-negative *Pseudomonas aeruginosa* and the gram-positive *Staphylococcus aureus*, are frequently co-isolated from infections of catheters, endotracheal tubes, skin, eyes, and the respiratory tract, including the airways of people with cystic fibrosis (CF) (*16*). Indeed, an increased trend in prevalence of *S. aureus* and *P. aeruginosa* co-infections in patients with CF from 30% to 50% was recorded in a longitudinal study between 2004 and 2016 (*17*), although detection frequencies of *S. aureus* and *P. aeruginosa* have decreased in more recent years thanks to the use of elexacaftor, tezacaftor and ivacaftor (*18*). Moreover, a number of studies have shown that co-infection is associated with diminished lung function, more rapid pulmonary decline, delayed wound repair, increased risk of death and increased resistance to antibiotics (*8*, *19–22*), which adds to *S. aureus* and *P. aeruginosa* intrinsic and acquired antibiotic resistance as individual species (*13*, *14*), making these infections difficult to treat (*23*, *24*).

The interactions between *P. aeruginosa* and *S. aureus* are complex. *P. aeruginosa* can display antagonistic interactions with *S. aureus* by secreting molecules that interfere with the growth, metabolism, and cellular homeostasis of *S. aureus*. These molecules include siderophores, such as pyoverdine and pyochelin, that sequester extracellular iron (*25*), proteases such as elastase that cleaves pentaglycine bridges in the *S. aureus* peptidoglycan (*26*), surfactants such as rhamnolipids that interfere with the *S. aureus* cell membrane (*6*), and redox-active secondary metabolites that inhibit *S. aureus* respiration, such as 2-heptyl-4-hydroxyquinoline-N-oxide (HQNO) (*27*), hydrogen cyanide and the phenazine molecule pyocyanin (*8*).

These interactions are strain-dependent (*28*, *29*) and often do not prevent co-colonization *in vivo* (*30*). For example, *P. aeruginosa* isolates from co-infected CF patients are less antagonistic towards *S. aureus* than isolates from mono-infected CF patients (*23*). Moreover, *P. aeruginosa* isolates displaying a mucoid phenotype are less competitive towards *S. aureus* compared to non-mucoid *P. aeruginosa* isolates (*28*) due to alginate overexpression that, in turn, reduces the production of siderophores, (HQNO), and rhamnolipids (*23*). Indeed, a large proportion of *S. aureus* isolates are either unaffected or only growth inhibited by mucoid *P. aeruginosa* (i.e. with a *S. aureus* population reduction smaller than 100-fold) but killed by non-mucoid *P. aeruginosa* (i.e. with a *S. aureus* population reduction larger than 100-fold). A smaller proportion of *S. aureus* isolates are killed by both non-mucoid and mucoid *P. aeruginosa*. Finally, a very small number of *S. aureus* isolates are not killed by either *P. aeruginosa* phenotypes (*28*). On the other hand, the growth of *P. aeruginosa* is generally not affected by the presence of *S. aureus*.

These interactions dramatically alter the susceptibility of *S. aureus* to the killing activities of clinically relevant antibiotics such as vancomycin, ciprofloxacin and tobramycin (*6*, *7*, *10*, *29*, *31–36*). For example, in response to the presence of *P. aeruginosa*, *S. aureus* can upregulate efflux pumps belonging to the Nor family leading to an increase in resistance of *S. aureus* to tetracycline and ciprofloxacin (*29*) or enter in a small-colony-variant (SCV) state becoming resistant to aminoglycosides and beta-lactams (*37*). In contrast, *P. aeruginosa* LasA protease potentiates killing of *S. aureus* by vancomycin, whereas *P. aeruginosa* rhamnolipids facilitate proton-motive force-independent tobramycin uptake in *S. aureus*, thus increasing *S. aureus* susceptibility to these two antibiotics (*6*).

This wealth of knowledge has been obtained by using population level studies, therefore less is known about the interactions between *P. aeruginosa* and *S. aureus* at the sub-population and single-cell level, leaving a gap in our understanding of possible heterogeneities in the strategies adopted by these two bacteria within polymicrobial communities. Here we investigate the dynamics of the interactions between *S. aureus* and *P. aeruginosa* using both laboratory strains and clinical isolates. We employed microbiology, microscopy, transcriptomic and lipidomic analyses to study the dynamics of these interactions at the population, at the single-cell and at the molecular level. We further unravelled the impact of such interactions on antibiotic treatment both at the whole population and at the sub-population level. These early bacterial interactions may shape the evolution, virulence, and resistance of pathogens co-infecting humans and animals and could provide new opportunities for early therapeutic interventions by impeding microbial community adaptations in chronic mixed infections and subsequent patient worsening.

## Results

### *S. aureus* population decline in the continued presence of *P. aeruginosa*

In order to investigate the interactions between *S. aureus* and *P. aeruginosa*, we firstly measured the population dynamics of both bacteria in mono-culture and co-culture. We followed the growth of a methicillin-susceptible *S. aureus* strain (ATCC25923) and a non-mucoid *P. aeruginosa* strain (PA14) in Lysogeny broth (LB) in well-mixed flasks via the colony forming unit (CFU) assay using selective agar plates (i.e. Pseudomonas isolation agar and Trypticase soy agar) (*28*). We found that the growth of *S. aureus* was not affected by the presence of *P. aeruginosa* during the first 4 h of co-culture compared to mono-culture (p-values > 0.05 according to unpaired t-tests at each time point); *S. aureus* growth was significantly reduced between 5 h and 9 h of co-culture compared to mono-culture (p-values < 0.05); at the 9 h time point *S. aureus* population dynamics inverted from expansion to contraction down to a minimum after 13 h of co-culture (Figure 1A and Data A-B in S1 File). In contrast, the growth of *P. aeruginosa* was not affected by the presence of *S. aureus* (Figure 1B and Data C-D in S1 File). We recorded a 200- and 700-fold reduction in *S. aureus* ATCC25923 population after 13 h of co-culture with two different non-mucoid strains of *P. aeruginosa*, i.e. PA14 and PAO1, respectively (Figure S1 and Data E in S1 File).

**Figure 1.**
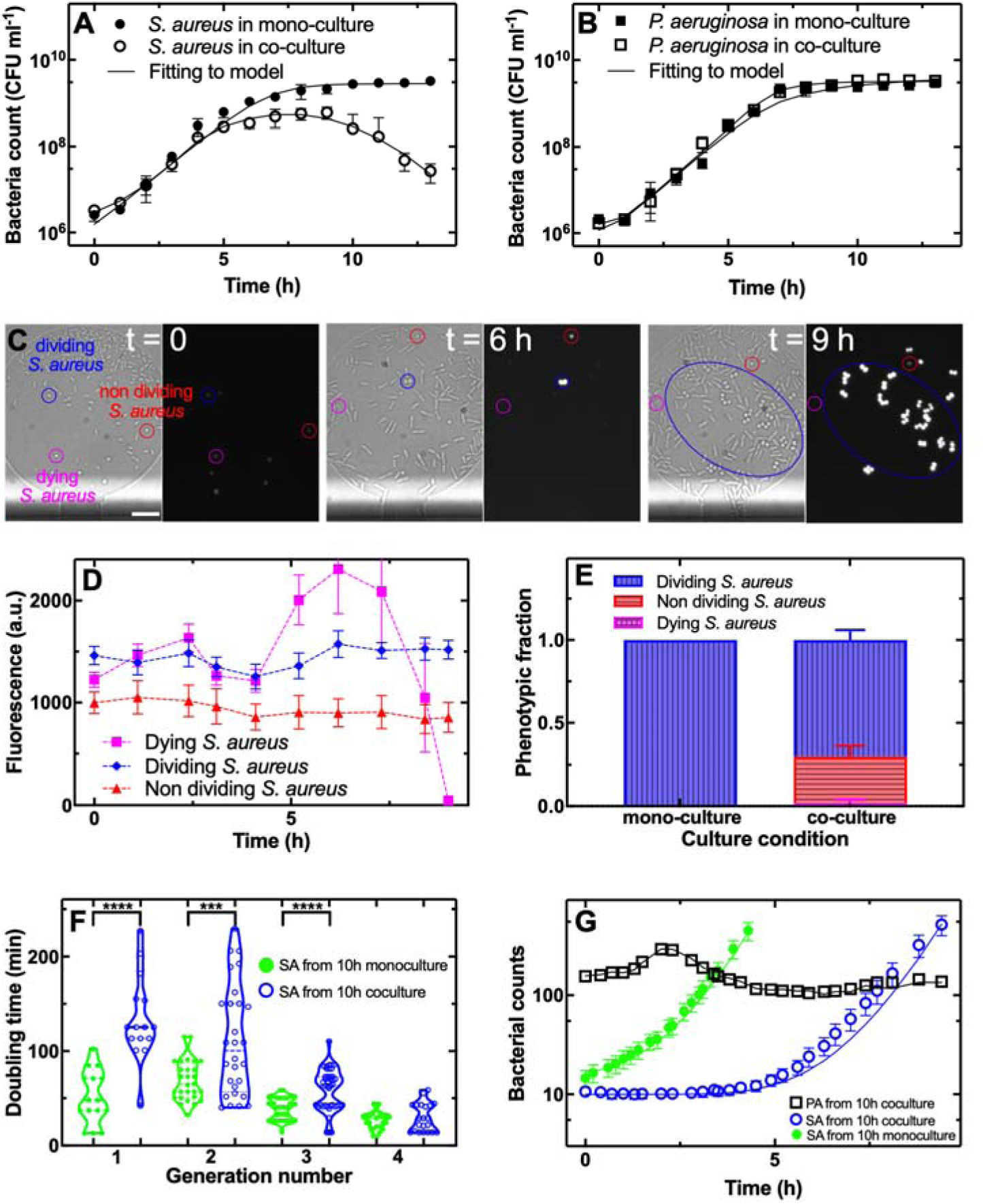
A sub-population of *S. aureus* enters a viable but non culturable state in the presence of *P. aeruginosa*. (a-b) Temporal dependence of bacterial population size for (a) *S. aureus* ATCC25923 growing in mono-culture (filled circles) or in co-culture with *P. aeruginosa* PA14 in well-mixed flasks (open circles) and (b) for *P. aeruginosa* PA14 growing in mono-culture (filled squares) or in co-culture with *S. aureus* ATCC25923 in well-mixed flasks (open squares). Data points are the mean and standard deviation of colony forming unit (CFU) measurements carried out in biological triplicate each consisting of technical duplicate. Numerical values for each replicate are reported in Data A-D in S1 File. The solid lines are fitting of these data to our mathematical model with parameter estimates reported in Table S1. (c) Brightfield and fluorescence microscopy images of *S. aureus* SH1000 mCherry and *P. aeruginosa* PA14 Δ*flgK* introduced in microfluidic chambers after 10 h of co-culture in flasks and after regrowth in the microfluidic chambers for 0, 6 h or 9 h. One representative dividing, non-dividing or dying *S. aureus* SH1000 mCherry is denoted with a blue, red or magenta circle, respectively. Scale bar: 10 µm. (d) Temporal dependence of single-cell mCherry fluorescence for dividing, non-dividing or dying *S. aureus* (blue, red or magenta data points, respectively). Data points are the mean and standard error of ten single-cell measurements from biological triplicate experiments. Numerical values are reported in Data M in S1 File. (e) Corresponding fraction of dividing (vertically patterned bars), non-dividing (horizontally patterned bars), or dying (checkered patterned bars) *S. aureus* in mono-culture or in co-culture with *P. aeruginosa* within microfluidic chambers. (f) Corresponding dependence of doubling time on the generation number for *S. aureus* in mono-culture (green filled circles) or in co-culture with *P. aeruginosa* (open blue circles) within microfluidic chambers. Each point reports the doubling time of an individual bacterium with numerical values reported in Data Q in S1 File. *** denotes a p-value < 0.001, **** denotes a p-value < 0.0001. (g) Corresponding temporal dependence of bacterial population size for *S. aureus* growing in mono-culture (filled green circles) or in co-culture with *P. aeruginosa* (open blue circles) and for *P. aeruginosa* growing in co-culture (open black squares) in microfluidic chambers inoculated with bacteria after 10 h of culture in well-mixed flasks. Data points are the mean and standard error of cell count measurements carried out in biological triplicate experiments each consisting of four microfluidic chambers. Numerical values for each replicate are reported in Data N-P in S1 File. The solid lines are fitting of these data to our mathematical model with parameter estimates reported in Table S2 and Table S3.

Next, we investigated the interactions between methicillin-resistant *S. aureus* isolate RYC157 and *P. aeruginosa* isolate RYC157 co-isolated from a dual-species infection in a CF patient. We recorded a slower growth of *P. aeruginosa* RYC157 compared to both *S. aureus* RYC157 and *P. aeruginosa* PA14 and we observed a population decline in *S. aureus* RYC157 only after 20 h of co-culture (Figure S2B and Data F and G in S2 File). Moreover, we observed a population decline in both *S. aureus* RYC157 and *S. aureus* RYC165 (a different methicillin-resistant *S. aureus* strain co-isolated from a dual-species infection with *P. aeruginosa* also in a CF patient) after 9 h and 10 h of co-culture with *P. aeruginosa* PA14 (Figure S2C and S2D and Data H-K in S1 File) consistent with the data obtained for *S. aureus* ATCC25923.

These data suggest that *P. aeruginosa* PA14 kills *S. aureus* ATCC25923, RYC157 and RYC165, considering that previous co-culture assays ascribed to killing bacterial population reductions larger than 100-fold (*28*, *38*) and that we measured an inversion in the *S. aureus* population dynamics. However, it is also conceivable that in the continued presence of *P. aeruginosa*, *S. aureus* progressively enters a viable but non-culturable state (*39*) and that individual *S. aureus* cells do not form colonies on selective agar plates, thus leading to an underestimation of the *S. aureus* population.

In order to rationalise these population dynamics, we developed a mathematical model based on the simplified assumptions that the growth rates of *S. aureus* and *P. aeruginosa* are modulated by the proportion of active biochemical processes which contribute towards replication, and that *P. aeruginosa* secretes molecules that can either kill or induce a non-culturable state in *S. aureus* (see Methods). By fitting this model to our population dynamics data via least-squares regression, we were able to capture the experimentally measured population dynamics for the different *S. aureus* and *P. aeruginosa* strains in both mono- and co-culture (Figure 1A, 1B and S2) with parameter estimations reported in Table S1. Taken together, these data and model suggest that the decline observed for *S. aureus* in co-culture is due to either *S. aureus* death or entrance in a viable non-culturable state.

### A sub-population of *S. aureus* enters a viable but non-culturable state in the presence of *P. aeruginosa*

In order to test the non-mutually exclusive hypotheses above, we employed our previously described single-cell microfluidics-based time-lapse microscopy platform(*40–42*) with bacteria hosted in 1 µm high and 150 µm wide circular chambers connected to a 20 µm high and 200 µm wide main delivery chamber. To facilitate long-term time-lapse microscopy of individual bacteria hosted within these chambers, we used a *P. aeruginosa* deletion mutant with reduced motility due to a lack of flagella (PA14 Δ*flgK*(*43*)) and a *S. aureus* fluorescent reporter strain (SH1000 mCherry (*44*)) to facilitate distinguishing between viable and non-viable *S. aureus* cells. We verified that after 13 h of co-culture with *P. aeruginosa* PA14 Δ*flgK* in well-mixed flasks, both *S. aureus* ATCC 25923 and *S. aureus* SH1000 mCherry displayed a significant reduction in population size with respect to *S. aureus* mono-cultures, similar to the reduction in population size observed after co-culture with *P. aeruginosa* PA14 or PAO1 (Figure S1 and Data E in S1 File). These data confirm that the antagonistic interactions recorded for the pairs *P. aeruginosa* PA14 (or PAO1) and *S. aureus* ATCC 25923 are well recapitulated when using the pair *P. aeruginosa* PA14 Δ*flgK* and *S. aureus* SH1000 mCherry.

Therefore, we harvested a 100 µL aliquot from a *S. aureus* SH1000 mCherry and *P. aeruginosa* PA14 Δ*flgK* co-culture after 10 h of growth in a well-mixed flask and inoculated it in four microfluidic chambers continuously supplied via the main delivery chamber with LB for 9 h. We imaged each chamber every 10 min for 9 h and repeated this experiment in biological triplicate using three independent 10 h old co-cultures as inoculum. By tracking individual *S. aureus* cells in each chamber in each experiment over time (Figure 1C and 1D), we found that 70% of *S. aureus* cells resumed growth in the presence of *P. aeruginosa*, 28% of *S. aureus* did not resume growth within the 9 h exposure to LB, and 2% of *S. aureus* cells died (Figure 1E and Data L in S1 File). *S. aureus* cells that did not resume growth and division retained motility, exploring the microfluidic chambers during the 9 h exposure to LB (Figure 1C). Furthermore, these viable but non culturable cells displayed a significantly lower mCherry fluorescence at t = 0 compared to *S. aureus* cells that than resumed growth, but maintained stable mCherry fluorescence over time and did not stain with propidium iodide similarly to dividing cells (Figure 1D and Data M in S1 File). In contrast, *S. aureus* cells that eventually died while in co-culture with *P. aeruginosa*, displayed a sharp decrease in mCherry fluorescence (Figure 1D) and stained with propidium iodide.

The ratio of dividing over non-dividing *S. aureus* cells varied across the different microfluidic chambers investigated but did not depend on the initial ratio of *P. aeruginosa* cell number over *S. aureus* cell number in each chamber (Pearson correlation coefficient r = 0.03, p-value = 0.92, Figure S3 and Data O in S1 File). In contrast, 100% of *S. aureus* cells resumed growth upon inoculation in the microfluidic chambers after 10 h of mono-culture in well-mixed flasks (Figure 1E and Data N in S1 File). Moreover, the lag times of individual *S. aureus* cells in co-culture with *P. aeruginosa* were significantly longer and more heterogeneous compared with the lag times of *S. aureus* cells in mono-culture (1^st^ generation in Figure 1F). Similarly, the doubling times of the 2^nd^ and 3^rd^ generation *S. aureus* daughter cells in co-culture were significantly longer and more heterogeneous compared with the doubling times of *S. aureus* cells in mono-culture and only 4^th^ generation daughter cells doubled at similar rates in mono- and co-culture (Figure 1F and Data Q in S1 File).

### Impact of the *S. aureus* viable but non-culturable state on population dynamics

As a result, the population dynamics of *S. aureus* in co-culture with *P. aeruginosa* in microfluidic chambers was different with respect to that measured for *S. aureus* in mono-culture in microfluidic chambers. In mono-culture, the *S. aureus* population in each chamber readily and homogeneously expanded exponentially, variations in cell number per chamber being dictated by variant initial inoculum per chamber in range from 5 to 30 cells (Figure 1G, Figure S4A and Data N in S1 File). In contrast, the *S. aureus* population started to increase only after 5 h of co-culture with *P. aeruginosa* and growth was heterogeneous across the different chambers (Figure 1G, Figure S4B and Data O in S1 File). The expansion in *S. aureus* population was preceded by an inversion in the trend of the *P. aeruginosa* population dynamics that expanded during the first 2.5 h of co-culture with *S. aureus* and decreased from the 2.5 h time point onwards (Figure 1G, Figure S4C and Data P in S1 File), due to escape out of the chambers, possibly linked to increased cellular motility via type IV pili (*43*). As a consequence, the ratio of *P. aeruginosa* cell number over *S. aureus* cell number per chamber was initially between 10 and 25 (upon inoculation after 10 h of co-culture in well-mixed flasks), it increased to a maximum of 35 after 2.5 h of co-culture in the microfluidic chambers and then decreased down to a minimum that was lower than 1 in most of the microfluidic chambers investigated (Figure S4D). These distinct populations dynamics of *S. aureus* and *P. aeruginosa* in mono- or co-culture were well described by a mathematical model that accounts for the resumption of intracellular activity and bacterial growth and the fact that *P. aeruginosa* cells are able to escape the microfluidic chambers (see Methods and Figure 1G) with parameter estimations reported in Table S2 and Table S3.

In order to further test whether the non-dividing *S. aureus* sub-population had an impact on *S. aureus* population dynamics, we used both LB and selective agar plates (i.e. Pseudomonas isolation agar and Trypticase soy agar) (*28*) to enumerate *S. aureus* and *P. aeruginosa* after co-culture in well-mixed flasks. In accordance with our microscopy data, we found that *S. aureus* counts on selective plates were higher than counts on LB agar plates (Figure S5 and Data B, D and R-S in S1 File), possibly due to a larger sub-population of *S. aureus* remaining in a non-dividing state on LB agar plates, where also *P. aeruginosa* can grow compared to selective plates where *P. aeruginosa* cannot grow.

Taken together these data demonstrate that the presence of *P. aeruginosa* induces a sub-population of *S. aureus* in a viable but non culturable state that is not detectable using standard microbiology techniques, such as the colony forming unit assay, leading to a significant decrease of the culturable *S. aureus* population over time but to a limited decrease of the viable *S. aureus* population.

### *P. aeruginosa* rewires its transcriptome faster than *S. aureus* in co-culture

In order to investigate the molecular mechanisms involved in the interactions between *S. aureus* and *P. aeruginosa*, we employed an unbiased approach to examine changes in gene expression by performing genome-wide comparative transcriptome analysis between mono-and co-cultures (*25*, *45*). Specifically, we extracted RNA from *S. aureus* ATCC25923 and *P. aeruginosa* PA14 from biological triplicates after 3 h of mono- or co-culture. At this time point, the population size of *S. aureus* from the co-culture was only 1.5-fold lower compared to the population size of *S. aureus* from the mono-culture, suggesting that effects of competition had not fully translated into changes in population dynamics (*46*), and both *S. aureus* and *P. aeruginosa* were in their exponential phase of growth (Figure 1A-1B). Moreover, we also extracted RNA from *S. aureus* and *P. aeruginosa* after 10 h of mono- and co-culture. At this time point, the population size of *S. aureus* from the co-culture was 10-fold lower compared to the population size of *S. aureus* from the mono-culture, suggesting that effects of competition had fully translated into changes in population dynamics, and both *S. aureus* and *P. aeruginosa* mono-cultures were in their stationary phase of growth (Figure 1A-1B).

Principal component analysis revealed both major divergences across different culturing conditions and high within group reproducibility, i.e. transcriptome replicates from each condition clustered together. We found that the transcriptomes of *S. aureus* after 3 h in co-culture were close to those measured for *S. aureus* after 3 h in mono-culture confirming that by this time point *P. aeruginosa* did not have a significant impact on *S. aureus* (Figure S6A and S2 File). In contrast, the transcriptomes of *S. aureus* after 10 h in co-culture were very different, in both principal components, to those measured for *S. aureus* after 10 h in mono-culture confirming that by this time point *P. aeruginosa* did have a significant impact on *S. aureus* (Figure S6A). Moreover, the transcriptomes of *P. aeruginosa* after 10 h in co-culture were different to those measured for *P. aeruginosa* after 10 h in mono-culture in terms of principal component 2 (Figure S6B and S2 File).

Next, we examined changes in gene expression between *S. aureus* and *P. aeruginosa* co-cultures and mono-cultures at both time points by using a conservative cut-off of 2-fold change (i.e. log_2_ fold change > 1 or log_2_ fold change < -1) and adjusted p-value (via Benjamini-Hochberg) smaller than 0.05 to identify differentially expressed genes (*46*, *47*). This analysis revealed that *P. aeruginosa* reacted faster to the presence of the other species, whereas *S. aureus* responded at a later stage: the number of significantly differentially regulated *P. aeruginosa* genes decreased from 397 to 250 (i.e. from 7% to 4% of the genome) from 3 h to 10 h of co-culture with respect to mono-culture; in striking contrast, the number of significantly differentially regulated *S. aureus* genes dramatically increased from 153 to 814 (i.e. from 6% to 33% of the genome) from 3 h to 10 h of co-culture with respect to mono-culture (Figure S7 and S3-S6 Files, RNA transcript counts for all genes and individual replica are reported in S2 File).

### *P. aeruginosa* upregulates the expression of the pyochelin pathway and DNA-templated transcription initiation in the continued presence of *S. aureus*

Next, we performed gene ontology enrichment analysis on each subset of differentially regulated genes between mono- and co-culture and identified biological processes that were significantly upregulated or downregulated across each comparison (i.e. with an enrichment score larger or smaller than zero, respectively (*47–49*)). We found that quinone biosynthesis, cellular respiration and molybdopterin cofactor metabolism processes were upregulated in *P. aeruginosa* in co-culture with respect to mono-culture at t = 3 h, whereas membrane transport, carbohydrate and fatty acid biosynthesis and DNA-templated transcription initiation processes were downregulated (Figure 2A and S7 File). At t = 10 h instead iron transport, DNA-templated transcription initiation, protein folding, peptide and dicarboxylic acid metabolism processes were upregulated, whereas biosynthesis regulation, homeostasis and carbohydrate metabolism processes were downregulated in *P. aeruginosa* in co-culture with respect to mono-culture (Figure 2B and S8 File).

**Figure 2.**
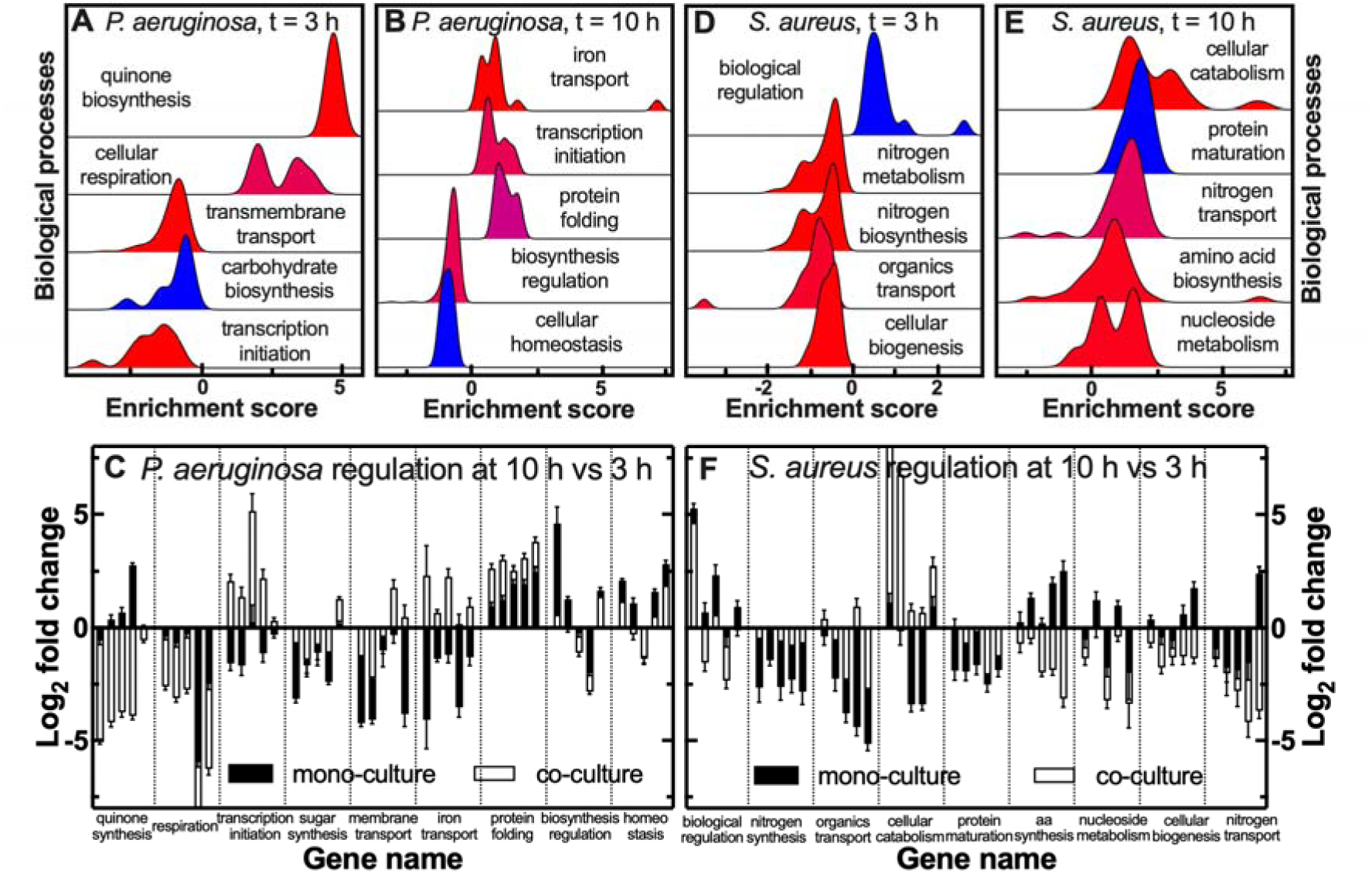
Molecular interactions between *P. aeruginosa* and *S. aureus*. (a and b) Biological processes that are significantly enriched in the comparisons of the transcriptomes of *P. aeruginosa* in mono-culture vs co-culture at (a) t = 3 h or (b) at t = 10 h. (d and e). Biological processes that are significantly enriched in the comparisons of the transcriptomes of *S. aureus* in mono-culture vs co-culture at (d) t = 3 h or (e) at t = 10 h. The colour coding indicates the calculated adjusted p-values reported in S7 and S8 File, red indicating the lowest and blue indicating the highest adjusted p-value. (c and f) Differential expression of (c) *P. aeruginosa* and (f) *S. aureus* biological processes at t = 10 h compared to t = 3 h in mono-culture (filled bars) or co-culture (open bars). The bars and error bars report the mean and standard error of the transcript log_2_ fold change (calculated from biological triplicate transcriptomic measurements) of five representative genes for each pathway. log_2_ fold change data for all genes in each pathway are reported in Figure S8 and Figure S12 with numerical values reported in S9-S10 and S14-S15 Files.

Next, we compared the temporal regulation of these processes in *P. aeruginosa* in mono-culture and co-culture. Genes in the quinone biosynthesis, cellular respiration and molybdopterin cofactor metabolism processes were significantly more downregulated at t =10 h vs t = 3 h in co-culture compared to mono-culture. In contrast, genes in the membrane transport and peptide metabolism processes were significantly more downregulated at t = 10 h in mono-culture compared to co-culture (Figure 2C, Figure S8 and S9-S10 Files).

Genes in the regulation of nitrogen metabolism, carbohydrate metabolism and homeostasis were significantly more upregulated at t =10 h in mono-culture compared to co-culture. In contrast, genes in the carbohydrate biosynthesis and protein folding processes were significantly more upregulated at t =10 h in co-culture compared to mono-culture (Figure 2C, Figure S8 and S9-S10 Files).

Strikingly, genes in the DNA-templated transcription initiation (including the sigma factor *pvdS*), iron transport and dicarboxylic acid metabolism processes displayed opposite regulation when *P. aeruginosa* was in co-culture compared to mono-culture: in mono-culture these genes were downregulated at t = 10 h compared to t = 3 h, whereas in co-culture these genes were upregulated at t = 10 h compared to t = 3h (Figure 2C, Figure S8 and S9-S10 Files). Notably, *P. aeruginosa* employs two siderophores, pyochelin and pyoverdine, for the uptake of iron and other metals. The pyoverdine synthesis pathway was upregulated at t = 10 h compared to t = 3 h both in mono-culture and co-culture (Figure S9A); in striking contrast, the pyochelin synthesis pathway was strongly downregulated at t = 10 h compared to t = 3 h in mono-culture and strongly upregulated at t = 10 h compared to t = 3 h in co-culture (Figure S9B and S9-S10 Files), suggesting that secretion of pyochelin, but not of pyoverdine, is a *P. aeruginosa* response to the presence of *S. aureus*. We assessed the effect of pyochelin on *S. aureus* viability by co-culturing *S. aureus* in the presence of a *P. aeruginosa* siderophore-negative knockout mutant (PAO1 Δ*pvdD*Δ*pchEF* (*50*)). A similar reduction in *S. aureus* population was seen as with the *P. aeruginosa* PAO1 parental strain, (Figure S1), suggesting that siderophore secretion alone was not responsible for the observed inversion in the population dynamics of *S. aureus*.

Pathways previously implicated in the response of *P. aeruginosa* to the presence of *S. aureus*, i.e. the phenazine, HQNO, hydrogen cyanide, alginate, rhamnolipid, exopolysaccharide, or elastase pathways and the quorum sensing master regulators *lasR*, *rhlR* and *mvfR* (*6*, *8*, *26*, *27*, *46*, *51*) displayed a similar temporal regulation in co-culture and mono-culture (Figure S10 and S9-S10 Files), suggesting that these pathways are not induced by the presence of *S. aureus*. These RNA-seq data were further validated by quantitative real-time PCR for twenty genes (including genes in the *P. aeruginosa* pyoverdine, pyochelin and HQNO pathways), showing a strong correlation with the transcriptome data (Figure S11 and S11 File).

Taken together, these data suggest that after prolonged exposure to *S. aureus*, *P. aeruginosa* transports more substrates, particularly iron, performs more transcription initiation, protein repair and carbohydrate biosynthesis, while reducing respiration and control over biosynthesis and homeostasis.

### *S. aureus* increases the expression of cellular catabolism and chaperone genes in the continued presence of *P. aeruginosa*

Gene ontology enrichment analysis revealed that biological regulation was the only process upregulated in *S. aureus* in co-culture with respect to mono-culture at t = 3 h, whereas nitrogen metabolism and biosynthesis, organic substance transport and cellular organization processes were downregulated (Figure 2D and S12 File). At t = 10 h instead cellular catabolism and protein maturation processes were upregulated, whereas alpha-amino acid biosynthesis, nucleoside metabolism, biogenesis and nitrogen transport processes were downregulated in *S. aureus* in co-culture with respect to mono-culture (Figure 2E and S13 File).

Next, we compared the temporal regulation of these processes in *S. aureus* in mono-culture and co-culture. Genes in the protein maturation process were significantly more downregulated at t = 10 h in mono-culture than in co-culture. In contrast, genes in the nucleoside monophosphate metabolism and cellular biogenesis processes were significantly more downregulated at t = 10 h in co-culture than in mono-culture (Figure 2E, Figure S12 and S14-S15 Files).

Genes in the cellular catabolism process displayed a remarkably opposite regulation in co-culture with respect to mono-culture: these genes were downregulated at t = 10 h compared to t = 3h in mono-culture, whereas were upregulated in co-culture. The opposite was true for genes in the alpha-amino acids biosynthesis process that were upregulated at t = 10 h compared to t = 3h in mono-culture, whereas were downregulated in co-culture (Figure 2E, Figure S12 and S14-S15 Files). These RNA-seq data were further validated by quantitative real-time PCR for twenty genes, showing a strong correlation with the transcriptome data (Figure S11 and S11 File).

Taken together, these data suggest that after prolonged exposure to *P. aeruginosa*, *S. aureus* transports less nitrogen, produces less nucleosides and amino acids and reduces biogenesis, while increasing its catabolic rates and chaperone activity, possibly to reduce protein aggregation that is linked to bacterial dormancy (*42*, *52*). These features are consistent with our data showing that sub-populations of *S. aureus* enter in a viable but non-culturable state during co-culture with *P. aeruginosa* (Figure 1). Moreover, our transcriptomic analysis also revealed that in the presence of *P. aeruginosa*, *S. aureus* downregulated the biosynthesis of key amino acids, such as valine, leucine, isoleucine, lysine and tryptophan, and aminoacyl-tRNA synthetases, whereas *S. aureus* upregulated the heat shock stimulon, the *codY* regulon and the ribosome recycling gene *frr* (S6 File), similarly to gene expression changes previously reported for intracellular persisters (*53*), a comparison on which we expand in the Discussion.

### During co-culture *P. aeruginosa* and *S. aureus* upregulate phosphatidylglycerol lipids

Next, we used ultra-high performance liquid chromatography-quadrupole time-of-flight mass spectrometry (*47*, *54*) to quantify the relative abundance of membrane lipids in *P. aeruginosa* PA14 and *S. aureus* ATCC25923 after 10 h of co-culture with respect to after 10 h mono-cultures, all comparisons having being performed in biological triplicates. Within the lipidomes extracted from the co-cultures, lipids could not be assigned unambiguously to *P. aeruginosa* or *S. aureus*; therefore, Table 1 reports quantitative information only on lipids that were detected in either species in mono-cultures and in co-cultures.

**Table 1.** Lipids differentially regulated in co-cultures compared with mono-cultures. . Lipid class, fatty acid composition, adduct type, retention time (RT), theoretical and experimental mass-to-charge ratio (m/z), molecular formula, variable importance in projection (VIP) score, and mass error (Δppm) measured via quantitative time-of-flight liquid chromatography mass spectrometry.

| Lipid | Composition | Adduct | RT<br>(min) | Theor. mass<br>(m/z) | Exp. mass (m/z) | Formula | VIP<br>Score | Δppm |
| --- | --- | --- | --- | --- | --- | --- | --- | --- |
| **Lipids upregulated after 10 h co-culture compared to *P. aeruginosa* mono-culture** |  |  |  |  |  |  |  |  |
| MGDG 29:0 | 13:0;16:0 | -C<sub>2</sub>O<sub>2</sub>H<sub>4</sub><sup>-</sup> | 3.35 | 748.5337 | 748.5342 | C<sub>38</sub>H<sub>72</sub>O<sub>10</sub> | 1.46 | -0.66 |
| MGDG 29:1 | 13:0;16:1 | -C<sub>2</sub>O<sub>2</sub>H<sub>4</sub><sup>-</sup> | 2.81 | 746.5180 | 746.5186 | C<sub>38</sub>H<sub>70</sub>O<sub>10</sub> | 1.40 | -0.80 |
| MGDG 30:0 | 14:0;16:0 | -C<sub>2</sub>O<sub>2</sub>H<sub>4</sub><sup>-</sup> | 3.86 | 750.5495 | 750.5499 | C<sub>38</sub>H<sub>74</sub>O<sub>10</sub> | 1.10 | -0.53 |
| PG 33:0 | 16:0;17:0 | -H<sup>+</sup> | 3.61 | 735.5227 | 735.5223 | C<sub>39</sub>H<sub>77</sub>O<sub>10</sub>P | 1.74 | 0.54 |
| PG 34:0 | 16:0;18:0 | -H<sup>+</sup> | 4.11 | 749.5341 | 749.5340 | C<sub>40</sub>H<sub>79</sub>O<sub>10</sub>P | 1.54 | 0.13 |
| PG 35:0 | 16:0;19:0 | -H<sup>+</sup> | 4.40 | 763.5511 | 763.5541 | C<sub>41</sub>H<sub>81</sub>O<sub>10</sub>P | 1.40 | -3.92 |
| PE 32:0 | 16:0;16:0 | -H<sup>+</sup> | 4.05 | 690.5123 | 690.5120 | C<sub>37</sub>H<sub>74</sub>NO<sub>8</sub>P | 1.02 | 0.43 |
| PE 34:0 | 16:0;18:0 | -H<sup>+</sup> | 5.02 | 718.5436 | 718.5432 | C<sub>39</sub>H<sub>78</sub>NO<sub>8</sub>P | 1.30 | 0.55 |
| **Lipids downregulated after 10 h co-culture compared to *P. aeruginosa* mono-culture** |  |  |  |  |  |  |  |  |
| PC 34:1 | 16:0;18:1 | -H<sup>+</sup> | 4.05 | 758.5705 | 758.5700 | C<sub>42</sub>H<sub>82</sub>NO<sub>8</sub>P | 1.10 | 0.65 |
| PC 34:0 | 16:0;18:0 | +H<sup>+</sup> | 5.01 | 762.6007 | 762.6001 | C<sub>42</sub>H<sub>84</sub>NO<sub>8</sub>P | 1.12 | 0.78 |
| PC 35:1 | 16:0;19:1 | +H<sup>+</sup> | 4.66 | 774.6007 | 774.6005 | C<sub>43</sub>H<sub>84</sub>NO<sub>8</sub>P | 1.02 | 0.25 |
| PC 36:2 | 18:1;18:1 | +H<sup>+</sup> | 4.14 | 786.6007 | 786.6004 | C<sub>44</sub>H<sub>84</sub>NO<sub>8</sub>P | 1.07 | 0.38 |
| PE 33:1 | 16:0;17:1 | +H<sup>+</sup> | 3.72 | 704.5225 | 704.5219 | C<sub>38</sub>H<sub>74</sub>NO<sub>8</sub>P | 1.00 | 0.85 |
| PE 36:1 | 18:0;18:1 | +H<sup>+</sup> | 5.13 | 746.5694 | 746.5687 | C<sub>41</sub>H<sub>80</sub>NO<sub>8</sub>P | 1.09 | 0.93 |
| PE 36:2 | 18:1;18:1 | +H<sup>+</sup> | 4.22 | 744.5538 | 744.5537 | C<sub>41</sub>H<sub>78</sub>NO<sub>8</sub>P | 1.21 | 0.13 |
| **Lipids upregulated after 10 h co-culture compared to *S. aureus* mono-culture** |  |  |  |  |  |  |  |  |
| PG 30:0 | 16:0;14:0 | -H<sup>+</sup> | 2.47 | 693.4752 | 693.4755 | C<sub>36</sub>H<sub>71</sub>O<sub>10</sub>P | 1.28 | -0.43 |
| PG 31:0 | 16:0;15:0 | -H<sup>+</sup> | 2.85 | 707.4807 | 707.4808 | C<sub>37</sub>H<sub>73</sub>O<sub>10</sub>P | 1.13 | -0.14 |
| PG 32:0 | 15:0;17:0 | -H<sup>+</sup> | 3.08 | 721.5060 | 721.5076 | C<sub>38</sub>H<sub>75</sub>O<sub>10</sub>P | 1.27 | -2.21 |
| PG 33:0 | 16:0;17:0 | -H<sup>+</sup> | 3.61 | 735.5172 | 735.5177 | C<sub>39</sub>H<sub>77</sub>O<sub>10</sub>P | 1.05 | -0.67 |
| PG 34:0 | 16:0;18:0 | -H<sup>+</sup> | 4.11 | 749.5376 | 749.5382 | C<sub>40</sub>H<sub>79</sub>O<sub>10</sub>P | 1.24 | -0.80 |
| PG 35:0 | 16:0;19:0 | -H<sup>-</sup> | 4.40 | 763.5511 | 763.5541 | C<sub>41</sub>H<sub>81</sub>O<sub>10</sub>P | 1.27 | -3.92 |
| PG 36:0 | 18:0;18:0 | -H<sup>-</sup> | 4.71 | 777.5634 | 777.5688 | C<sub>42</sub>H<sub>83</sub>O<sub>10</sub>P | 1.10 | -6.94 |
| FA | 16:0 | -H<sup>-</sup> | 1.56 | 255.2330 | 255.2342 | C<sub>16</sub>H<sub>32</sub>O<sub>2</sub> | 1.11 | -4.70 |
| FA | 18:0 | -H<sup>-</sup> | 2.14 | 283.2643 | 283.2647 | C<sub>16</sub>H<sub>36</sub>O<sub>2</sub> | 1.26 | -1.41 |
| **Lipids downregulated after 10 h co-culture compared to *S. aureus* mono-culture** |  |  |  |  |  |  |  |  |
| DGDG 32:0 | 15:0;17:0 | -H<sup>-</sup> | 3.40 | 937.6157 | 937.6154 | C<sub>47</sub>H<sub>88</sub>O<sub>15</sub> | 1.04 | 0.31 |
| DGDG 35:0 | 15:0;20:0 | -H<sup>-</sup> | 4.84 | 979.6621 | 979.6620 | C<sub>50</sub>H<sub>94</sub>O<sub>15</sub> | 1.06 | 0.10 |
| LPC | 16:0 | +H<sup>+</sup> | 0.97 | 496.3398 | 496.3391 | C<sub>24</sub>H<sub>50</sub>NO<sub>7</sub>P | 1.10 | 1.41 |
| LPC | 18:0 | +H<sup>+</sup> | 1.19 | 524.3711 | 524.3702 | C<sub>26</sub>H<sub>54</sub>NO<sub>7</sub>P | 1.03 | 1.71 |
| LPC | 18:1 | +H<sup>+</sup> | 1.00 | 522.3554 | 522.3546 | C<sub>26</sub>H<sub>52</sub>NO<sub>7</sub>P | 1.05 | 1.53 |

We found that after 10 h of co-culture, the expression of phosphatidylglycerol (PG) and monogalactosyl diacylglycerol (MGDG) lipids was up regulated and the expression of phosphatidylcholine (PC) lipids was downregulated with respect to 10 h of *P. aeruginosa* mono-culture. The regulation of phosphatidylethanolamine (PE) lipids was instead mixed with PE 32:0 and PE34:0 being upregulated in co-culture and PE 33:1, PE 36:1 and PE 36:2 being downregulated in co-culture (Table 1).

Furthermore, the expression of a large subset of PG lipids and some fatty acids was upregulated after 10 h of co-culture with respect to 10 h of *S. aureus* mono-culture, whereas the expression of glycosyldiacylglycerol (DGDG) and lysophosphatidylcholine lipids (LPC) was downregulated in co-culture with respect to 10 h of *S. aureus* mono-culture (Table 1).

Taken together, these data suggest that phosphatidylglycerol lipids are upregulated in both *P. aeruginosa* and *S. aureus* after 10 h of co-culture, whereas phosphatidylcholine lipids are downregulated in both *P. aeruginosa* and *S. aureus* after 10 h of co-culture with respect to the corresponding mono-cultures.

### The presence of *P. aeruginosa* decreases the bactericidal efficacy of ciprofloxacin against *S. aureus* when used at supra-inhibitory concentrations

Next, we set out to determine the impact of the interactions between *S. aureus* and *P. aeruginosa* on antibiotic treatment. We employed two antibiotics that are routinely used for the treatment of cystic fibrosis patients: ciprofloxacin, a quinolone that has bactericidal activity against both *S. aureus* and *P. aeruginosa* and is commonly used to treat *P. aeruginosa* during coinfection; vancomycin, a frontline glycopeptide that is used to treat methicillin-resistant *S. aureus* in cystic fibrosis patients, but it is not effective against *P. aeruginosa* (*6*). We performed time-killing assays (*49*) trying to reflect an infection-like condition wherein the microbial community is already established prior to the application of any treatment (*31*). Therefore, we added ciprofloxacin or vancomycin to 3 h old mono- or co-cultures, i.e. in the exponential phase of growth and before we recorded an impact of *P. aeruginosa* PA14 presence on *S. aureus* ATCC25923 growth, or to 10 h old mono- or co-cultures, i.e. in the stationary phase of growth and after we recorded an impact of *P. aeruginosa* presence on *S. aureus* growth.

The minimum inhibitory concentration (MIC) of ciprofloxacin against *P. aeruginosa* and *S. aureus* was 0.125 µg mL^-1^ and 0.5 µg mL^-1^, respectively. Therefore, when we performed time-killing assays using ciprofloxacin at the MIC value measured for *S. aureus* (i.e. 0.5 µg mL^-1^), we found that ciprofloxacin had a three or one orders of magnitude lower efficacy against *S. aureus* compared to *P. aeruginosa* mono-cultures in the exponential (with a 3-log vs a 6-log reduction) or stationary (with a 2-log vs a 3-log) phase of growth (Figure 3A-B and S16 File). Consistently, when ciprofloxacin was employed at 10× or 25× its MIC (against *S. aureus*) it had a weaker efficacy against *S. aureus* compared to *P. aeruginosa* mono-cultures in the stationary phase of growth (with a 4-log vs a 6-log reduction, respectively, Figure S13E-H).

**Figure 3.**
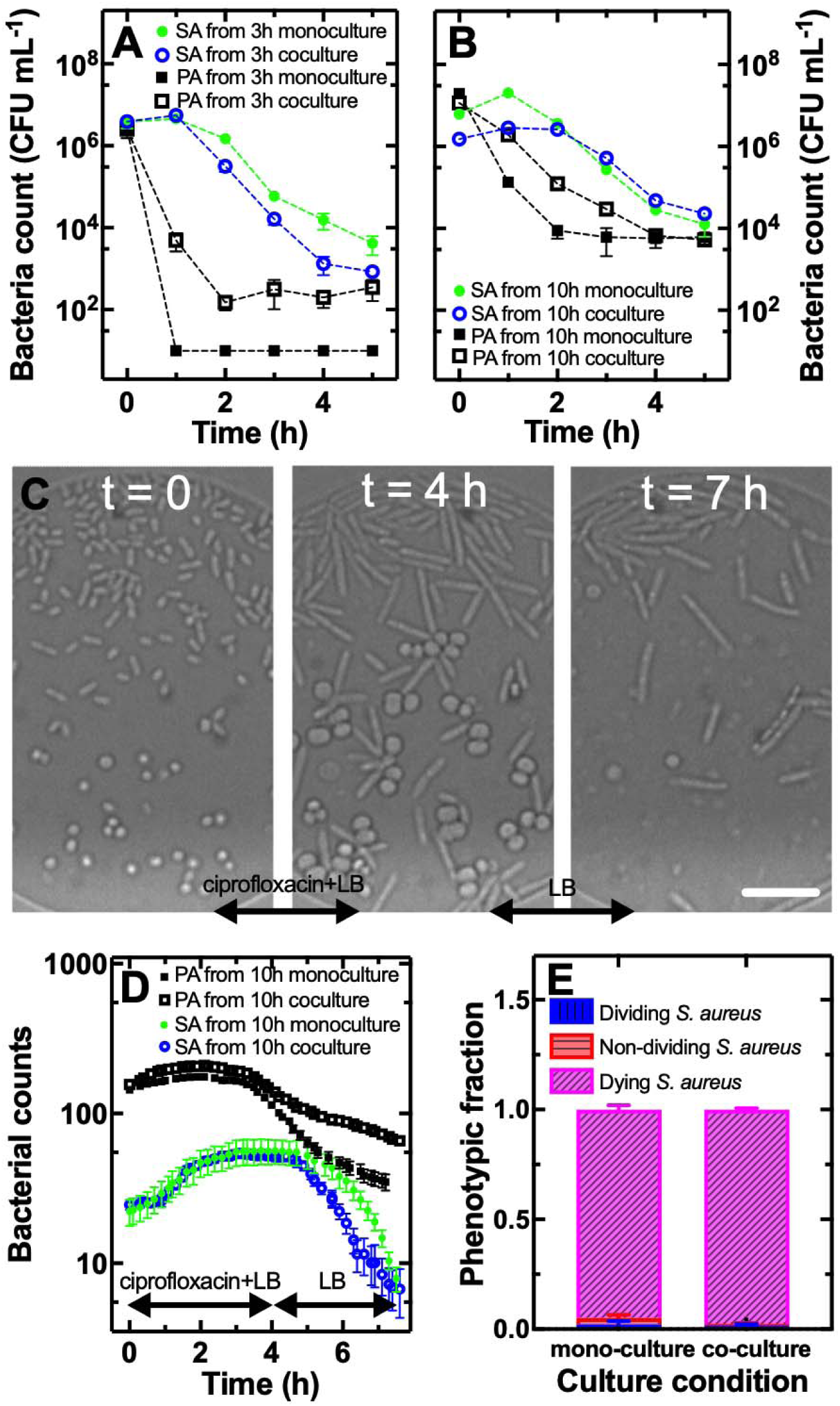
The presence of *P. aeruginosa* does not affect the bactericidal efficacy of ciprofloxacin against *S. aureus* when used at inhibitory concentrations. (a-b) Temporal dependence of bacterial population size for *S. aureus* (filled circles) or *P. aeruginosa* (filled squares) in mono-culture, *S. aureus* (open circles) or *P. aeruginosa* (open squares) in co-culture in well-mixed flasks treated with ciprofloxacin at the MIC value measured against *S. aureus*, i.e. 0.5 µg mL^-1^. Bacteria were subjected to a 5 h ciprofloxacin treatment after being harvested from either (a) 3 h or (b) 10 h of mono-culture or co-culture. Data points are the mean and standard deviation of colony forming unit (CFU) measurements carried out in biological triplicate each consisting of technical duplicate. Numerical values for each replicate are reported in S16 File. (c) Microscopy images of *S. aureus* and *P. aeruginosa* introduced in microfluidic chambers after 10 h of co-culture in flasks and after exposure to ciprofloxacin (0-4 h) and LB (4-7 h) in the microfluidic chambers. Scale bar: 10 µm. (d) Corresponding temporal dependence of bacterial population size for *S. aureus* growing in mono-culture (filled green circles) or in co-culture with *P. aeruginosa* (open blue circles) and for *P. aeruginosa* growing in mono-culture (filled black squares) or co-culture (open black squares). Data points are the mean and standard error of cell count measurements carried out in biological triplicate experiments each consisting of four microfluidic chambers. Numerical values for each replicate are reported in S17 File. (e) Corresponding fraction of dividing (vertically patterned bars), non-dividing (horizontally patterned bars) or lysing (diagonally patterned bars) *S. aureus* in mono-culture or in co-culture with *P. aeruginosa* within microfluidic chambers.

After 3 h or 10 h of co-culture, *S. aureus* displayed similar survival to ciprofloxacin treatment compared to that measured for *S. aureus* after 3 h or 10 h of monoculture when ciprofloxacin was employed at its MIC (Figure 3A and Figure 3B, respectively). In contrast, when ciprofloxacin was employed at 10× or 25× its MIC, *S. aureus* displayed increased survival to ciprofloxacin after both 3 h or 10 h of co-culture compared to the corresponding mono-cultures. Indeed, persisters constituted 0.01% of the *S. aureus* population in 3 h old co-cultures, whereas 3 h old *S. aureus* mono-cultures did not display detectable persisters (Figure S13E and S13G). Furthermore, persisters constituted 1% of *S. aureus* population in 10 h old co-cultures and only 0.01% of *S. aureus* population in 10 h old mono-cultures (Figure S13F and S13H and S16 File).

After 3 h of co-culture, *P. aeruginosa* displayed increased survival to ciprofloxacin treatment compared to that measured for *P. aeruginosa* after 3 h of mono-culture when ciprofloxacin was employed at its MIC (with a 4-log vs a 6-log reduction, Figure S13C). However, we did not observe a difference in terms of ciprofloxacin efficacy against *P. aeruginosa* between 10 h mono- and co-cultures (Figure S13D). Moreover, when ciprofloxacin was employed at 10× or 25× its MIC (against *S. aureus*), *P. aeruginosa* did not display any detectable persisters after 3 h or 10 h of mono- or co-culture (Figure S13E-H and S16 File).

Taken together, these data demonstrate that the presence of a competitor species does not affect the efficacy of ciprofloxacin against *S. aureus* or *P. aeruginosa* when this antibiotic is used at its minimum inhibitory concentration; in contrast, when ciprofloxacin is used above its inhibitory concentration its efficacy against *S. aureus* decreases in the presence of *P. aeruginosa* with the emergence of persisters also in the exponential phase of growth.

Next, we set out to determine whether the decline in the *S. aureus* and *P. aeruginosa* populations during ciprofloxacin treatment was due to cell death or entrance in a viable but non-culturable state. We employed the above described single-cell microfluidics-based platform to image and track the fate of individual *S. aureus* and *P. aeruginosa* cells harvested from 10 h old well-mixed mono-cultures or 10 h old well-mixed co-cultures, while exposing them first to ciprofloxacin dissolved in LB medium for 4 h (at its MIC value against *S. aureus*, i.e. 0.5 µg mL^-1^) and then to LB medium for 3 h within microfluidic chambers (Figure 3C). These experiments were performed in biological triplicates.

1. *S. aureus* harvested from 10 h old well-mixed mono-cultures grew and doubled during the 4 h exposure to ciprofloxacin in all the twelve microfluidic chambers imaged across three biological replicates; during the subsequent exposure to LB medium *S. aureus* continued to double in some chambers, while cells stopped doubling or lysed in other chambers. However, by the end of the experiments, *S. aureus* lysis was observed in all the imaged chambers (Figure S14A and S17 File) which led to a significant decrease in *S. aureus* population (Figure 3D). By the end of the experiments, 5 *S. aureus* cells were in a dividing state, 9 *S. aureus* cells were in a viable but non culturable state, whereas 252 *S. aureus* cells had lysed, i.e. 2%, 3% and 95% of the population, respectively (Figure 3E).
2. *S. aureus* populations harvested from 10 h old well-mixed co-cultures displayed an overall similar response to ciprofloxacin to the one described above for mono-cultures (Figure 3D) in accordance with our time-killing assay data (Figure 3B). However, cell lysis (Figure 3C) and population decline started at an earlier time point in co-culture compared to mono-culture (Figure 3D, Figure S14B and S17 File) and by the end of the experiment fewer *S. aureus* cells were in a dividing or viable but non culturable state (i.e. 1.5% and 0.8%, respectively) compared to mono-culture (Figure 3E).
3. *P. aeruginosa* harvested from 10 h old well-mixed mono-cultures displayed a slower growth compared to *S. aureus* mono-cultures during the 4 h exposure to ciprofloxacin, followed by population decline due to both cell lysis and cell escape from the microfluidic chambers. *P. aeruginosa* harvested from 10 h old well-mixed co-cultures displayed a similar population dynamics, but with a slightly higher surviving fraction (Figure 3D, Figure S14 and S17 File).

Taken together, these data demonstrate that the decline in the *S. aureus* and *P. aeruginosa* populations during ciprofloxacin treatment is primarily due to cell death rather than entrance in a viable but non-culturable state, that ciprofloxacin has a bactericidal effect against both *S. aureus* and *P. aeruginosa* and that the presence of a competitor species does not significantly affect the efficacy of ciprofloxacin against either *S. aureus* or *P. aeruginosa* when this antibiotic is used at its minimum inhibitory concentration.

### Synergistic growth inhibitory effect of vancomycin and *P. aeruginosa* against *S. aureus*

Next, we performed time-killing assays to evaluate the impact of the interactions between *P. aeruginosa* and *S. aureus* on the efficacy of vancomycin, an antibiotic that is only effective against *S. aureus*. We found that both at its MIC value against *S. aureus* (i.e. 1 µg mL^-1^), or at 10× or 25× its MIC, vancomycin had a moderate efficacy when added for 5 h to *S. aureus* mono-cultures in the exponential or stationary phase (i.e. less than 1-log reduction in bacterial counts, Figure 4AB, Figure S15 and S18 File). As expected, *P. aeruginosa* was not affected by vancomycin at any of the antibiotic concentrations employed (Figure S15 and S18 File).

**Figure 4.**
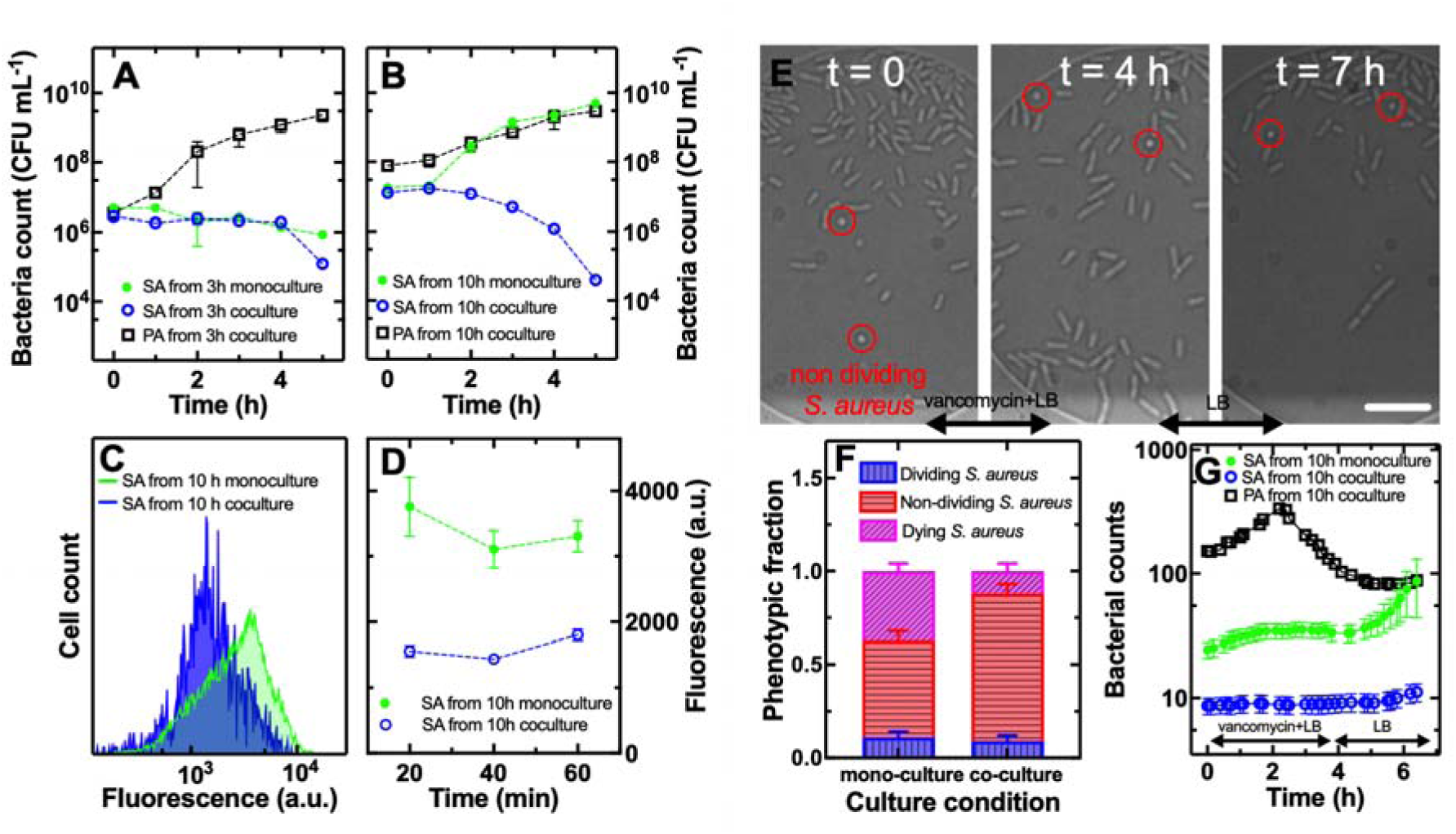
Synergistic growth inhibitory effect of vancomycin and *P. aeruginosa* against *S. aureus*. (a-b) Temporal dependence of bacterial population size for *S. aureus* in mono-culture (filled circles), *S. aureus* (open circles) or *P. aeruginosa* (open squares) in co-culture in well-mixed flasks treated with vancomycin at its MIC value against *S. aureus*, i.e. 1 µg mL^-1^. Bacteria were subjected to a 5 h vancomycin treatment after being harvested from either (a) 3 h or (b) 10 h of mono-culture or co-culture. Data points are the mean and standard deviation of colony forming unit (CFU) measurements carried out in biological triplicate each consisting of technical duplicate. Numerical values for each replicate are reported in S18 File. (c) Distribution of accumulation of vancomycin-NBD in individual *S. aureus* cells harvested from 10 h old mono- (green histogram) or co-culture (blue histogram), exposed to 16 µg mL^-1^ vancomycin-NBD for 1 h and measured via flow cytometry. The data presented are representative of biological triplicate measurements with numerical values reported in Data A in S19 File. (d) Corresponding mean values of vancomycin-NBD fluorescence distributions in *S. aureus* harvested from 10 h old mono- (filled circles) or co-culture (open circles), exposed to vancomycin-NBD for 20 min, 40 min, or 60 min. The data reported are the mean and standard deviation of mean values extracted from fluorescence distributions from biological triplicate each constituted of technical triplicate. (e) Microscopy images of *S. aureus* and *P. aeruginosa* introduced in microfluidic chambers after 10 h of co-culture in flasks and after exposure to vancomycin (0-4 h) and LB (4-7 h) in the microfluidic chambers. Representative non-dividing *S. aureus* are denoted with red circles. Scale bar: 10 µm. (f) Corresponding fraction of dividing (vertically patterned bars) or non-dividing (horizontally patterned bars) *S. aureus* in mono-culture or in co-culture with *P. aeruginosa* within microfluidic chambers. (g) Corresponding temporal dependence of bacterial population size for *S. aureus* in mono-culture (filled green circles) or in co-culture with *P. aeruginosa* (open blue circles) and for *P. aeruginosa* in co-culture (open black squares). Data points are the mean and standard error of cell count measurements carried out in biological triplicate experiments each consisting of four microfluidic chambers. Numerical values for each replicate are reported in S20 File.

After 3 h of co-culture, *S. aureus* displayed similar survival to vancomycin treatment compared to that measured for *S. aureus* after 3 h of mono-culture when vancomycin was employed either at its MIC (Figure 4A), at 10×, or 25× its MIC (Figure S15E and S15G). In striking contrast, after 10 h of co-culture, *S. aureus* displayed a 5-log reduction in survival to vancomycin treatment at its MIC compared to *S. aureus* mono-culture (Figure 4B and S18 File). These data suggest a synergistic bactericidal or growth inhibitory effect between vancomycin, that when used against *S. aureus* mono-cultures had a negligible efficacy (Figure S11D), and *P. aeruginosa* that caused only a 2-log reduction of *S. aureus* counts in co-culture compared to mono-culture in the absence of vancomycin (Figure S15B). This synergistic effect was possibly due to the effect of the *P. aeruginosa* extracellular protease, LasA, that cleaves pentaglycine bridges in *S. aureus* peptidoglycan (*26*) and was strongly upregulated in 10 h old *P. aeruginosa* mono-cultures and co-cultures compared to 3 h old *P. aeruginosa* mono-cultures and co-cultures (8.7 and 8.8 log_2_ fold change, respectively, Figure S10B). Moreover, we recorded a 3-log reduction of *S. aureus* counts for vancomycin treatment at 10×, or 25× its MIC in co-culture compared to that measured for *S. aureus* after 10 h of mono-culture (Figure S15F and S15H).

It is also important to observe the differences in the population dynamics, in response to vancomycin treatment, between *S. aureus* mono-cultures and co-cultures. After 3 h mono-culture, the *S. aureus* population gradually decreased during vancomycin at all concentrations tested. This population dynamics suggests that *S. aureus* is tolerant to vancomycin at the population level, rather than at the sub-population level with the presence of persisters that would instead lead to a bi-phasic killing dynamics (*55*, *56*). Similar population dynamics were measured for *S. aureus* after 3 h of co-culture, except for a sharp decrease in cell counts after 5 h treatment with vancomycin (Figure 4A, S15C, S15E and S15G). These data suggest that after 3 h of co-culture and 5 h of vancomycin treatment in co-culture, a different type of growth inhibition of *S. aureus* cells becomes evident because *P. aeruginosa* starts to exert its negative effect on *S. aureus* in accordance with the data presented in Figure 1.

After 10 h mono-culture, the *S. aureus* population did not decrease during vancomycin treatment neither at 1×, 10×, nor at 25× its MIC indicating complete tolerance to vancomycin at the population level. In contrast, after 10 co-culture, the *S. aureus* population displayed a bi-phasic dynamics with a gradual decrease during the first 3 h of vancomycin treatment followed by a sharp decrease in cell counts (Figure 4B, S15D, S15F and S15H), i.e. the opposite trend observed when a bacterial population harbours persisters (*55*, *56*).

Taken together, these data suggest that the presence of *P. aeruginosa* enhances either killing or growth inhibition of *S. aureus* via vancomycin that has an otherwise negligible efficacy against stationary phase *S. aureus*.

### Vancomycin accumulation in *S. aureus* is lower in the presence of *P. aeruginosa*

Next, we set out to test the hypothesis that the accumulation of vancomycin in *S. aureus* could be altered by the continued presence of *P. aeruginosa*. In fact, we observed an increase in PG lipids in co-culture with respect to *S. aureus* mono-culture (Table 1). PG lipids are negatively charged and could therefore bind and sequester positively charged vancomycin molecules(*57*) hindering their translocation through the membrane and their binding to the D-alanyl-D-alanine (D-Ala-D-Ala) peptide motif of the peptidoglycan precursor (*58*).

In order to test this hypothesis, we used flow cytometry and a fluorescent derivative of vancomycin, vancomycin-nitrobenzofuran (NBD) (*47*, *59*). Upon incubation in vancomycin-NBD at a concentration of 16 µg mL^-1^ (i.e. 8× the MIC value of vancomycin-NBD against *S. aureus*) in M9 minimal medium for 1 h, we found similar distributions of vancomycin-NBD single-cell fluorescence for 10 h old mono-cultures of *S. aureus* ATCC 25923 and *S. aureus* SH1000 mCherry; vancomycin-NBD did not accumulate instead in 10 h old mono-cultures of *P. aeruginosa* PA14 (Figure S16A and Data A in S19 File). As expected, *S. aureus* SH1000 mCherry displayed ECD fluorescence levels distinguishable from M9 minimal medium signal, whereas neither *S. aureus* ATCC25923 nor *P. aeruginosa* PA14 displayed ECD fluorescence levels distinguishable from M9 minimal medium (Figure S16B). Next, we incubated 10 h old co-cultures of *S. aureus* SH1000 mCherry and *P. aeruginosa* PA14 in vancomycin-NBD at a concentration of 16 µg mL^-1^ in M9 minimal medium. By simultaneously using the FITC and ECD flow cytometer detectors, we gated only cells that expressed mCherry at levels distinguishable from M9 minimal medium, i.e. *S. aureus* SH1000 mCherry cells (Figure 4C and S16). We compared the distributions of vancomycin-NBD single-cell fluorescence for gated *S. aureus* SH1000 mCherry cells from 10 h old co-cultures and 10 h old mono-cultures after 20 min, 40 min and 60 min incubation in vancomycin-NBD at a concentration of 16 µg mL^-1^ in M9 minimal medium. According to our hypothesis above, we found that *S. aureus* SH1000 mCherry cells from 10 h old co-cultures accumulated significantly lower levels of vancomycin-NBD compared to *S. aureus* SH1000 mCherry cells from 10 h old co-cultures (Figure 4C, Figure 4D, Figure S17 and S19 File).

Moreover, using ultra-high performance liquid chromatography-quadrupole time-of-flight mass spectrometry we found a further increase in *S. aureus* PG lipid expression in both mono-cultures and co-cultures treated with vancomycin (Table 2). These data suggest that the upregulation of charged PG lipids is a strategy employed by *S. aureus* to counteract exposure to either *P. aeruginosa* or vancomycin, indirectly suggesting that both stressors elicit a similar response in *S. aureus* in terms of membrane lipid composition.

**Table 2.**
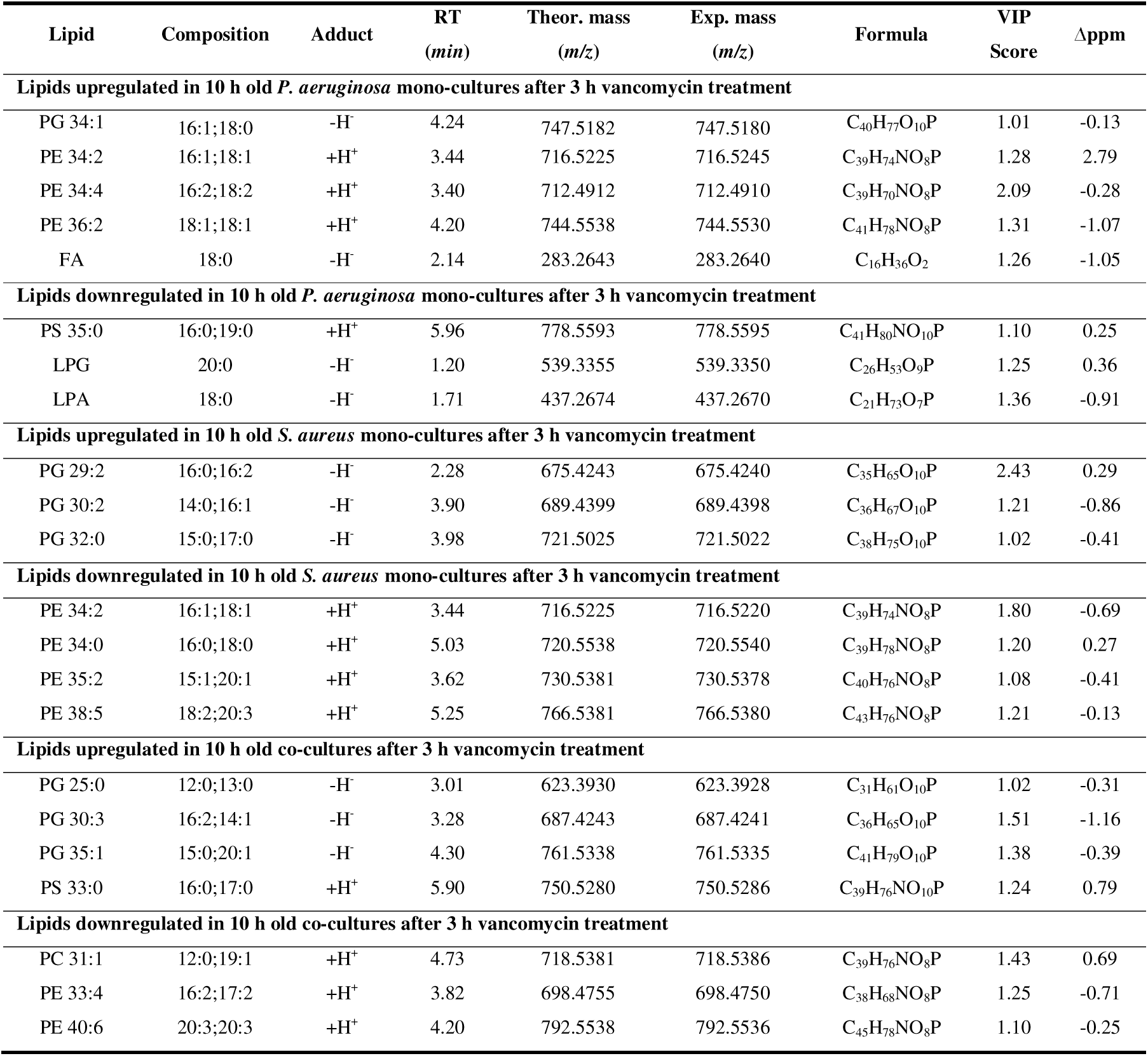
Lipids differentially regulated after vancomycin treatment. . Lipid class, fatty acid composition, adduct type, retention time (RT), theoretical and experimental mass-to-charge ratio (m/z), molecular formula, variable importance in projection (VIP) score, and mass error (Δppm) measured via quantitative time-of-flight liquid chromatography mass spectrometry.

| Lipid | Composition | Adduct | RT<br>(min) | Theor. mass<br>(m/z) | Exp. mass<br>(m/z) | Formula | VIP<br>Score | Δppm |
| --- | --- | --- | --- | --- | --- | --- | --- | --- |
| **Lipids upregulated in 10 h old *P. aeruginosa* mono-cultures after 3 h vancomycin treatment** |  |  |  |  |  |  |  |  |
| PG 34:1 | 16:1;18:0 | -H<sup>+</sup> | 4.24 | 747.5182 | 747.5180 | C<sub>40</sub>H<sub>77</sub>O<sub>10</sub>P | 1.01 | -0.13 |
| PE 34:2 | 16:1;18:1 | +H<sup>+</sup> | 3.44 | 716.5225 | 716.5245 | C<sub>39</sub>H<sub>74</sub>NO<sub>8</sub>P | 1.28 | 2.79 |
| PE 34:4 | 16:2;18:2 | +H<sup>+</sup> | 3.40 | 712.4912 | 712.4910 | C<sub>39</sub>H<sub>70</sub>NO<sub>8</sub>P | 2.09 | -0.28 |
| PE 36:2 | 18:1;18:1 | +H<sup>+</sup> | 4.20 | 744.5538 | 744.5530 | C<sub>41</sub>H<sub>78</sub>NO<sub>8</sub>P | 1.31 | -1.07 |
| FA | 18:0 | -H<sup>+</sup> | 2.14 | 283.2643 | 283.2640 | C<sub>16</sub>H<sub>36</sub>O<sub>2</sub> | 1.26 | -1.05 |
| **Lipids downregulated in 10 h old *P. aeruginosa* mono-cultures after 3 h vancomycin treatment** |  |  |  |  |  |  |  |  |
| PS 35:0 | 16:0;19:0 | +H<sup>+</sup> | 5.96 | 778.5593 | 778.5595 | C<sub>41</sub>H<sub>80</sub>NO<sub>10</sub>P | 1.10 | 0.25 |
| LPG | 20:0 | -H<sup>+</sup> | 1.20 | 539.3355 | 539.3350 | C<sub>26</sub>H<sub>53</sub>O<sub>9</sub>P | 1.25 | 0.36 |
| LPA | 18:0 | -H<sup>+</sup> | 1.71 | 437.2674 | 437.2670 | C<sub>21</sub>H<sub>73</sub>O<sub>7</sub>P | 1.36 | -0.91 |
| **Lipids upregulated in 10 h old *S. aureus* mono-cultures after 3 h vancomycin treatment** |  |  |  |  |  |  |  |  |
| PG 29:2 | 16:0;16:2 | -H<sup>+</sup> | 2.28 | 675.4243 | 675.4240 | C<sub>35</sub>H<sub>65</sub>O<sub>10</sub>P | 2.43 | 0.29 |
| PG 30:2 | 14:0;16:1 | -H<sup>+</sup> | 3.90 | 689.4399 | 689.4398 | C<sub>36</sub>H<sub>67</sub>O<sub>10</sub>P | 1.21 | -0.86 |
| PG 32:0 | 15:0;17:0 | -H<sup>+</sup> | 3.98 | 721.5025 | 721.5022 | C<sub>38</sub>H<sub>75</sub>O<sub>10</sub>P | 1.02 | -0.41 |
| **Lipids downregulated in 10 h old *S. aureus* mono-cultures after 3 h vancomycin treatment** |  |  |  |  |  |  |  |  |
| PE 34:2 | 16:1;18:1 | +H<sup>+</sup> | 3.44 | 716.5225 | 716.5220 | C<sub>39</sub>H<sub>74</sub>NO<sub>8</sub>P | 1.80 | -0.69 |
| PE 34:0 | 16:0;18:0 | +H<sup>+</sup> | 5.03 | 720.5538 | 720.5540 | C<sub>39</sub>H<sub>78</sub>NO<sub>8</sub>P | 1.20 | 0.27 |
| PE 35:2 | 15:1;20:1 | +H<sup>+</sup> | 3.62 | 730.5381 | 730.5378 | C<sub>40</sub>H<sub>76</sub>NO<sub>8</sub>P | 1.08 | -0.41 |
| PE 38:5 | 18:2;20:3 | +H<sup>+</sup> | 5.25 | 766.5381 | 766.5380 | C<sub>43</sub>H<sub>76</sub>NO<sub>8</sub>P | 1.21 | -0.13 |
| **Lipids upregulated in 10 h old co-cultures after 3 h vancomycin treatment** |  |  |  |  |  |  |  |  |
| PG 25:0 | 12:0;13:0 | -H<sup>+</sup> | 3.01 | 623.3930 | 623.3928 | C<sub>31</sub>H<sub>61</sub>O<sub>10</sub>P | 1.02 | -0.31 |
| PG 30:3 | 16:2;14:1 | -H<sup>+</sup> | 3.28 | 687.4243 | 687.4241 | C<sub>36</sub>H<sub>65</sub>O<sub>10</sub>P | 1.51 | -1.16 |
| PG 35:1 | 15:0;20:1 | -H<sup>+</sup> | 4.30 | 761.5338 | 761.5335 | C<sub>41</sub>H<sub>79</sub>O<sub>10</sub>P | 1.38 | -0.39 |
| PS 33:0 | 16:0;17:0 | +H<sup>+</sup> | 5.90 | 750.5280 | 750.5286 | C<sub>39</sub>H<sub>76</sub>NO<sub>10</sub>P | 1.24 | 0.79 |
| **Lipids downregulated in 10 h old co-cultures after 3 h vancomycin treatment** |  |  |  |  |  |  |  |  |
| PC 31:1 | 12:0;19:1 | +H<sup>+</sup> | 4.73 | 718.5381 | 718.5386 | C<sub>39</sub>H<sub>76</sub>NO<sub>8</sub>P | 1.43 | 0.69 |
| PE 33:4 | 16:2;17:2 | +H<sup>+</sup> | 3.82 | 698.4755 | 698.4750 | C<sub>38</sub>H<sub>68</sub>NO<sub>8</sub>P | 1.25 | -0.71 |
| PE 40:6 | 20:3;20:3 | +H<sup>+</sup> | 4.20 | 792.5538 | 792.5536 | C<sub>45</sub>H<sub>78</sub>NO<sub>8</sub>P | 1.10 | -0.25 |

Taken together, these data demonstrate that the synergistic killing or growth inhibitory effect of vancomycin and *P. aeruginosa* is not due to increased accumulation of vancomycin in *S. aureus* in the presence of *P. aeruginosa*.

### The combined effect of vancomycin and *P. aeruginosa* induces *S. aureus* in a viable but non culturable state

Next, we set out to determine whether the decline in the *S. aureus* population measured in the presence of *P. aeruginosa* and vancomycin is due to killing of *S. aureus* cells or *S. aureus* cells entering in a viable but non-culturable state. As above, we employed our single-cell microfluidics-based platform to image and track the fate of individual *S. aureus* harvested from 10 h old well-mixed mono-cultures or individual *S. aureus* and *P. aeruginosa* cells harvested from 10 h old well-mixed co-cultures. We exposed these bacteria, while hosted within microfluidic chambers, first to vancomycin dissolved in LB medium at its MIC value against *S. aureus*, i.e. 1 µg mL^-1^, for 4 h and then to LB medium for 3 h.

The growth of *S. aureus* harvested from 10 h old well-mixed mono-cultures was highly heterogeneous: in two of the twelve microfluidic chambers imaged across three biological replicates, we recorded sustained bacterial doubling both during exposure to vancomycin and during exposure to LB medium; in three chambers we recorded sustained doubling only during exposure to LB medium; in three chambers we recorded only sporadic bacterial doubling or no doubling at all; in four chambers we recorded sporadic growth during exposure to vancomycin followed by cell lysis during exposure to LB medium or cells stained by PI at the end of the experiment (Figure S18A and Data A in S20 File). Overall, 11% of the cells within the initial *S. aureus* population were growing and dividing by the end of the experiment, 52% of the *S. aureus* cells were in a motile but non-dividing state and 37% of the *S. aureus* cells either lysed or stained with PI at the end of the experiments (Figure 4F and Data B in S20 File). As a result, the *S. aureus* population dynamics was characterised by very slow and sporadic growth during exposure to vancomycin from 24 ± 13 bacteria per chamber at t = 0 to 33 ± 17 bacteria per chamber at t = 4 h (Figure 4G and Data A S20 File) with an average lag time of 44 min followed by an average doubling time of 180 min (Figure S19). During the subsequent exposure to LB, the *S. aureus* population dynamics was characterised by faster but heterogeneous growth up to 87 ± 149 bacteria per chamber at t = 7 h (Figure 4G and Data A S20 File) and an average doubling time of 32 min (Figure S19). This population dynamics is in accordance with the one we recorded via bulk CFU assays (Figure 4B) that did not allow us, however, to capture the non-dividing *S. aureus* phenotype and the large heterogeneity in growth during and after treatment with vancomycin.

*S. aureus* harvested from 10 h old well-mixed co-cultures did not display any bacterial doubling across the twelve imaged microfluidic chambers during exposure to vancomycin; sustained doubling during the following exposure to LB medium was recorded only in three chambers, whereas the dominant phenotype observed in the remaining chambers was a motile but non-dividing state of *S. aureus* that did not stain with PI (Figure S18 B and Data C in S20 File). Overall, 9% of the cells within the initial *S. aureus* population were growing and dividing by the end of the experiment, 79% of the *S. aureus* cells were in a motile but non-dividing state and 12% of the *S. aureus* cells either lysed or stained with PI at the end of the experiments (Figure 4F and Data B in S20 File). As a result, *S. aureus* population dynamics was characterised by no growth during exposure to vancomycin (Figure 4G and Data C in S20 File) with an average lag time of around 6 h (hence regrowth started only after vancomycin had been removed, Figure S19). During the subsequent exposure to LB, the *S. aureus* population dynamics was characterised by limited growth with an overall increase from 9 ± 5 bacteria per chamber at t = 0 to 12 ± 8 bacteria per chamber at t = 7 h (Figure 4G and Data C in S20 File) and an average doubling time of 34 min (Figure S19). This population dynamics is different with respect to the one we recorded via bulk CFU assays showing that in the presence of *P. aeruginosa* a 4 h treatment with vancomycin at its MIC value against *S. aureus*, causes a 1-log decline in the *S. aureus* population (Figure 4B). It is conceivable that the decline observed in our bulk assays is not due cell death but to a proportion of the population entering a viable but non-culturable state that we can capture via microfluidics-microscopy but that is undetectable via CFU assays. However, it is also conceivable that these differences are due to decreased concentrations of LasA, or other molecules secreted by *P. aeruginosa*, in the open microfluidic chambers (where fresh nutrients are constantly supplied, secreted bacterial molecules are washed away and *P. aeruginosa* can escape, see below) compared to the closed well-mixed flasks (where secreted bacterial molecules and *P. aeruginosa* cannot escape the flasks).

During exposure to vancomycin, *P. aeruginosa* harvested from 10 h old well-mixed co-cultures displayed growth comparable to that recorded in the absence of vancomycin (Figure 4G and Figure 1G, respectively). *P. aeruginosa* population expanded during the first 2 h of exposure to vancomycin and decreased from the 2 h time point onwards in all imaged chambers (Figure S18C and Data E in S20 File) due to increased cellular motility, possibly via type IV pili(*43*), and escape out of the chambers. As a consequence, the ratio of *P. aeruginosa* cell number over *S. aureus* cell number per microfluidic chamber was 27 ± 23 at t = 0, it increased to a maximum of 58 ± 49 after 2 h and then decreased down to a minimum of 15 ± 15 at t = 7 h (Figure S18D).

## Discussion

### *S. aureus* population decline in the presence of *P. aeruginosa* is conserved across a diverse set of *S. aureus* and *P. aeruginosa* strains

*S. aureus* population initial expansion followed by a steady decline in the continued presence of *P. aeruginosa* had been previously reported. Specifically, planktonic and biofilm *S. aureus* 8325-4 displayed a population decline after 6 h or 14 h co-culture with *P. aeruginosa* PA14 on monolayers of cystic fibrosis bronchial epithelial airway cells or plastic, respectively (*25*). Furthermore, planktonic *S. aureus* Newman displayed a population decline after 4 h co-culture with *P. aeruginosa* PA14 in microtiter plates (*46*). *S. aureus* JE2 also displayed a population decline after 12 h co-culture with *P. aeruginosa* PAO1 on agar plates (*28*), whereas *S. aureus* SA2597 displayed a population decline after 4 h co-culture with *P. aeruginosa* PA2596 on agar plates (*29*). We further demonstrate the generality of these population dynamics by showing that planktonic *S. aureus* ATCC 25923, RYC157 and RYC165 displayed a population decline after 9 or 10 h co-culture with *P. aeruginosa* PA14. However, *S. aureus* RYC157 displayed a population decline only after 20 h of co-culture with *P. aeruginosa* RYC157, in accordance with previous data suggesting that *P. aeruginosa* isolates from co-infected CF patients are less antagonistic towards *S. aureus* than isolates from mono-infected CF patients (*23*).

The dynamics of these microbial interactions have been generally rationalised using Lotka-Volterra pairwise models which are based on the fitness effect of such interactions (*60*, *61*). However, these pairwise models often perform less well than mechanistic models in terms of capturing diverse microbial interactions (*62*). We introduce a simplified mechanistic model that allows us to capture the population dynamics of *S. aureus* and *P. aeruginosa* both in mono- and co-culture and to output parameter estimates that might allow, in future studies, to explain the time difference in the onset of *S. aureus* decline across different strains, media and environmental setups.

### *S. aureus* enters a viable but non-culturable state in the presence of *P. aeruginosa*

The observed *S. aureus* population decline could be due either to killing of *S. aureus* by *P. aeruginosa* or *S. aureus* entering a viable but non-culturable state or a persister state with very long lag time (*63–65*) and therefore becoming undetectable via colony forming unit assays. The viable but non-culturable state is a ubiquitous survival strategy adopted by many bacteria (*66*, *67*), including *S. aureus* (*68–70*), in response to adverse environmental conditions such as starvation (*39*, *71*), cold stress (*72*), host-induced oxidative stress and ATP depletion (*73*), or antibiotic treatment (*69*, *74*). Intriguingly, *S. aureus* in a viable but non-culturable state has been recently detected via qPCR in the sputum of cystic fibrosis patients concurrently with poor lung functionality (*75*) and in biofilm samples from central venous catheters (*76*). Furthermore, recent data suggested that populations of *S. aureus* ATCC 25923 and of *S. aureus* clinical isolates entered a viable but non-culturable state after 48 h co-culture in dual-species biofilms with *P. aeruginosa* PA14 (*77*).

Here we show that a subpopulation of *S. aureus* enters a viable but non-culturable state already after 10 h of planktonic co-culture with *P. aeruginosa* and that individual viable but non-culturable *S. aureus* cells remain in a motile non-dividing state for at least a further 9 h, whereas other clonal *S. aureus* cells resume growth. Consistent with these data we also observed that *S. aureus* cells that resumed growth displayed a longer lag time compared to *S. aureus* cells harvested from mono-culture and displayed an overall more heterogeneous growth. These findings are consistent with previously observed heterogeneity in the activation of *S. aureus* stationary markers and persister formation (*78*) and previous data suggesting that several *S. aureus* isolates are growth inhibited but not killed by *P. aeruginosa* (*28*). Our single-cell microscopy data suggest that *S. aureus* cell death and lysis plays a limited role in the *S. aureus* population decline observed in the presence of *P. aeruginosa* in our study and possibly also in previous studies using bulk microbiological assays (*25*, *28*, *46*). The observed population decline in the presence of *P. aeruginosa* is instead primarily due to *S. aureus* entering a viable but non-culturable state leading to a possible underestimation or non-detection in clinical samples which could constitute a serious risk to human health. Future studies should investigate whether the presence of *P. aeruginosa* cells is necessary to induce *S. aureus* in a viable but non-culturable state or whether *P. aeruginosa* supernatants are sufficient for this, considering that a previous study at the population-level showed that *P. aeruginosa* supernatants have an impact on the growth of *S. aureus* (*31*).

### Similarities between *S. aureus* viable but non-culturable cells in the presence of *P. aeruginosa* and *S. aureus* intracellular persisters

Consistent with our data showing that *S. aureus* enters a viable but non-culturable state, our RNA-sequencing analysis demonstrates that in the presence of *P. aeruginosa*, *S. aureus* drastically rewires its transcriptome and downregulates key metabolic processes including nucleoside metabolism, nitrogen metabolism and transport, amino acid biosynthesis, biogenesis and cellular organization, while upregulating cellular catabolism and protein maturation. These molecular processes were reprogrammed in response to an increase in molecular transport, particularly iron, DNA-templated transcription initiation, protein repair and carbohydrate biosynthesis in *P. aeruginosa*, with a downregulation of carbon and nitrogen metabolism.

The observed downregulation of nitrogen metabolism in both *S. aureus* and *P. aeruginosa* is consistent with previous transcriptomic data highlighting nitrogen starvation as a co-culture specific response in both *P. aeruginosa* and *S. aureus* with the upregulation of the master nitrogen regulator *ntrC* and the downregulation of *gltB* and *gltD*, coding for glutamine to glutamate conversion (*46*). Dysregulation of genes involved in major metabolism pathways of carbohydrates and amino acids, such as *acsA* which encodes the acetyl-coenzyme A synthetase, was also reported for *S. aureus* SA2597 co-cultured with *P. aeruginosa* PA2596 (*29*).

We found some important similarities between the transcriptomic profile of *S. aureus* in co-culture with *P. aeruginosa* and *S. aureus* intracellular persisters treated with oxacillin within macrophages. Intracellular persisters down regulated the biosynthesis of key amino acids, such as valine, leucine, isoleucine, lysine and tryptophan (*53*). Accordingly, we found down regulation of genes encoding lysine biosynthesis, *dapA-D*, glutamine biosynthesis, *glnA*, valine and isoleucine biosynthesis, *ilvD*, in *S. aureus* co-cultured with *P. aeruginosa*. Moreover, intracellular persisters induced the heat shock stimulon, including genes *dnaK*, *grpE*, *clpB*, *groEL* and *groES* (*53*) that participate in the degradation of defective proteins (*79*). Accordingly, we found upregulation of these and other protein maturation genes in *S. aureus* co-cultured with *P. aeruginosa* but we did not find evidence of dark foci induced by protein aggregation (*42*, *52*, *73*, *80*).

Intracellular *S. aureus* persisters also displayed ATP levels comparable to control *S. aureus* cells (*53*). Similarly, we did not find evidence of strong downregulation of genes involved in energetic metabolism and redox reactions (e.g. the L-lactate dehydrogenase or the NADP-dependent isocitrate dehydrogenase) in *S. aureus* in co-culture with *P. aeruginosa*, in accordance with a previous study (*29*). These data are instead different from previous findings that *P. aeruginosa* inhibits *S. aureus* respiration inducing a low ATP state (*6*) and that persister formation in *S. aureus* is associated with ATP depletion (*78*). We also found upregulation of the *codY* regulon that is de-repressed under amino acid starvation (*81*) and upon *S. aureus* uptake in macrophages, playing a role in intracellular persistence through ppGpp regulation (*53*).

Intracellular *S. aureus* persisters also displayed dysregulated but active protein synthesis: aminoacyl-tRNA synthetase encoding genes were downregulated, whereas members of the protein synthesis machinery and ribosome recycling were upregulated(*53*). Similarly, we found a downregulation of aminoacyl-tRNA synthetase encoding genes such as *thiL*, *mnmA* or *trnB* in co-culture with *P. aeruginosa* and a slight upregulation of elongation factors *infAB* and ribosome recycling gene *frr*. Moreover, we observed reduced but active expression of mCherry in viable but non-culturable *S. aureus* in co-culture with *P. aeruginosa*, similarly to reduced but active expression of GFP in intracellular persisters (*53*). Taken together these data suggest that viable but non-culturable *S. aureus* in co-culture with *P. aeruginosa* adopt a survival lifestyle that resembles the one adopted by intracellular *S. aureus* persisters (*53*).

Entrance in a viable but non-culturable state could also be partly due to inhibition of the electron transport chain of *S. aureus* via the downregulation of *qoxA-D* encoding the cytochrome aa3 quinol oxidase and the upregulation of the *narK1* antiporter gene, in accordance with a previous study on the early responses between *P. aeruginosa* PA14 and *S. aureus* Newman (*46*) and a previous study on *S. aureus* intracellular persisters(*53*). However, we also found upregulation of cytochrome C oxidase genes *coxAB*, in contrast with a previous study (*46*). We also found increased expression of the lactose operon *lacA-G* in *S. aureus* co-cultured with *P. aeruginosa*, similarly to what observed for intracellular *S. aureus* persisters (*53*); however, we did not find a significant downregulation of glucose metabolism enzyme encoding genes (a part from *gap* that was strongly downregulated), whereas previous work suggested that a carbon source shift between glucose and lactose is a trigger factor for intracellular persistence (*82*). Moreover, we did not find strong evidence of increased expression of genes encoding enzymes for fermentation in *S. aureus* such as *pflAB*, *ldh*, *adhEP*, *gap*, *tpiA*. In contrast, two previous studies that investigated genome-wide gene expression profile changes in *S. aureus* in coculture for 6 h or 3 h with *P. aeruginosa* found increased expression of genes encoding enzymes for fermentation in *S. aureus*(*25*, *46*). Similarly, L-lactate dehydrogenase *lldA* was the most upregulated gene of *P. aeruginosa* in response to *S. aureus* Newman lactate secretion (*46*), whereas according to our transcriptomic data it was only slightly upregulated after both 3 h and 10 h co-culture, consistent with reduced lactate secretion from *S. aureus*. Therefore, our transcriptomic data do not fully support that *S. aureus* entrance in a viable but non-culturable state in the presence of *P. aeruginosa* is linked to inhibition of the electron transport chain of *S. aureus* or a switch from glucose to lactose metabolism or a switch from respiration to fermentation in *S. aureus*.

Finally, our data suggest that *P. aeruginosa*-produced HQNO, rhamnolipids and siderophores might not underlie the *S. aureus* population decline that we observe in our culture conditions. In fact, coculture with Δ*pvdA*Δ*pchE* or Δ*pqsA P. aeruginosa* strains (that lack synthesis of pyoverdine and pyochelin or the production of HQNO, respectively) markedly reduced *P. aeruginosa*-mediated killing of *S. aureus* 8325-4 biofilms (*25*). In contrast, we found that coculture with the Δ*pvdA*Δ*pchEF* strain did not reduce *S. aureus* population decline. Moreover, we found that the pyoverdine, rhamnolipid and HQNO pathways were upregulated to similar levels after 10 h of mono- or coculture. In contrast, the pyochelin pathway, DNA-templated transcription initiation, including the sigma factors *pvdS* and *rpoD*, and dicarboxylic acid metabolism processes were strongly upregulated in co-culture with *S. aureus* but strongly downregulated in mono-culture, thus suggesting that the continued presence of *S. aureus* induces these processes in *P. aeruginosa*. Our data also suggest downregulation of genes involved in intracellular accumulation of purine in *S. aureus*, such as the uracil transporter PyrP, the pyrimidine *de novo* synthesis operon *pyrPEBCAA*-*carB*-*pyrF*, and the purine *de novo* synthesis gene *purA*. In contrast, a previous study suggested that these genes were among the most highly induced in *S. aureus* in co-culture compared to mono-culture (*46*).

Taken together these data demonstrate that multiple pathways contribute to a different extent to the generation and maintenance of *S. aureus* viable but non-culturable cells in the presence of *P. aeruginosa* and that *S. aureus* adopts a survival lifestyle that resembles the one adopted by intracellular *S. aureus* persisters (*53*). However, we acknowledge that our transcriptomic data are acquired from *S. aureus* populations that contain a mixture of viable but non-culturable cells and growing cells and that our data do not take into account post-translational modifications that represent a second control point to survive external stressors.

### Differential efflux regulation contributes to ciprofloxacin decreased efficacy against *S. aureus* in the presence of *P. aeruginosa*

Ciprofloxacin treatment was less effective against stationary phase compared to exponential phase *S. aureus* or *P. aeruginosa* mono-cultures in line with previous findings and possibly due to a larger subset of bacteria being in a non-replicating and low metabolically active state in the stationary phase of growth (*49*, *78*, *83–86*). In line with this hypothesis, ciprofloxacin treatment became even less effective against *S. aureus* in co-culture with *P. aeruginosa* compared to *S. aureus* mono-culture, due to a subpopulation of *S. aureus* entering a viable but non-culturable state. Reduced ciprofloxacin efficacy against *S. aureus* while in co-culture with *P. aeruginosa* is in accordance with a previous study reporting that the supernatants of several *P. aeruginosa* isolates were antagonistic towards ciprofloxacin killing (*6*). Increased *S. aureus* survival to ciprofloxacin was due to the downregulation of the cellular targets of ciprofloxacin, i.e. the DNA gyrase GyrAB and the topoisomerase IV ParCE; the downregulation of genes that were previously found to be downregulated in *S. aureus* persisters hosted by high-oxidative-stress cells (*73*) or induced by reactive oxygen species (*87*), i.e. *rplA*, *rplM*, *atpA*, *prfA* and *sdhAB*; the upregulation of the NorA, NorB LmrS, SepA, Tet(38) and SbnD efflux pumps which are known to extrude ciprofloxacin and other quinolones (*88*) in *S. aureus*, while in co-culture with *P. aeruginosa* compared to mono-culture as recorded in our transcriptomic data. These data were also consistent with previous findings that while in co-culture with *P. aeruginosa*, *S. aureus* upregulates efflux pumps belonging to the Nor family leading to an increase in resistance of *S. aureus* to tetracycline and ciprofloxacin (*29*). Alternatively, HQNO secreted by *P. aeruginosa* could inhibit *S. aureus* respiration inducing a low ATP, multidrug-tolerant state in *S. aureus* (*6*). However, we observed limited induction of the HQNO pathway in *P. aeruginosa* both in mono- and co-cultures and we did not observe HQNO-induced inhibition of *S. aureus* respiration (*6*).

Finally, our single-cell analysis revealed that both *S. aureus* and *P. aeruginosa* grew and doubled during exposure to ciprofloxacin and started to lyse only during the subsequent exposure to LB medium, leading to a significant decrease in both bacterial populations which was primarily due to cell death rather than entrance in a viable but non-culturable state.

### Vancomycin reduced accumulation in viable but non culturable *S. aureus* in the presence of *P. aeruginosa*

Vancomycin had a moderate efficacy against exponential phase *S. aureus* and was not effective against stationary phase *S. aureus* in accordance with previous clinical data(*89*). In contrast, *S. aureus* displayed a 1-log or a 5-log reduction in survival to vancomycin treatment at its MIC after co-culture with *P. aeruginosa* PA14 for 3 h or 10 h, respectively, according to our time-killing assays. These data are in agreement with a previous study that investigated vancomycin efficacy against exponential phase *S. aureus* HG003 exposed to the supernatants of several *P. aeruginosa* strains and found a 2-log reduction in survival to vancomycin treatment after exposure to *P. aeruginosa* PA14 supernatant (*6*). This study also found that vancomycin killing of *S. aureus* was mediated by *P. aeruginosa* extracellular lytic enzyme, LasA, that cleaves pentaglycine cross bridges in *S. aureus* peptidoglycan and has been shown to attack the cell wall of *S. aureus* during *in vivo* competition (*90*). In line with these data we found that *lasAB* were strongly upregulated at t = 10 h compared to t = 3 h both in mono- and co-culture. Our data therefore further corroborate the hypothesis that LasA mediates population decline of *S. aureus* in co-culture with *P. aeruginosa* and demonstrate that LasA secretion is not induced by the presence of *S. aureus*. Our single-cell analyses, however, favour the hypothesis that the observed population decline of *S. aureus* under vancomycin treatment in co-culture is due to *S. aureus* entering a viable but non-culturable state rather than being killed, since we observed limited *S. aureus* cell death in co-culture in the presence of vancomycin within the timeframe of our experiments.

We also found that vancomycin accumulation in *S. aureus* is reduced in co-culture compared to mono-culture, which is consistent with *S. aureus* entrance in a viable but non-culturable state, with a possible downregulation of cell wall turnover which is targeted by vancomycin. Accordingly, our transcriptomic analysis revealed upregulation of cell wall induction stimulon in co-culture compared to mono-culture, including the two-component regulatory system VraS/VraR involved in the control of the cell wall peptidoglycan biosynthesis (*91*), as previously observed for intracellular persisters (*53*). We also measured downregulation of penicillin-binding protein 4 (Pbp4) that has transpeptidase and carboxypeptidase activities and catalyzes the final step in peptidoglycan synthesis (*92*). Moreover, our lipidomic analysis revealed changes in the *S. aureus* membrane, with an increase in PG lipids and a decrease in PC lipids, in co-culture compared to mono-culture. PG lipids were further upregulated in co-cultures treated with vancomycin, sequestering positively charged vancomycin molecules (*57*) and hindering their translocation through the membrane and their binding to the D-alanyl-D-alanine peptide motif of the peptidoglycan precursor (*58*). These alterations in membrane lipid composition may play a critical role in modulating membrane rigidity and permeability, ultimately shaping *S. aureus* response to antibiotic treatment. Additionally, HQNO, pyoverdine and pyochelin secreted by *P. aeruginosa* could also decrease vancomycin accumulation in *S. aureus* by slowing down cellular growth. However, we observed limited induction of the HQNO pathway in *P. aeruginosa* both in mono- and co-cultures and we observed neither HQNO-induced *S. aureus* shift to fermentative growth (*31*) nor HQNO-induced inhibition of *S. aureus* respiration (*6*).

Our findings demonstrate that *S. aureus* population decline in the presence of *P. aeruginosa* is primarily due to a subpopulation of *S. aureus* entering in a viable but non-culturable state that cannot be detected via traditional microbiology and antibiotic susceptibility assays. Failure to detect these subpopulations could lead to an underestimation or non-detection of *S. aureus* in clinical samples, which would constitute a serious risk to human health, particularly in chronic infections, since entrance in a viable but non culturable state is the primary survival strategy of *S. aureus* in response to vancomycin treatment.

## Materials and Methods

### Chemicals and bacterial strains

All chemicals were purchased from Merck unless otherwise stated. LB medium (10 g L^-1^ tryptone, 5 g L^-1^ yeast extract and 10 g L^-1^ NaCl) and LB agar plates (LB with 15 g L^-1^ agar) were used for planktonic growth and streak plates as previously described (*93*). Selective media for *P. aeruginosa* was Pseudomonas isolation agar (PIA; Merck product number: 17208), while selective medium for *S. aureus* was Trypticase soy agar (TSA; 22091) with added 7.5% NaCl, called Staphylococcus isolation agar (SIA) (*28*). Both PIA and SIA were prepared following the protocols provided by the producer. Carbon-free M9-minimal medium (M6030) used for flow cytometry was prepared using 100 mL of 5× M9 minimal salts, with an additional 1 mL of 1 M MgSO_4_ and 50 μL of 1 M CaCl_2_ and make up to a volume of 500 mL using Milli-Q water. M9 was then filtered through a 0.22 μm Minisart® Syringe Filter (Sartorius, Germany). Phosphate buffered saline solution (PBS; 4417) was prepared by dissolving one tablet in 200 mL of Milli-Q water, autoclaved and used for growth and killing assays. Ciprofloxacin (17850) stock solution was obtained by dissolving antibiotic powder in 0.1 M HCl at a concentration of 10 mg mL^-1^ and then diluted with LB to the required experimental concentration. Vancomycin (94747) stock solution was obtained by dissolving powder in Milli-Q water at a concentration of 10 mg mL^-1^ and then diluted with LB to required experimental concentration. Vancomycin fluorescent analogue was designed and synthesised based on structure-activity-relationship studies and protocols reported in prior publications (*59*, *94*), introducing an azidolysine residue for the subsequent ‘click’ reaction with nitrobenzoxadiazole (NBD)-alkyne (*95*).

*S. aureus* ATCC 25923 and *P. aeruginosa* PA14 were purchased from Dharmacon™ (Horizon Discovery, UK). *S. aureus* SH1000 mCherry was kindly provided by Prof Simon Foster, University of Sheffield, *P. aeruginosa* PA14 *flgK*::Tn5B30(Tc) was kindly provided by Prof George O’Toole, Dartmouth College, USA, *P. aeruginosa* PAO1 and PAO1 Δ*pvdD*Δ*pchEF* were kindly provided by Prof Pierre Cornelis, Universiteit Brussels, *S. aureus* and *P. aeruginosa* clinical isolates RYC157 and RYC165 were provided by Dr Maria Garcia and Prof Rafael Canton, IRYCIS, Madrid. All strains were stored in a 50% glycerol, 50% Milli-Q water stock at -70 °C. Streak plates for each strain were produced by thawing a small aliquot of the corresponding glycerol stock every week and plated onto LB agar plates.

### Mono- and co-culture growth assays

Overnight *S. aureus* (strain ATCC25923, RYC157 or RYC165) or *P. aeruginosa* (strain PA14 or RYC157) mono-cultures were prepared by inoculating 100 mL fresh LB medium with a single bacterial colony from a streak plate and incubated on a shaking platform at 200 rpm and 37 °C for 17 h. *S. aureus* and *P. aeruginosa* mono-cultures and co-cultures for growth assays were prepared by transferring 100 μL of *S. aureus* or 100 μL *P. aeruginosa* (or both together for co-culture) overnight mono-cultures into 100 mL LB and incubated on a shaking platform at 200 rpm and 37 °C for 13 h while monitoring bacterial growth hourly. To do so, each culture was sampled every hour in triplicate by taking three 500 µL aliquots, these aliquots were centrifuged at 13000 rpm for 5 min, the supernatant was removed, each pellet was resuspended in PBS and serial dilutions were plated using 10 uL droplets on LB agar plates or PIA, TSA agar plates for the isolation of *P. aeruginosa* and *S. aureus*, respectively (*28*) and colony forming units (CFU) counted. For both mono-cultures and co-cultures growth assays all the plates after bacterial plating were dried and incubate for 16 h at 37 °C, before counting CFU.

### Determination of minimum inhibitory concentration

Single *S. aureus* or *P. aeruginosa* bacterial colonies were picked and cultured overnight in LB at 37 °C for 17 h, then diluted 40-fold (by adding 1 mL of overnight culture to 39 mL of LB) and grown at 37 °C on a shaking platform at 200 rpm to an OD_600_ of 0.5. The mid-log phase cultures (i.e. OD_600_ = 0.5) were diluted to 10^6^ CFU by removing from 15 mL of LB 30 µL and replacing with 30 µL of exponential phase bacterial solution. 90 µL of LB were added to the first column, and 50 µL to all other wells of a 96-well plate. 10 µL of 1.28 mg mL^-1^ of each unlabelled antibiotic or fluorescent antibiotic derivative stocks were added to the first column of the 96 well plate. 50 µL were then withdrawn from the first column and serially transferred to the next column until 50 µL withdrawn from the last column was discharged. After this procedure all wells contained 50 µL of antibiotic in LB solution. 50 µL of exponential phase bacterial solution was then added to each well, to give a final concentration of 5×10^5^ CFU mL^-1^. Each antibiotic in LB was tested in triplicates (three rows of a 96 well plate). Each plate also contained two rows of 12 positive control experiments (i.e. bacteria in LB without antibiotics added) and two rows of 12 negative control experiments (i.e LB only). Plates were covered with aluminium foil and incubated at 37 °C overnight. The minimum inhibitory concentration (MIC) of each antibiotic was determined visually, with the MIC being the lowest concentration well with no visible growth compared to the positive control experiments (*96*).

### Time-kill assays

Exponential (i.e. 3 h) or stationary phase (i.e. 10 h) *S. aureus* (strain ATCC25923) or *P. aeruginosa* (strain PA14) mono- or co-cultures were prepared as described above. Four aliquots of each mono- or co-culture were diluted to an OD of 0.03 in 50 mL of LB in four separate flasks. The four flasks were challenged by using four different ciprofloxacin or vancomycin concentrations, i.e. 0×, 1×, 10×, or 25× the MIC value measured against *S. aureus*. In order to perform CFU counting we withdrew three 500 µL aliquots from each flask at t = 0, 1, 2, 3, 4 and 5 h after the addition of the antibiotic and added them to Eppendorf tubes that were incubated at 200 rpm at 37 °C. At each time point three aliquots were centrifuged at 13000 rpm for 5 min, the supernatant was then removed, each pellet was resuspended in PBS and serial dilutions (made in the 96-well plate) were plated using 10 uL droplets on LB agar plates or PIA, TSA agar plates for the isolation of *P. aeruginosa* and *S. aureus*, respectively (*28*). Determination of CFU was obtained after overnight incubation of each plate at 37 °C. Means and standard deviations of measurements carried out in biological triplicate were calculated and graphs plotted using GraphPad Prism 10.

### Fabrication of microfluidic devices

The mould for the mother machine microfluidic device was fabricated using previously established multi-layer photolithography processes (*97*, *98*). This mould is equipped with twenty 1 µm high circular chambers with a diameter of 150 µm. These chambers are connected to a main delivery chamber via channels of different width in range from 20 to 50 µm. The main delivery chamber has a height of 20 µm and a width of 200 µm. Polydimethylsiloxane (PDMS) replicas of this device were realised as previously described (*99*). Briefly, a 10:1 (base:curing agent) PDMS mixture was cast on the mould and cured at 70 °C for 120 min in an oven (*100*). The cured PDMS was peeled from the epoxy mould and fluidic accesses were created by using a 0.75 mm biopsy punch (Harris Uni-Core, WPI). The PDMS chip was washed with absolute ethanol and dried with a nitrogen stream the day before the experiment. The chip was then irreversibly sealed on a glass coverslip (22×50 mm, 0.13-0.17 mm thick, Fisher) by exposing both the chip and glass surfaces to oxygen plasma treatment (10 s exposure to 30 W plasma power, Plasma etcher, Diener, Royal Oak, MI, USA) on the day of the experiment as previously described (*101*). We have also made available a step-by-step experimental protocol for the fabrication and handling of microfluidic devices for investigating the interactions between antibiotics and individual bacteria (*102*).

### Microfluidics-based time-lapse microscopy

Overnight *S. aureus* ATCC25923 or SH1000 mCherry and *P. aeruginosa* PA14 Δ*flgK* mono-cultures or co-cultures were prepared as described above. A 50 mL aliquot of each culture was centrifuged for 10 min at 4000 rpm and 20 °C. The supernatant was filtered twice (Medical Millex-GS Filter, 0.22 μm, Millipore Corp.) to remove bacterial debris from the solution and used to resuspend the bacteria in their spent LB to an OD_600_ of 10 as previously described(*103*). A 10 μL aliquot of this suspension was injected by using pipette in the above described microfluidic device. The high bacterial concentration favours bacteria entering the lateral chambers from the main delivery chamber of the microfluidic device (*39*). The lateral chambers were filled seconds after the bacterial injection. The microfluidic device was completed by the integration of fluorinated ethylene propylene tubing (P68514, 1/32” × 0.20mm ID, Restek). The inlet tubing was connected to the inlet reservoir which was connected to a computerised pressure-based flow control system (MFCS-4C, Fluigent). This instrumentation was controlled by MAESFLO software (Fluigent) (*104*). The chip was mounted on an inverted microscope (IX73 Olympus, Tokyo, Japan) and the bacteria remaining in the delivery chamber of the microfluidic device were washed into the outlet tubing and into the waste reservoir by flowing LB medium or LB medium containing ciprofloxacin or vancomycin at 200 μL h^-1^ for 2 min. In the unchallenged mono- and co-culture experiments LB was supplied to the bacteria at 100 μL h^-1^ for *S. aureus* mono-cultures and 40 μL h^-1^ for *P. aeruginosa* mono-cultures or co-cultures—for 9 h. The lower flow rate for *P. aeruginosa* mono-cultures or co-cultures was used due to their higher motility compared to *S. aureus*. In the mono- and co-culture experiments with ciprofloxacin or vancomycin challenge, LB medium containing either ciprofloxacin at a concentration of 0.5 µg mL^-1^ or vancomycin at a concentration of 1 µg mL^-1^ was supplied for 4 h at a flow rate of 100 or 40 μL h^-1^ followed by delivery of LB medium only for 3 h at a flow rate of 100 or 40 μL h^-1^. During the change of medium from LB containing antibiotic to LB only the main channel was flushed for 8 min at 200 μL h^-1^ to ensure that the antibiotic was fully removed from the device. Bright-field images were acquired every 10 min during entire experiment. Images were collected via a 60×, 1.2 N.A. objective (UPLSAPO60XW, Olympus) and a sCMOS camera (Zyla 4.2, Andor, Belfast, UK). The region of interest of the camera was adjusted to visualise one of the lateral chambers in the device and images of four different lateral chambers of the microfluidic device were acquired at each time point. The device was moved by two automated stages (M-545.USC and P-545.3C7, Physik Instrumente, Karlsruhe, Germany, for coarse and fine movements, respectively). A corresponding fluorescence image was acquired by exposing the bacteria for 0.03 s to the green excitation band of a broad-spectrum LED (CoolLED pE300white, maximal power = 200 mW Andover, UK) at 30% of its intensity (with a power associated with the beam light of 8 mW at the sample plane) and a TEXAS RED filter to quantify mCherry fluorescence from each bacterium. Fluorescence images of SH1000 mCherry were acquired every 30 min during the entire experiment, to avoid overexposing the bacteria to the LED light. In the end of all experiments propidium iodide (P1304MP, Invitrogen), was added to the main delivery chamber in order to detect cells with compromised membranes as previously described (*39*). The entire assay was carried out at 37 °C in an environmental chamber (Solent Scientific, Portsmouth, UK) surrounding the microscope and microfluidics equipment.

### Image and data analysis

Images were processed using ImageJ software as previously described (*105*, *106*), counting and tracking individual *S. aureus* bacteria throughout each experiment, whereas individual *P. aeruginosa* bacteria were counted but not tracked at each time point in each experiment due to their higher motility. In order to estimate mCherry fluorescence from individual *S. aureus* bacteria, a circle was drawn around each bacterium in each bright-field image at every time point. The same circle was then used in the corresponding fluorescence image to measure the mean fluorescence intensity for each bacterium. The same circle was then moved to a section of the same chamber that did not host any bacteria in order to measure the background fluorescence. This mean background fluorescence value was subtracted from the bacterium’s fluorescence value. All data were then analysed and plotted using GraphPad Prism 10. Statistical significance was tested using either paired or unpaired, two-tailed, Welch’s *t*-test. Pearson correlation, means, standard deviations, coefficients of variation and medians were also calculated using GraphPad Prism 10.

### Flow cytometric accumulation assays

The accumulation of vancomycin-NBD in individual cells from *S. aureus* ATCC25923 or SH1000 mCherry and *P. aeruginosa* PA14 Δ*flgK* mono- or co-cultures were carried out as previously described (*47*, *95*). Briefly, overnight *S. aureus* ATCC25923, SH1000 mCherry and *P. aeruginosa* PA14 ΔflgK mono-cultures and 10 h subcultures were prepared as described above. 10 h old mono-cultures and co-cultures were then adjusted to an OD_600_ of 5 for S. aureus mono-cultures and 2 for co-cultures and *P. aeruginosa* mono-cultures. 10 h old mono-cultures and co-cultures were transferred to a microcentrifuge tube for centrifugation in a SLS4600 microcentrifuge (Scientific Laboratory Supplies, UK) at 4,000 ×*g* and room temperature for 5 min. The supernatant was removed, and pellets were resuspended in M9 medium (control; samples without any antibiotic) or vancomycin-NBD at a concentration of 16 µg mL^-1^ by diluting in M9 medium at a final volume of 130 μL. Each sample containing antibiotic was incubated at 37 °C and shaken at 1,000 rpm on a ThermoMixer^®^ C (Eppendorf, Germany) for 1 h. For control samples (bacterial solutions without any antibiotic), pellets were dissolved in 130 μL of M9 and were then diluted ten times in M9. 2 μL of these samples were then transferred to 96-well plates containing 98 μL of double filtered M9, giving a final dilution of 500 times. For the samples containing vancomycin, 30 μL of such samples were then transferred to microcentrifuge tubes at appropriate time points (20 min, 40 min and 60 min). A washing step was immediately performed on the vancomycin containing aliquots to eliminate any unaccumulated drug via centrifugation at 4000 ×*g* for 5 min and its supernatant removed. Then we added 300 μL of filtered M9 to the pellets to perform a 10 times dilution and added 2 μL of these samples to 98 μL of M9 in wells of a 96 well plate. Flow cytometric measurements were performed on the CytoFLEX S Flow Cytometer (Beckman Coulter, USA) equipped with a 488 nm (50 mW) and a 405 nm (80 mW) laser. Vancomycin-NBD and mCherry fluorescence of individual bacteria was measured using the FITC-A channel (488 nm, 525/40BP) and the ECD channel (488 nm, 610/20BP), respectively. Cell size was measured using SSC-A (forward scatter) and Violet SSC-A (side scatter) channels. Avalanche photodiode gains of FSC: 1,000, SSC: 500, Violet SSC: 1, FITC: 250, ECD 250 and a threshold value of SSC-A: 10,000 to limit background noise was used. Bacteria were gated to separate cells from background noise by plotting SSC-A and Violet SSC-A. Background noise was then further separated based on cellular autofluorescence measured on the FITC-A channel for cells not treated with fluorescent-peptides. An additional gate on the ECD-A channel was then used for cells treated with vancomycin-NBD to further separate background noise from mCherry expressing *S. aureus*. CytoFLEX Sheath Fluid (Beckman Coulter, USA) was used as sheath fluid. Data was collected using CytExpert software (Beckman Coulter, USA) and exported to FlowJo™ version 10.9 software (BD Biosciences, USA) for analysis.

### Transcriptomic analysis

Exponential (i.e. 3 h) and stationary phase (i.e. 10 h) *S. aureus* ATCC25923 and *P. aeruginosa* PA14 Δ*flgK* mono- or co-cultures were prepared as described above. RNA extractions were performed on biological triplicate of each culture condition as previously described (*48*), RNA extractions for stationary phase co-cultures were performed in six biological replicates to compensate for the relative low content of *S. aureus* with respect to *P. aeruginosa* cells. Briefly, to make an RNA extraction from one biological replicate, 100 mL of 3 h or 10 h old bacterial cultures were centrifuged at 4000 rpm for 15 min. The supernatant was then removed. 100 μL of bacterial solution was then transferred into an RNase free tube and 200 μL of RNA-protect bacteria reagent was added, vortexed for 5 s and then incubated for 5 min at room temperature. The sample was then centrifuged at 5000 g for 10 min and the supernatant was removed. Enzymatic lysis of *S. aureus* bacteria was carried out using lysostaphin stock (2 mg mL^-1^) in TE buffer. Briefly, 200 μL of 100 μg mL^-1^ lysostaphin was added to the bacterial pellet, vortexed and incubated for 1 h at 37 °C. Then the sample was vortexed again, proteinase K was added and the sample was incubated for 10 min at 37 °C, while vortexing every 2 min. Enzymatic lysis of *P. aeruginosa* bacteria was carried out using lysozyme at a concentration of 15 mg mL^-1^ in TE buffer and proteinase K digestion of bacteria was performed following protocol 4 in the RNAprotect^®^ Bacteria Reagent Handbook (Qiagen, Germany). Purification of total RNA from bacterial lysate using the RNeasy^®^ Mini Kit (Qiagen, Germany) was carried out following protocol 7 in the handbook. A further on-column DNase digestion using the RNase-Free DNase set (Qiagen, Germany) was performed following appendix B in the handbook. RNA concentration and quality were assessed using the Agilent High Sensitivity RNA ScreenTape System (Agilent, USA) following the provided protocol. Samples for the analysis were normalised to a mean RNA concentration of 40 ng μL^-1^ and the measured mean RNA integrity number equivalent (RIN^e^) score was 9.1. To start the library preparation, we used 400 ng total RNA. rRNA depletion was performed with the Illumina Stranded Total RNA Prep Ligation with Ribo-Zero Plus Microbiome kit (Illumina, USA) following the manufacturer’s instructions. cDNA libraries were compiled using as well the Illumina Stranded Total RNA Prep Ligation with Ribo-Zero Plus Microbiome kit (Illumina, USA). Transcript abundance was quantified using Salmon for each gene in all samples. Differential gene expression was performed with DESeq2 in R software to quantify the log_2_ fold-change in transcript reads (*49*) for each gene and subsequent principal component analysis using DESeq2 and a built-in R method (prcomp) (*45*). Gene ontology enrichment analysis was performed using the clusterProfiler package (version 4.10.0) for R (*107*, *108*). Enrichment in terms belonging to the “Biological Process” ontology was calculated for each gene cluster, relative to the set of all genes quantified in the experiment, via a one-sided Fisher exact test (hypergeometric test). *P* values were adjusted for false discovery by using the method of Benjamini and Hochberg. Finally, the lists of significantly enriched terms were simplified to remove redundant terms, as assessed via their semantic similarity to other enriched terms, using clusterProfiler’s simplify function.

### RT and qPCR

One hundred nanograms of RNA were used for reverse transcription (RT) with random hexamer primers, following the protocol of the High-Capacity cDNA Reverse Transcription Kit (Applied Biosystems). No-RT controls were included for all samples to confirm that transcript levels were not due to residual DNA. The RT reaction was carried out at 25 °C for 10 minutes for primer annealing, followed by cDNA synthesis at 37 °C for 120 minutes and termination at 85 °C for 5 minutes in a final volume of 20 µL. The resulting cDNA samples were diluted 5-fold in 10 mM Tris-HCl (pH 8) and stored at −20 °C prior to qPCR analysis. Real-time qPCR reactions were performed in a total volume of 20 µL containing 1X PowerUp™ SYBR Green Master Mix (Applied Biosystems), 500 nM of each forward and reverse primer (Supplementary Table X), and 5 ng of template cDNA. Amplification was conducted on a QuantStudio 1 Real-Time PCR system (Thermo Fisher Scientific) using the following cycling conditions: an initial uracil-DNA glycosylase (UDG) activation step at 50 °C for 2 minutes, followed by DNA polymerase activation at 95 °C for 2 minutes. PCR cycling consisted of 35 cycles of denaturation at 95 °C for 15 seconds and combined annealing/extension at 60 °C for 1 minute. At the end of amplification, a melt curve analysis was conducted by increasing the temperature from 60 °C to 95 °C at a rate of 0.1 °C/second, with continuous fluorescence data acquisition. Threshold cycle (Ct) values were determined using the Design & Analysis (DA2) software v2.6.0 (Thermo Fisher Scientific). Changes in gene expression levels (log2 fold change) were calculated using the ΔΔCt method, with an internal control gene selected for normalization for each bacterial strain.

### Lipid extraction and lipidomic analysis

Lipidomic analysis of *S. aureus* ATCC25923 and *P. aeruginosa* PA14 mono- or co-cultures untreated or treated with vancomycin at a concentration of 1 µg mL^-1^ were carried out as previously described (*47*). Briefly, lipid extractions were performed following a modified version of the Folch extraction (*54*, *109*): 100 μL of sample was added to 250 μL of methanol and 125 μL of chloroform in a microcentrifuge tube; samples were incubated for 60 min and vortexed every 15 min. Then, 380 μL of chloroform and 90 μL of 0.2 M potassium chloride was added to each sample. Cells were centrifuged at 14,000 ×*g* for 10 min to obtain a lipophilic phase which was transferred to a glass vial and dried under a nitrogen stream. The lipophilic phase was then resuspended in 20 μL of a methanol:chloroform solution (1:1 *v/v*), then 980 μL of an isopropanol:acetonitrile:water solution (2:1:1, *v/v/v*) was added. Samples were analysed on a Agilent 6560 Q-TOF-MS (Agilent, USA) coupled with a Agilent 1290 Infinity II LC system (Agilent, USA). An aliquot of 0.5 μL for positive ionisation mode and 2 μL for negative ionisation mode from each sample was injected into a Kinetex^®^ 5 μm EVO C18 100 A, 150 mm × 2.1 μm column (Phenomenex, USA). The column was maintained at 50 °C at a flow rate of 0.3 mL min^-1^. For the positive ionisation mode, the mobile phases consisted of acetonitrile:water (2:3 *v/v*) with ammonium formate (10 mM) and acetonitrile:isopropanol (1:9, *v/v*) with ammonium formate (10 mM). For the negative ionisation mode, the mobile phases consisted of (A) acetonitrile:water (2:3 *v/v*) with ammonium acetate (10 mM) and acetonitrile:isopropanol (1:9, *v/v*) with ammonium acetate (10 mM). The chromatographic separation was obtained with the following gradient: 0-1 min 70% B; 1-3.5 min 86% B; 3.5-10 min 86% B; 10.1-17 min 100% B; 17.1-10 min 70% B. The mass spectrometry platform was equipped with an Agilent Jet Stream Technology Ion Source (Agilent, USA), which was operated in both positive and negative ion modes with the following parameters: gas temperature (200 °C); gas flow (nitrogen), 10 L min^-1^; nebuliser gas (nitrogen), 50 psig; sheath gas temperature (300 °C); sheath gas flow, 12 L min^-1^; capillary voltage 3500 V for positive, and 3000 V for negative; nozzle voltage 0 V; fragmentor 150 V; skimmer 65 V, octupole RF 7550 V; mass range, 40-1,700 *m/z*; capillary voltage, 3.5 kV; collision energy 20 eV in positive, and 25 eV in negative mode. MassHunter software (Agilent, USA) was used for instrument control.

### Mathematical model to describe *S. aureus* and *P. aeruginosa* population dynamics

In order to model the population dynamics recorded for mono- and co-cultures of different strains of *S. aureus* and *P. aeruginosa* in well-mixed flasks, where nutrients become limiting, we used the following set of differential equations:

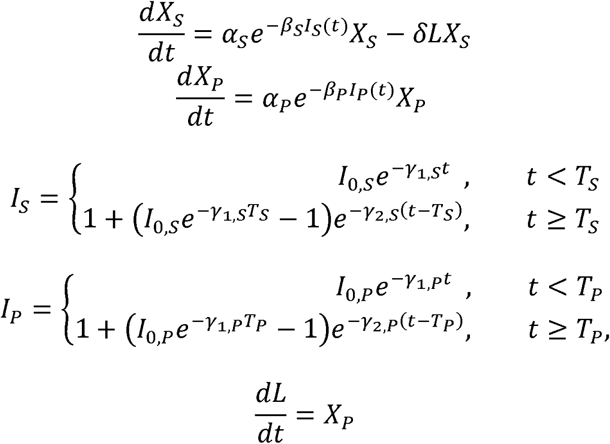

where *X* is the number of *S. aureus* (*S*) and *P. aeruginosa* (*P*) cell counts,β, is a constant of biochemical process activation when nutrients are in abundance, _2_ is the rate of biochemical process inactivation when nutrients are limiting, *L* is the concentration of compound(s) secreted by *P. aeruginosa* and affecting the growth of *S. aureus*, is a constant representing inactive, γ_1_ is the time at which nutrients become limiting.

In the microfluidic chambers where nutrients do not become limiting (i.e. an open system), the growth of *S. aureus* in mono-culture or in co-culture with *P. aeruginosa* can be described by the following differential equation:

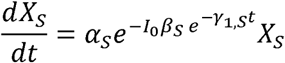

Whereas the growth of *P. aeruginosa* in co-culture with *S. aureus* in the microfluidic chambers can be described by the following differential equations:

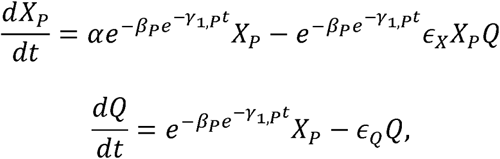

Where ε*_X_* is the coefficient of the rate of escape of *P. aeruginosa* from microfluidic chambers, is the coefficient of the rate of escape of quorum-sensing molecules secreted by *P*.

*aeruginosa*, ε_Q_ is the concentration of quorum-sensing molecules secreted by *P. aeruginosa*.

The equations above were fitted to our corresponding population dynamics data via least-squares regression with parameter estimations reported in Tables S1-S3. A full description of the model can be found at https://github.com/tt386/GeometricBacterialGrowth/tree/main.

**Supplemetary Materials**

This document includes:

Figs. S1 to S19 Tables S1 to S3

Legends for Datasets S1 to S20

**Other Supplementary Material for this manuscript includes the following:**

Datasets S1 to S20

## Acknowledgments

We thank Dr Seana Duggan and Prof Simon Foster for providing the *S. aureus* SH1000 mCherry strain.

## Author contributions

Conceptualization, S.P.; methodology, U.L., N.O.G., G.C., T.T., S.Z., G.T., A.D.V., M.G.-C., P.A.O.N., A.F., A.R.J., A.J.Y., R.C., P.C., M.A.T.B., A.W., K.T.A. and S.P.; formal analysis, U.L., N.O.G., G.C., T.T., S.Z., G.T., P.A.O.N., P.C., A.W., K.T.A. and S.P.; generation of figures, U.L. and S.P.; investigation, U.L., N.O.G., G.C., T.T., S.Z., G.T. and S.P.; resources, A.J.Y., R.C., P.C., M.A.T.B., A.W., K.T.A. and S.P.; data curation, U.L. and S.P., writing – original draft, U.L. and S.P.; writing – review & editing, U.L., N.O.G., G.C., T.T., S.Z., G.T., A.D.V., M.G.-C, P.A.O.N., A.F., A.R.J., A.J.Y., R.C., P.C., M.A.T.B., A.W., K.T.A. and S.P.; visualization, U.L. and S.P.; supervision, A.J.Y., P.C., A.W., K.T.A. and S.P.; project administration, S.P.; funding acquisition, U.L., A.J., A.J.Y., M.A.T.B., A.W., K.T.A. and S.P..

## Data and materials availability

All data are available in the main text or the supplementary materials.

