## Supplementary Materials for "Competition with *Pseudomonas aeruginosa* induces *Staphylococcus aureus* in an antibiotic-tolerant viable but non culturable state"

- Christèle Aubry, Amanda Luraschi-Eggemann, Maria Georgevia & Gilbert Greub, Lausanne, Switzerland
- Gino Cathomen, Danuta Cichocka & Alexander Sturm, Muttenz, Switzerland
- Maria Gracia-Castillo & Rafael Canton, Madrid, Spain
- Nicola Oswaldo Trinler & Susanne Häußler, Helmholtz, Germany
- Niilo Kaldalu, Kristiina Vind & Tanel Tenson, Tartu, Estonia
- Tailise Rodrigues, Maureen Micaletto, Urszula Łapińska & Stefano Pagliara, Exeter, UK

**Supplementary figures**

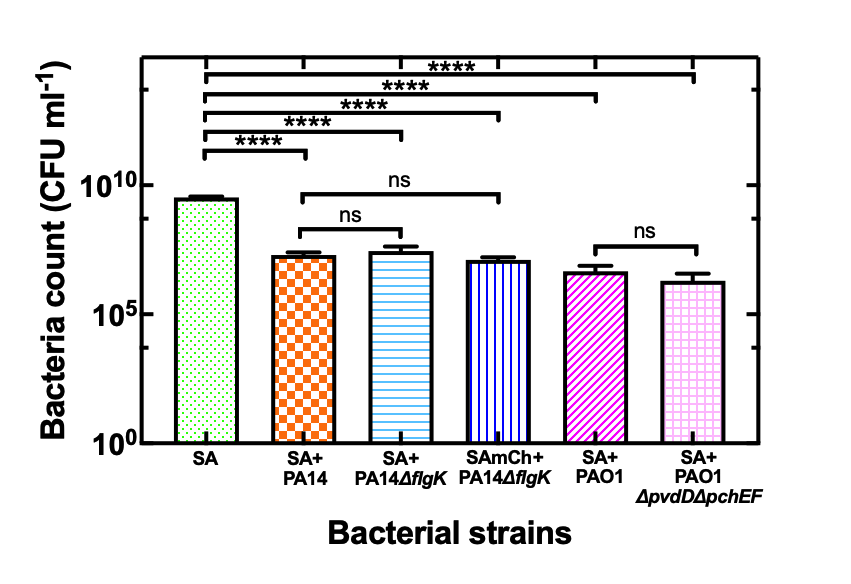

Figure S1 Population size for *S. aureus* (SA) after 13 h in mono-culture (green bar) or 13 h in co-culture with either *P. aeruginosa* PA14 (orange bar), *P. aeruginosa* PA14 Δ*flgK* (cyan bar), *P. aeruginosa* PAO1 (magenta bar) or *P. aeruginosa* PAO1 Δ*pvdD*Δ*pchEF* (pink bar). Population size for *S. aureus* SH1000 mCherry (SAmCh) after 13 h in co-culture with *P. aeruginosa* PA14 Δ*flgK* (blue bar) and enumerated using selective plates. Data points are the mean and standard deviation of colony forming unit (CFU) measurements carried out in biological triplicate each consisting of technical duplicate. Numerical values for each replicate are reported in Data E in S1 File. **** denotes a p-value < 0.0001, * denotes a p-value < 0.05 ns denotes non significant.

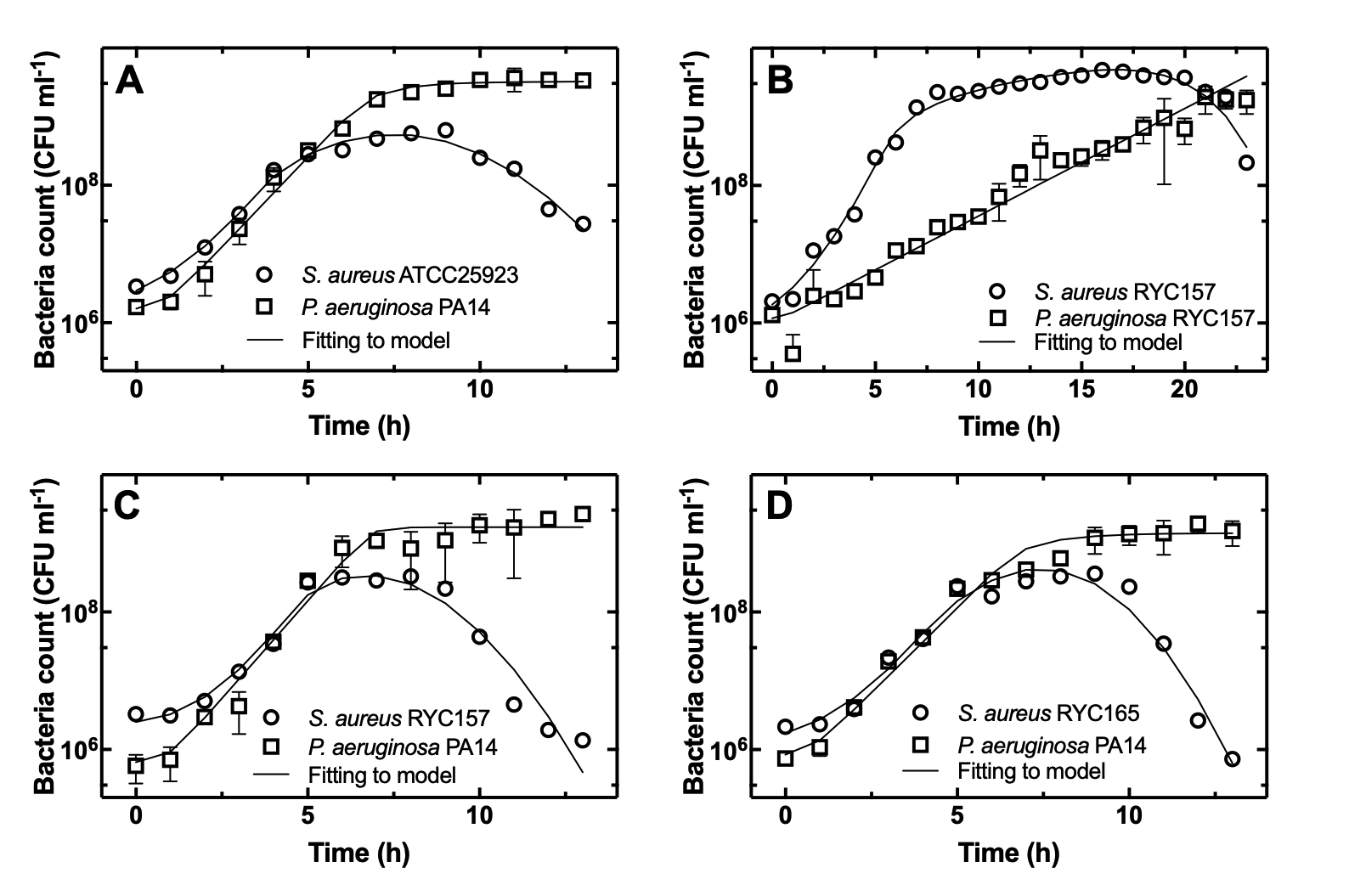

Figure S2 Temporal dependence of bacterial population size for (a) *S. aureus* ATCC25923 and *P. aeruginosa* PA14, (b) *S. aureus* RYC157 and *P. aeruginosa* RYC157, (c) *S. aureus* RYC157 and *P. aeruginosa* PA14, or (d) *S. aureus* RY165 and *P. aeruginosa* PA14 in co-culture in well-mixed flasks and enumerated using selective plates. Data points are the mean and standard deviation of colony forming unit (CFU) measurements carried out in biological triplicate each consisting of biological duplicate. Numerical values for each replicate are reported in Data B, D and F-K in S1 File. Solid lines are fitting of the data to our mathematical model, with parameter estimates reported in Table S1. The data in A are reproduced from Figure 1A and 1B

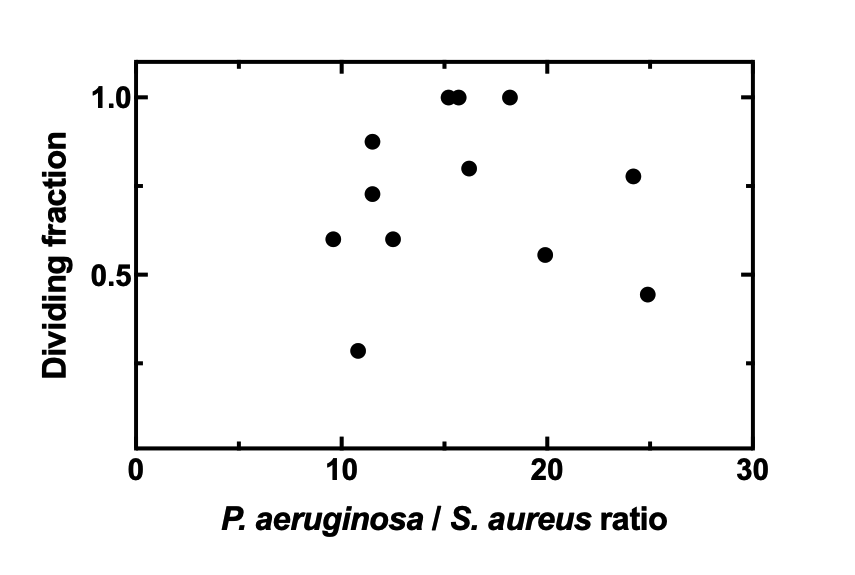

Figure S3 Dependence of the *S. aureus* ATCC25923 dividing fraction over 9 h regrowth in LB on the initial ratio of the number of *P. aeruginosa* PA14 cells over the number of *S. aureus* ATCC25923 cells within each of 12 different microfluidic chambers investigated (4 chambers from each biological replicate) at t = 0. Numerical values are reported in Data O-P in S1 File.

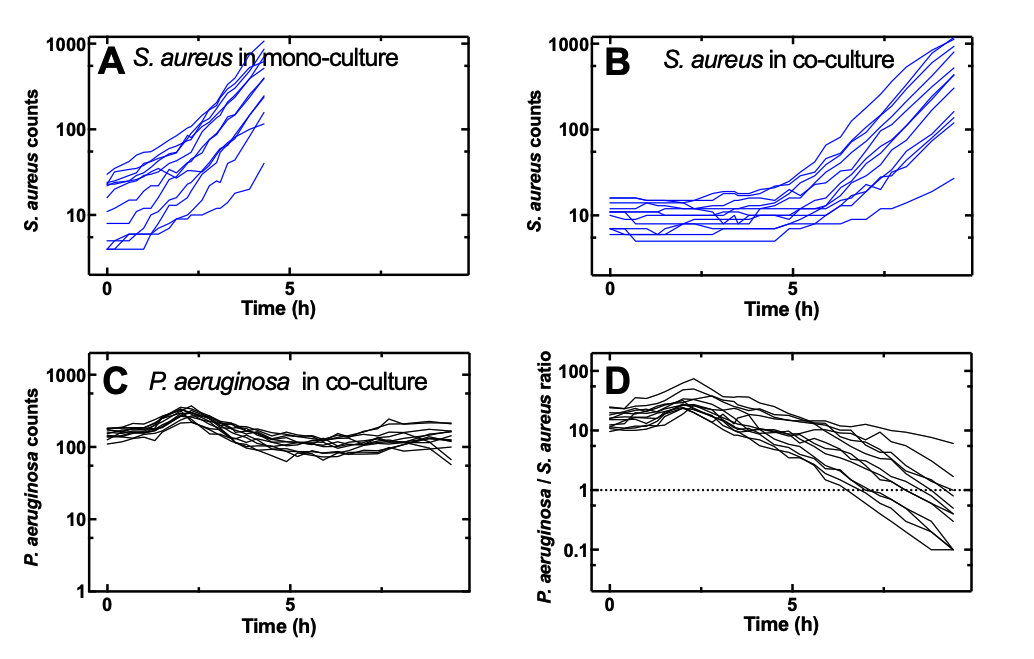

Figure S4 Temporal dependence of the number of *S. aureus* ATCC25923 cells in 12 different microfluidic chambers (4 chambers from each biological replicate) harvested from (a) 10 h old mono-cultures or (b) 10 h old co-cultures and exposed to LB medium for 4.5 h or 9.5 h within each chamber. (c) Corresponding temporal dependence of the number of *P. aeruginosa* PA14 cells. (d) Corresponding temporal dependence of the ratio of the number of *P. aeruginosa* cells over the number of *S. aureus* cells. Numerical values are reported in Data N-P in S1 File.

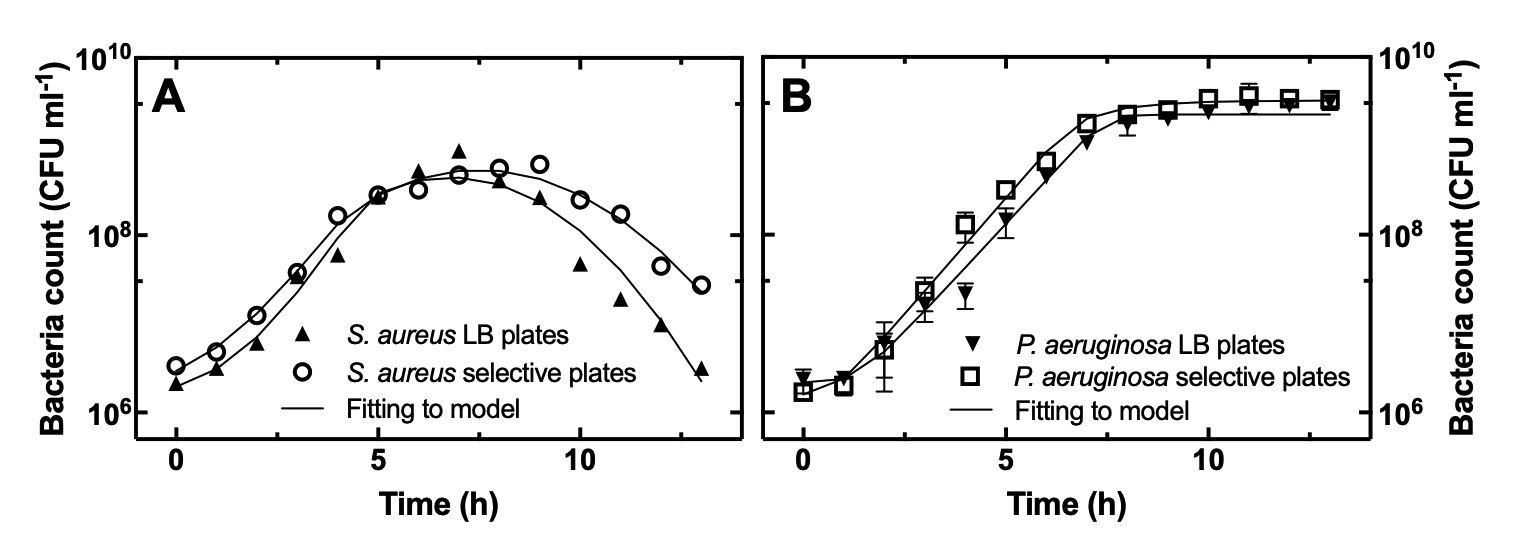

Figure S5 Temporal dependence of bacterial population size for (a) *S. aureus* ATCC25923 in co-culture with *P. aeruginosa* PA14 in well-mixed flasks and enumerated using LB plates (open circles) or selective plates (filled upward triangles) or (b) for *P. aeruginosa* PA14 in co-culture with *S. aureus* ATCC25923 in well-mixed flasks and enumerated using LB plates (open squares) or selective plates (filled downward triangles). Data points are the mean and standard deviation of colony forming unit (CFU) measurements carried out in biological triplicate each consisting of biological duplicate. Numerical values for each replicate are reported in Data B, D and R-S in S1 File. Solid lines are fitting of the data to our mathematical model with parameter estimates reported in Table S1. The data obtained using selective plates are reproduced from Figure 1A and 1B.

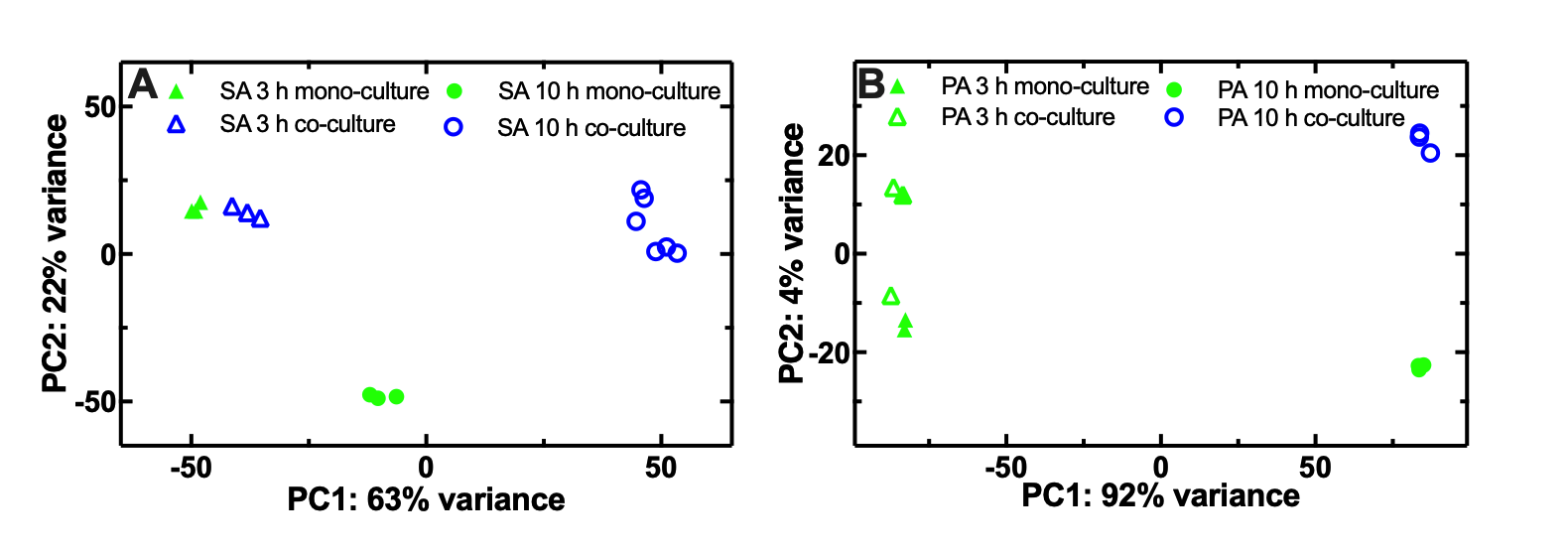

Figure S6 (a) Principal component analysis of replicate transcriptomes of *S. aureus* ATCC25923 after 3 h of mono-culture (filled triangles) or co-culture with *P. aeruginosa* PA14 (open triangles) or after 10 h of mono-culture (filled circles) or co-culture with *P. aeruginosa* PA14 (open circles). (b) Principal component analysis of replicate transcriptomes of *P. aeruginosa* PA14 after 3 h of mono-culture (filled triangles) or co-culture with *S. aureus* ATCC25923 (open triangles) or after 10 h of mono-culture (filled circles) or co-culture with *S. aureus* ATCC25923 (open circles). Each transcriptome was measured in biological triplicate apart from the transcriptome of *S. aureus* ATCC25923 after 10 h of co-culture which was measured in six biological replicates because of the lower RNA yield for this experimental condition. Numerical values are reported in S2 File.

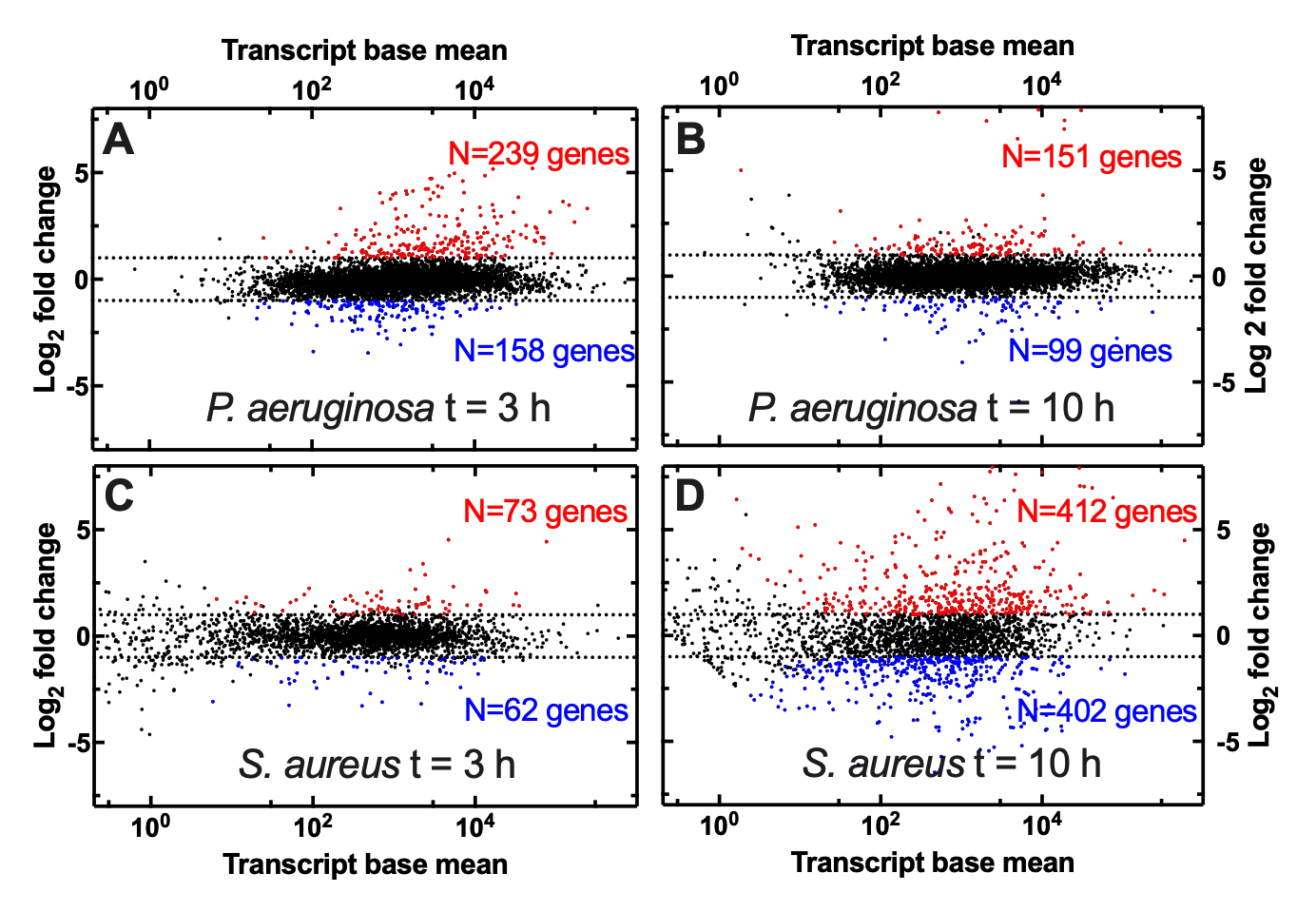

Figure S7 Whole-genome differential gene expression in *P. aeruginosa* PA14 after (a) 3 h or (b) 10 h co-culture with *S. aureus* ATCC25923 with respect to mono-culture, or in *S. aureus* ATCC25923 after (c) 3 h or (d) 10 h co-culture with *P. aeruginosa* PA14 with respect to mono-culture. Each dot represents the mean log_2_ fold change for each *P. aeruginosa* or *S. aureus* gene calculated from measurements carried out in biological triplicate. Dotted lines indicate a log_2_ fold change of 1 or -1, red and blue dots indicate upregulated or downregulated genes, respectively, in co-culture compared to mono-culture with a log_2_ fold change larger or lower than 1, respectively, and an adjusted p-valued smaller than 0.05. Numerical values are reported in S3-S6 Files.

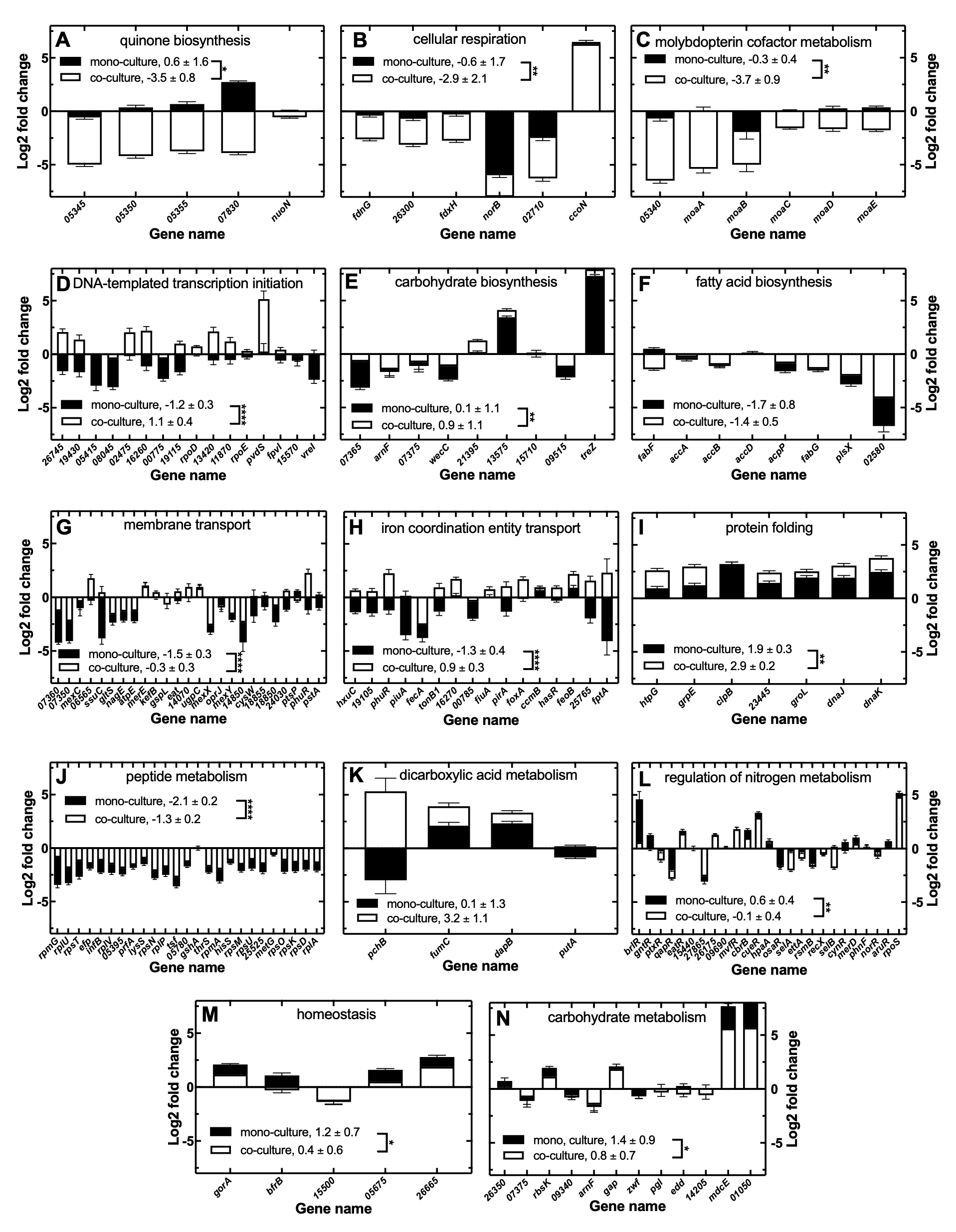

Figure S8 Differential expression of *P. aeruginosa* PA14 genes at t = 10 h compared to t = 3 h in mono-culture (filled bars) or co-culture with *S. aureus* ATCC25923 (open bars) for biological processes that are significantly enriched in the comparisons of the transcriptomes of *P. aeruginosa* PA14 in mono-culture vs co-culture either at t = 3h or at t = 10 h. The bars and error bars report the mean and standard error of the log_2_ fold change calculated for each gene from transcriptomic measurements carried out in biological triplicate. The numbers in each graph report the mean and standard error of the log_2_ fold change measured across all genes in each process in mono-culture or co-culture. ****: p-value < 0.0005, **: p-value < 0.01, * < p-value < 0.05 according to pair-wise t-tests between the log_2_ fold change measured for each gene in mono-culture vs co-culture. log_2_ fold change data are reported for all genes constituting the enrichment core in each process apart from membrane transport, peptide metabolism and regulation of metabolism processes. These processes contained more than 25 enrichment core genes that would have been difficult to visualise. Therefore, 25 genes were randomly selected for these three processes for visualization, whereas numerical values for all genes are reported in S9-S10 Files.

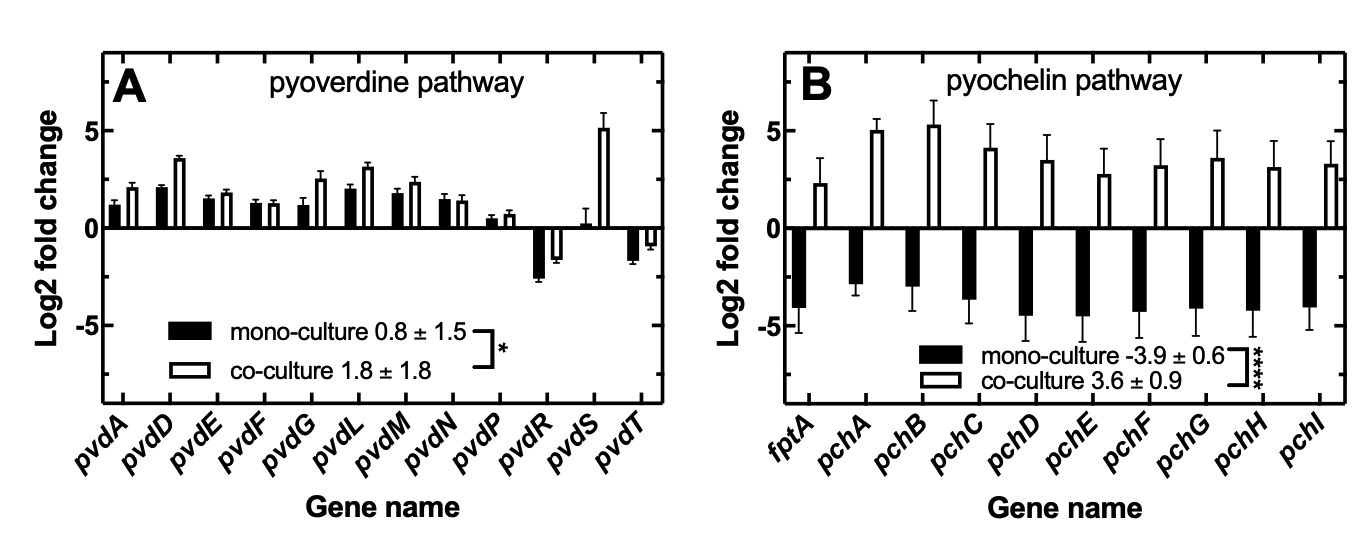

Figure S9 Differential expression of genes in the (a) pyoverdine or (b) pyochelin siderophore pathways at t = 10 h compared to t = 3 h for *P. aeruginosa* in mono-culture (filled bars) or co-culture with *S. aureus* (open bars). The bars and error bars report the mean and standard error of the log_2_ fold change calculated for each gene from transcriptomic measurements carried out in biological triplicate. The numbers in each graph report the mean and standard error of the log_2_ fold change measured across all genes in each process in mono-culture or co-culture. ****: p-value < 0.0005, * < p-value < 0.05 according to pair-wise t-tests between the log_2_ fold change measured for each gene in mono-culture vs co-culture. Numerical values are reported in S9-S10 Files.

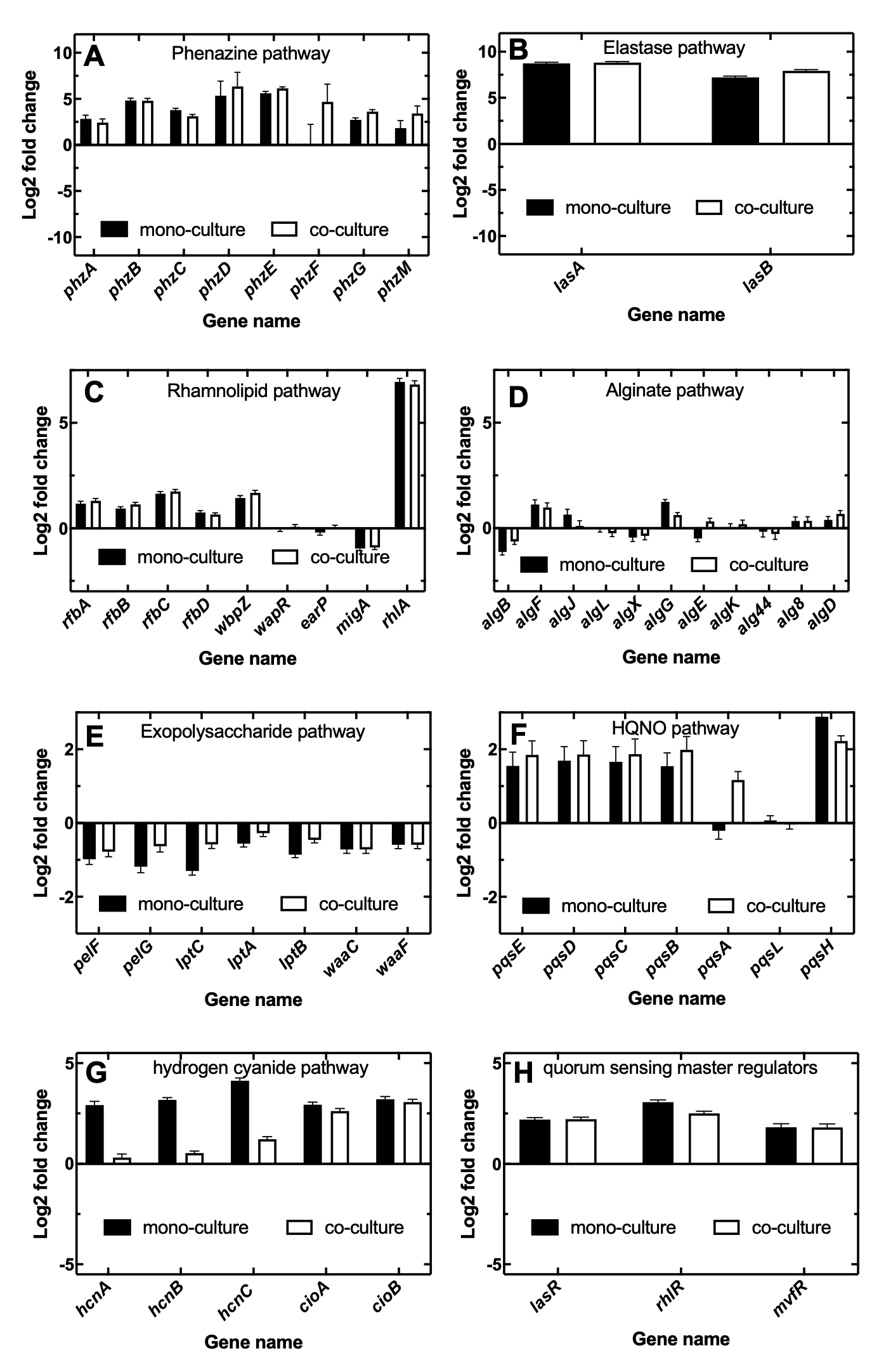

Figure S10 Differential expression of genes in the (a) phenazine, (b) elastase, (c) rhamnolipid, (d) alginate, (e) exopolysaccharide, (f) 2-heptyl-4-hydroxyquinoline-N-oxide (HQNO), (g), hydrogen cyanide pathways, or (h) quorum sensing master regulators at t = 10 h compared to t = 3 h for *P. aeruginosa* in mono-culture (filled bars) or co-culture (open bars). The bars and error bars report the mean and standard error of the log_2_ fold change calculated from transcriptomic measurements carried out in biological triplicate. Pair-wise t-tests between the log_2_ fold change measured for each gene in mono-culture vs co-culture returned not significant differences for all processes a part from the hydrogen cyanide and exopolysaccharide processes that were more upregulated and down-regulated in mono-culture, respectively. Numerical values are reported in S9-S10 Files.

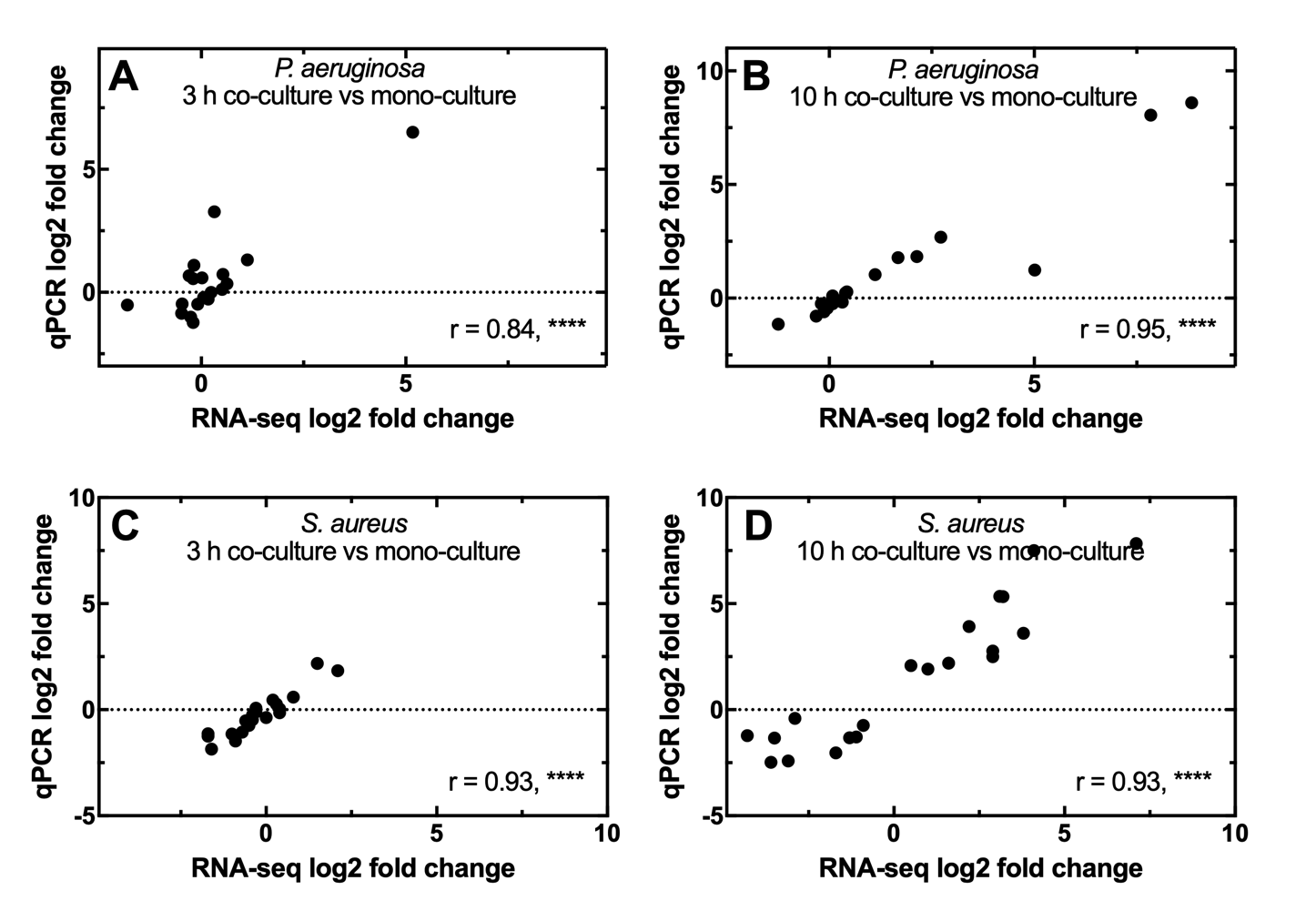

Figure S11 Comparison of the log2 fold changes obtained by RNA sequencing and qRT-PCR for twenty genes that were differentially regulated in *P. aeruginosa* after (a) 3 h or (b) 10 h of co-culture with *S. aureus* versus mono-culture or in in *S. aureus* after (c) 3 h or (d) 10 h of co-culture with *P. aeruginosa* versus mono-culture. Statistically significant correlations were calculated for each data set using Pearson correlation, with the calculate Pearson coefficients r and statistical significance indicated in each figure. Data points represent the average of RNA sequencing and qRT-PCR measurements carried out in biological triplicate. ****: p-value < 0.0005. Numerical values are reported in S11 File.

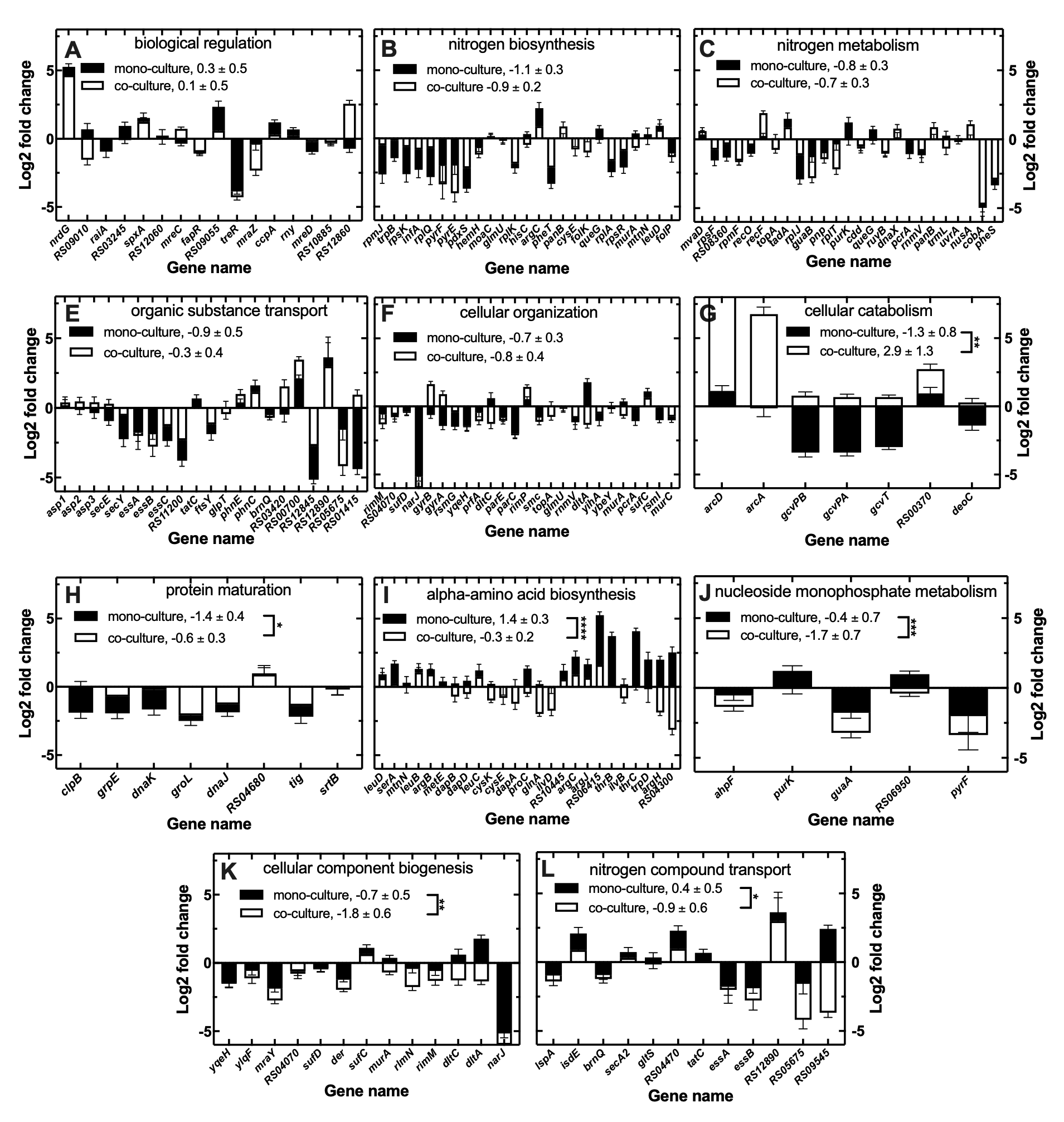

Figure S12 Differential expression of *S. aureus* ATCC25923 genes at t = 10 h compared to t = 3 h in mono-culture (filled bars) or co-culture (open bars) for biological processes that are significantly enriched in the comparisons of the transcriptomes of *S. aureus* in mono-culture vs co-culture with *P. aeruginosa* PA14 either at t = 3 h or at t = 10 h. The bars and error bars report the mean and standard error of the log_2_ fold change of each gene calculated from transcriptomic measurements carried out in biological triplicate. The numbers in each graph report the mean and standard error of the log_2_ fold change measured across all genes in each process in mono-culture or co-culture. ****: p-value < 0.0005, ***: p-value < 0.05, **: p-value < 0.01, * < p-value < 0.05 according to pair-wise t-tests between the log_2_ fold change measured for each gene in mono-culture vs co-culture. log_2_ fold change data are reported for all genes constituting the enrichment core in each process a part from nitrogen biosynthesis process. This process contained more than 25 enrichment core genes that would have been difficult to visualise. Therefore, 25 genes were randomly selected for this process for visualization, whereas numerical values for all genes are reported in S14-S15 Files.

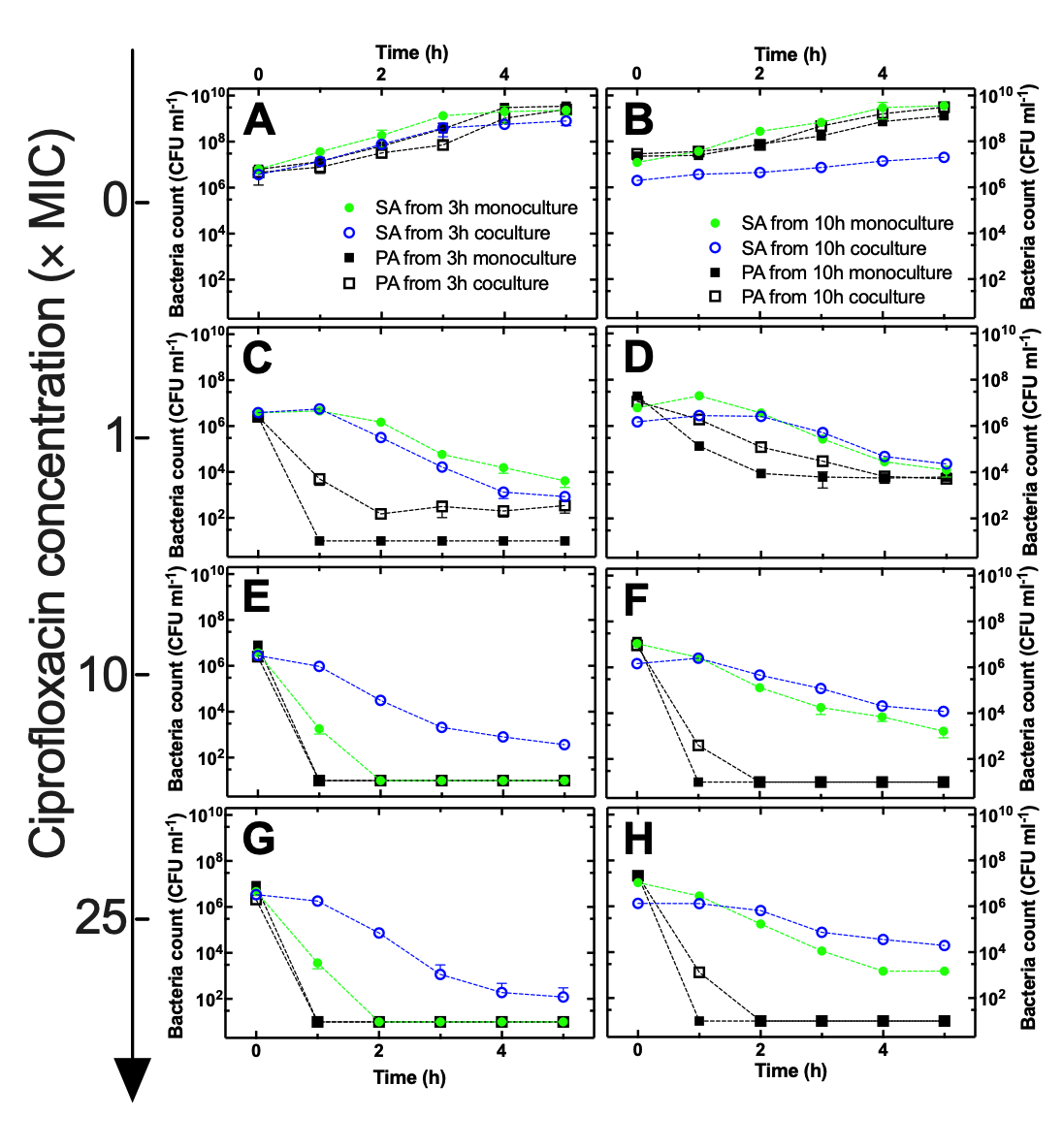

Figure S13 Temporal dependence of bacterial population size for *S. aureus* ATCC25923 in mono-culture (filled circles), *P. aeruginosa* PA14 in mono-culture (filled squares), *S. aureus* ATCC25923 (open circles) or *P. aeruginosa* PA14 (open squares) in co-culture in well-mixed flasks (A-B) untreated or treated with ciprofloxacin at a concentration of (C-D) 1×, (E-F) 10× or (G-H) 25× the MIC value measured against *S. aureus* ATCC25923. Bacteria were subjected to a 5 h ciprofloxacin treatment after being harvested from either (A, C, E, G) 3 h or (B, D, F, H) 10 h of mono-culture or co-culture. Data points are the mean and standard deviation of colony forming unit (CFU) measurements carried out in biological triplicate each consisting of technical duplicate. Numerical values for each replicate are reported in S16 File.

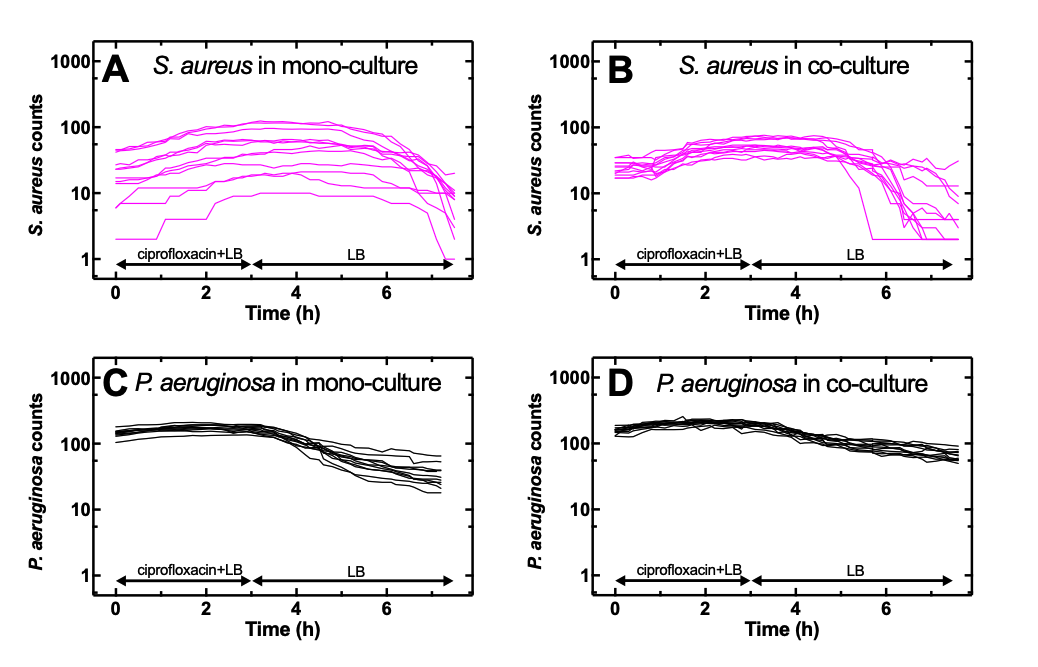

Figure S14 Temporal dependence of the number of *S. aureus* cells in 12 different microfluidic chambers (4 chambers from each biological replicate) harvested from (a) 10 h old mono-cultures or (b) 10 h old co-cultures and exposed first to ciprofloxacin at its MIC against *S. aureus* for 3 h and then LB medium for 4 h within each chamber. Corresponding temporal dependence of the number of *P. aeruginosa* cells from (c) 10 h old mono-cultures or (d) 10 h old co-cultures and exposed first to ciprofloxacin at its MIC against *S. aureus* for 3 h and then LB medium for 4 h within each chamber. Numerical values are reported in S17 File.

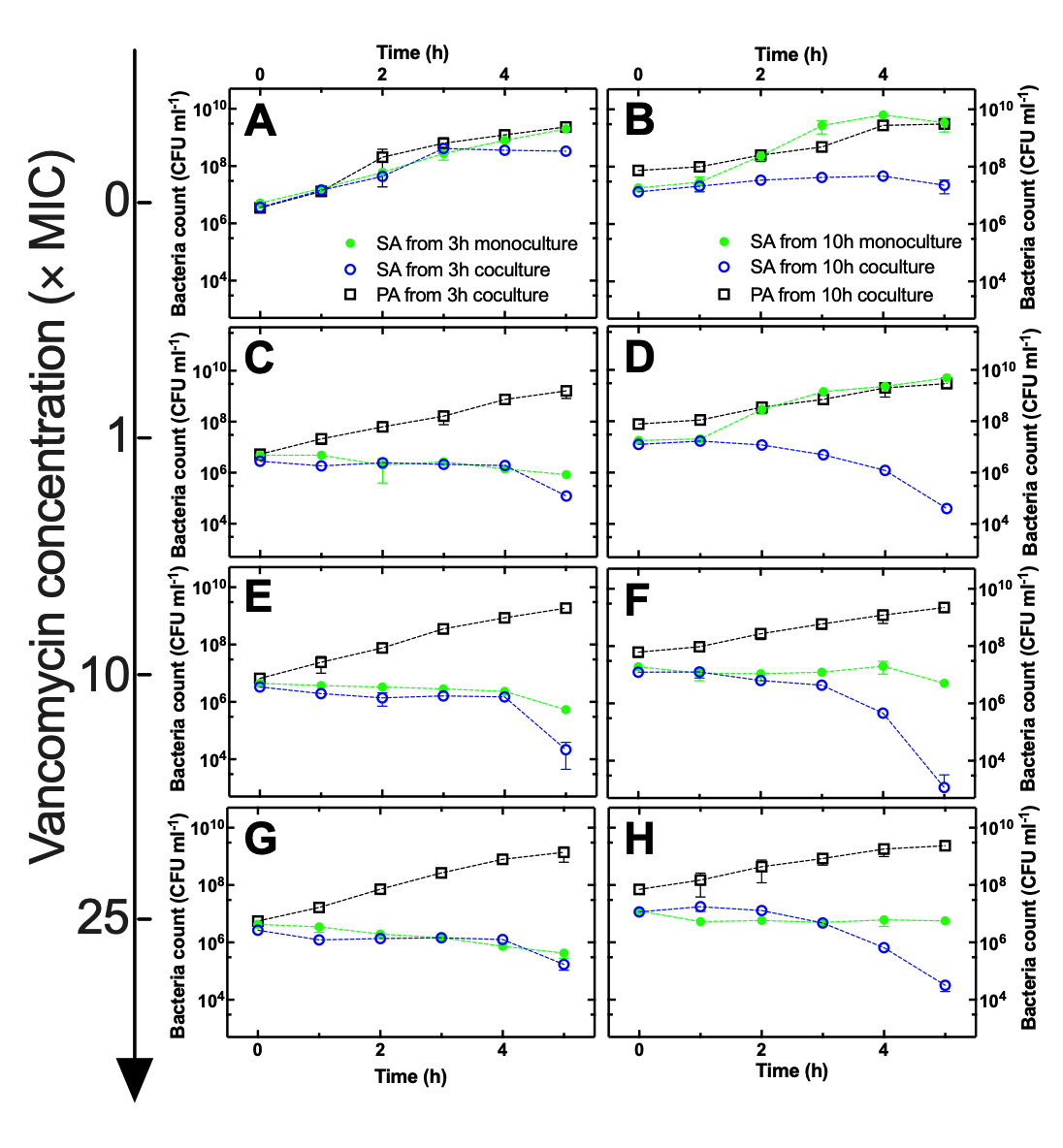

Figure S15 Temporal dependence of bacterial population size for *S. aureus* ATCC25923 in mono-culture (filled circles), *S. aureus* ATCC25923 (open circles) or *P. aeruginosa* PA14 (open squares) in co-culture in well-mixed flasks (A-B) untreated or treated with vancomycin at a concentration of (C-D) 1×, (E-F) 10× or (G-H) 25× the MIC value measured against *S. aureus* ATCC25923. Bacteria were subjected to a 5 h vancomycin treatment after being harvested from either (A, C, E, G) 3 h or (B, D, F, H) 10 h of mono-culture or co-culture. Data points are the mean and standard deviation of colony forming unit (CFU) measurements carried out in biological triplicate each consisting of technical duplicate. Numerical values for each replicate are reported in S18 File.

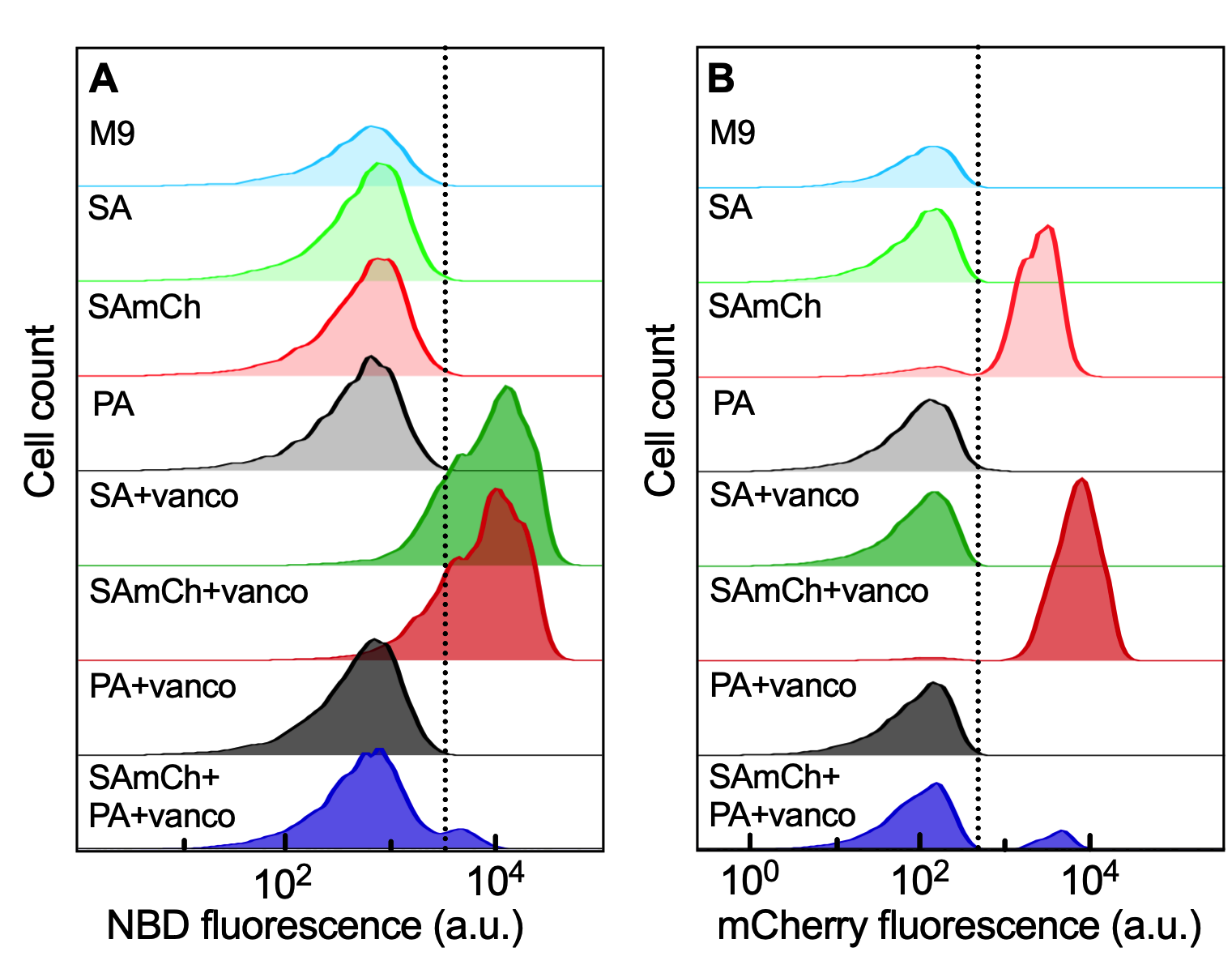

Figure S16 Single-cell distributions of fluorescence detected in (a) the FITC and (b) the ECD filter for the following experimental conditions from top to bottom: M9 minimal medium only, 10 h old *S. aureus* ATCC 25923 mono-culture, 10 h old *S. aureus* SH1000 mCherry mono-culture, 10 h old *P. aeruginosa* PA14 mono-culture, 10 h old *S. aureus* ATCC 25923 mono-culture incubated in vancomycin-NBD at a concentration of 16 µg mL^-1^ for 1 h, 10 h old *S. aureus* SH1000 mCherry mono-culture incubated in vancomycin-NBD at a concentration of 16 µg mL^-1^ for 1 h, 10 h old *P. aeruginosa* PA14 mono-culture incubated in vancomycin-NBD at a concentration of 16 µg mL^-1^ for 1 h or 10 h old co-cultures of *S. aureus* SH1000 mCherry and *P. aeruginosa* PA14 incubated in vancomycin-NBD at a concentration of 16 µg mL^-1^ for 1 h. Numerical values are reported in Data A in S19 File.

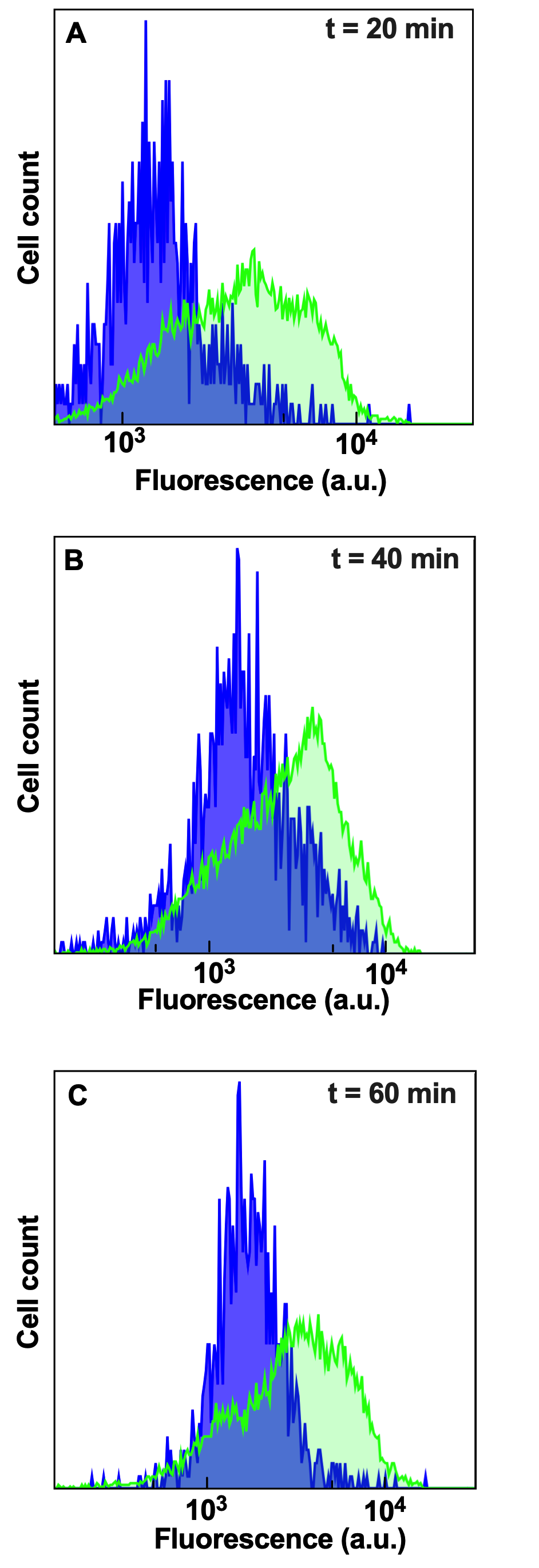

Figure S17 Accumulation of vancomycin-NBD in individual *S. aureus* SH1000 mCherry cells harvested from 10 h old mono- (green histogram) or co-culture (blue histogram), exposed to 16 µg mL^-1^ vancomycin-NBD for (a) 20 min, (b) 40 min or (c) 60 min and measured via flow cytometry. The data presented are representative of biological triplicate measurement reported in Data B in S19 File. In the co-culture experiments only data for cells expressing mCherry at levels above the ECD signal collected from M9 medium, i.e. only *S. aureus* SH1000 mCherry cells, are presented.

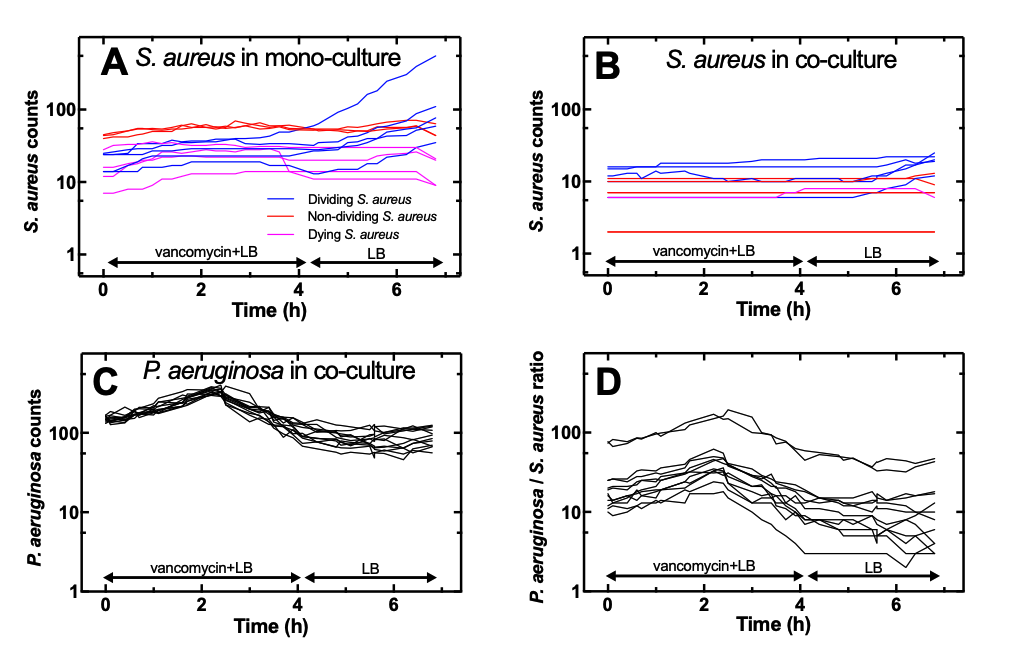

Figure S18 Temporal dependence of the number of *S. aureus* ATCC 25923 cells in 12 different microfluidic chambers (4 chambers from each biological replicate) harvested from (A) 10 h old mono-cultures or (b) 10 h old co-cultures with *P. aeruginosa* PA14 and exposed first to vancomycin at its MIC (against *S. aureus* ATCC 25923) for 3 h and then LB medium for 4 h within each chamber. (C) Corresponding temporal dependence of the number of *P. aeruginosa* PA14 cells. (D) Corresponding temporal dependence of the ratio of the number of *P. aeruginosa* cells over the number of *S. aureus* cells. Blue, red and magenta lines report temporal dependences for chambers in which the majority of *S. aureus* cells displayed sustained doubling, no doubling or death, respectively. Numerical values are reported in S20 File.

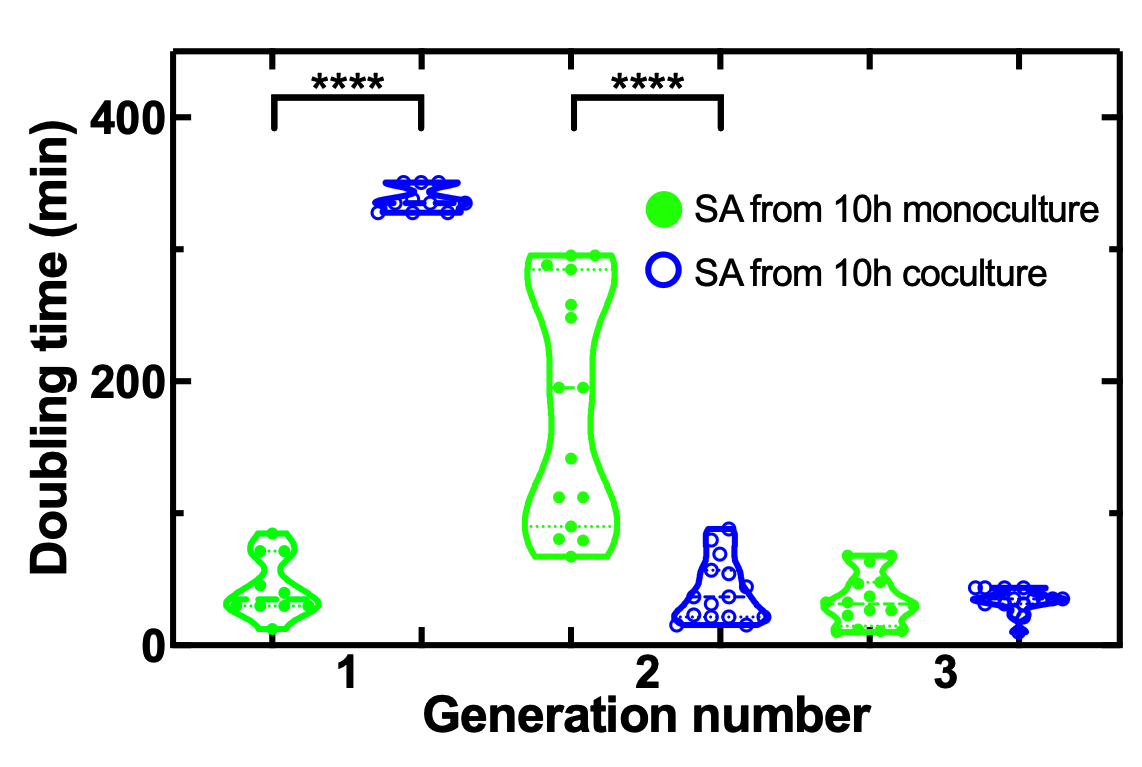

Figure S19 Dependence of doubling time on the generation number for *S. aureus* in mono-culture (green filled circles) or in co-culture with *P. aeruginosa* (open blue circles) within microfluidic chambers. Each point reports the doubling time of an individual bacterium with numerical values reported in Data F in S20 File. **** denotes a p-value < 0.0001.

| **Parameter** | ***S. aureus* ATCC25923 and *P. aeruginosa* PA14 (selective plates)** | | ***S. aureus* ATCC25923 and *P. aeruginosa* PA14 (LB plates)** | | ***S. aureus* RYC157 and *P. aeruginosa* RYC157** | | ***S. aureus* RYC157 and *P. aeruginosa* PA14** | | ***S. aureus* RYC165 and *P. aeruginosa* PA14** | |
| --- | --- | --- | --- | --- | --- | --- | --- | --- | --- | --- |
|  | ***S. aureus*** | ***P. aeruginosa*** | ***S. aureus*** | ***P. aeruginosa*** | ***S. aureus*** | ***P. aeruginosa*** | ***S. aureus*** | ***P. aeruginosa*** | ***S. aureus*** | ***P. aeruginosa*** |
| $\alpha$ | 1.74 | 1.20 | 1.79 | 1.13 | 1.76 | 0.360 | 1.68 | 1.31 | 1.68 | 1.14 |
| $\beta$ | 3.95 | 10.0 | 6.10 | 16.4 | 2.52 | 12.4 | 8.02 | 15.5 | 4.65 | 8.70 |
| $\gamma_{1}$ | 0.42 | 2.62 | 0.59 | 2.02 | 0.30 | 3.13 | 0.59 | 2.70 | 0.45 | 2.44 |
| $\gamma_{2}$ | 0.180 | 0.125 | 0.240 | 0.194 | 0.432 | 0.669 | 0.146 | 0.192 | 0.099 | 0.133 |
| $T$ | 3.56 | 6.22 | 4.32 | 7.17 | 5.40 | 37.1 | 4.65 | 6.54 | 4.11 | 6.22 |
| $I_{0}$ | 0.325 | 0.421 | 0.302 | 0.542 | 0.518 | 0.223 | 0.319 | 0.383 | 0.356 | 0.337 |
| $X_{0}$ | 2950000 | 1600000 | 1960000 | 2170000 | 1860000 | 1180000 | 2510000 | 655000 | 1730000 | 843000 |
| $\delta$ | $6.04\times{10}^{-11}$ | $6.04\times{10}^{-11}$ | $1.21\times{10}^{-10}$ | $1.21\times{10}^{-10}$ | $1.30\times{10}^{-10}$ | $1.30\times{10}^{-10}$ | $1.74\times{10}^{-10}$ | $1.74\times{10}^{-10}$ | $2.86\times{10}^{-10}$ | $2.86\times{10}^{-10}$ |

Table S1 Numerical estimates of the parameters used in the mathematical model employed to describe the population dynamics of co-cultures of different *S. aureus* and *P. aeruginosa* in well-mixed flasks obtained by fitting the model to the experimental data via least-squares regression.

| **Parameter** | **Mono-cultures** | **Co-cultures** |
| --- | --- | --- |
| $\alpha_{S}$ | 1.8 | 1.8 |
| $I_{0}\beta_{S}$ | 2.7 | 17.6 |
| $\gamma_{1,S}$ | 0.54 | 0.45 |

Table S2 Numerical estimates of the parameters used in the mathematical model employed to describe the population dynamics of mono- and co-cultures of *S. aureus* in microfluidic chambers obtained by fitting the model to the experimental data via least-squares regression.

| **Parameter** | **Co-cultures** |
| --- | --- |
| $\alpha_{P}$ | 1.17 |
| $\beta_{P}$ | 14.5 |
| $\gamma_{1}$ | 2.47 |
| $\epsilon_{C}$ | $7.70\times{10}^{-3}$ |
| $\epsilon_{Q}$ | 0.84 |

Table S3 Numerical estimates of the parameters used in the mathematical model employed to describe the population dynamics of co-cultures of *P. aeruginosa* in microfluidic chambers obtained by fitting the model to the experimental data via least-squares regression.

**Datasets legends**

S1 File. Data A: numerical values of the dynamics of *S. aureus* ATCC25923 population size in mono-culture performed in biological triplicate and technical duplicate using well-mixed flasks for growth and LB agar plates for enumeration. Data B: numerical values of the dynamics of *S. aureus* ATCC25923 population size in co-culture with *P. aeruginosa* PA14 performed in biological triplicate and technical duplicate using well-mixed flasks for growth and selective agar plates for enumeration. Data C: numerical values of the dynamics of *P. aeruginosa* PA14 population size in mono-culture performed in biological triplicate and technical duplicate using well-mixed flasks for growth and LB agar plates for enumeration. Data D: numerical values of the dynamics of *P. aeruginosa* PA14 population size in co-culture with *S. aureus* ATCC25923 performed in biological triplicate and technical duplicate using well-mixed flasks for growth and selective agar plates for enumeration. These numerical values are used in Figure 1A-B. Data E: numerical values of *S. aureus* ATCC25923 population size after 13h of mono-culture, or co-culture with *P. aeruginosa* PA14, *P. aeruginosa* PAO1, *P. aeruginosa* PA14 Δ*flgK*, *P. aeruginosa* PAO1 Δ*pvdD*Δ*pchEF* or *S. aureus* SH1000 mCherry population size after 13h of co-culture with *P. aeruginosa* PA14 Δ*flgK*. These numerical values are used in Figure S1. Data F-G: numerical values of the dynamics of *S. aureus* RYC157 or *P. aeruginosa* RYC157 population size in co-culture with *P. aeruginosa* RYC157 or *S. aureus* RYC157 performed in biological triplicate and technical duplicate using well-mixed flasks for growth and selective agar plates for enumeration. Data H-I: numerical values of the dynamics of *S. aureus* RYC157 or *P. aeruginosa* PA14 population size in co-culture with *P. aeruginosa* PA14 or *S. aureus* RYC157 performed in biological triplicate and technical duplicate using well-mixed flasks for growth and selective agar plates for enumeration. Data J-K: numerical values of the dynamics of *S. aureus* RYC165 or *P. aeruginosa* PA14 population size in co-culture with *P. aeruginosa* PA14 or *S. aureus* RY165 performed in biological triplicate and technical duplicate using well-mixed flasks for growth and selective agar plates for enumeration. These numerical values are used in Figure S2. Data L: number of dividing, non-dividing or lysing *S. aureus* ATCC25923 in co-culture with *P. aeruginosa* PA14 measured in biological triplicate and technical quadruplicate using microfluidic chambers. These numerical values are used in Figure 1E. Data M: temporal dependence of mCherry fluorescence value for individual dividing, non-dividing or lysing *S. aureus* ATCC25923 in co-culture with *P. aeruginosa* PA14 measured in biological triplicate and technical quadruplicate using microfluidic chambers. These numerical values are used in Figure 1D. Data N: numerical values of the dynamics of *S. aureus* ATCC25923 population size in mono-culture performed in biological triplicate and technical quadruplicate using microfluidic chambers for growth and enumeration. Data O: numerical values of the dynamics of *S. aureus* ATCC25923 population size in co-culture with *P. aeruginosa* PA14 performed in biological triplicate and technical quadruplicate. Data P: numerical values of the dynamics of *P. aeruginosa* PA14 population size in co-culture with *S. aureus* ATCC25923 performed in biological triplicate and technical quadruplicate. These numerical values are used in Figure 1G and S4. Data Q: doubling time of *S. aureus* ATCC25923 in mono-culture and in in co-culture with *P. aeruginosa* PA14. These numerical values are used in Figure 1F. Data R: numerical values of the dynamics of *S. aureus* ATCC25923 population size in co-culture with *P. aeruginosa* PA14 performed in biological triplicate and technical duplicate using well-mixed flasks for growth and selective agar plates for enumeration. Data S: corresponding numerical values of the dynamics of *P. aeruginosa* PA14 population size in co-culture with *S. aureus* ATCC25923 performed in biological triplicate and technical duplicate. These numerical values are used in Figure S5.

S2 File. Data A: principal component analysis values for triplicate transcriptome of *S. aureus* ATCC25923 or *P. aeruginosa* PA14 after 3 h or 10 h of mono-culture or co-culture. *S. aureus* transcriptome after 10 h of co-culture with *P. aeruginosa* was measured in six biological triplicate due to the relatively lower RNA concentration compared to the other bacterial samples. These numerical values are used in Figure S6. Data B: RNA transcript counts of *P. aeruginosa* genes after 3 h or 10 h of mono-culture or co-culture with *S. aureus*. Data C: RNA transcript counts of *S. aureus* genes after 3 h or 10 h of mono-culture or co-culture with *P. aeruginosa*. Analysis of these numerical values led to subsequent S Files and associated figures.

S3 File. Differential expression of *P. aeruginosa* genes after 3 h of co-culture with *S. aureus* compared to in mono-culture. Transcript base mean, log_2_ fold change, standard error of log_2_ fold change, p-value, adjusted p-value, gene identifier, gene and gene product are reported for each *P. aeruginosa* gene. These numerical values are used in Figure S7.

S4 File. Differential expression of *P. aeruginosa* genes after 10 h of co-culture with *S. aureus* compared to in mono-culture. Transcript base mean, log_2_ fold change, standard error of log_2_ fold change, p-value, adjusted p-value, gene identifier, gene and gene product are reported for each *P. aeruginosa* gene. These numerical values are used in Figure S7.

S5 File. Differential expression of *S. aureus* genes after 3 h of co-culture with *P. aeruginosa* compared to in mono-culture. Transcript base mean, log_2_ fold change, standard error of log_2_ fold change, p-value, adjusted p-value, gene identifier, gene and gene product are reported for each *S. aureus* gene. These numerical values are used in Figure S7.

S6 File. Differential expression of *S. aureus* genes after 10 h of co-culture with *P. aeruginosa* compared to in mono-culture. Transcript base mean, log_2_ fold change, standard error of log_2_ fold change, p-value, adjusted p-value, gene identifier, gene and gene product are reported for each *S. aureus* gene. These numerical values are used in Figure S7.

S7 File. Gene ontology enrichment analysis of differentially expressed *P. aeruginosa* genes after 3 h of co-culture with *S. aureus* compared to in mono-culture. Identifier, description, number of genes contained, enrichment score, normalized enrichment score, p-value, adjusted p-value, q-value, rank, leading edge and identifier of each gene are reported for each significantly enriched biological process. These numerical values are used in Figure 2A.

S8 File. Gene ontology enrichment analysis of differentially expressed *P. aeruginosa* genes after 10 h of co-culture with *S. aureus* compared to in mono-culture. Identifier, description, number of genes contained, enrichment score, normalized enrichment score, p-value, adjusted p-value, q-value, rank, leading edge and identifier of each gene are reported for each significantly enriched biological process. These numerical values are used in Figure 2B.

S9 File. Differential expression of *P. aeruginosa* genes after 10 h of mono-culture compared to after 3 h of mono-culture. Transcript base mean, log_2_ fold change, standard error of log_2_ fold change, p-value, adjusted p-value, gene identifier, gene and gene product are reported for each *P. aeruginosa* gene. These numerical values are used in Figure 2C, S8, S9 and S10.

S10 File. Differential expression of *P. aeruginosa* genes after 10 h of co-culture with *S. aureus* compared to after 3 h of co-culture. Transcript base mean, log_2_ fold change, standard error of log_2_ fold change, p-value, adjusted p-value, gene identifier, gene and gene product are reported for each *P. aeruginosa* gene. These numerical values are used in Figure 2C, S8, S9 and S10.

S11 File. Differential expression of 20 selected *P. aeruginosa* or *S. aureus* genes after 3 h or 10 h of co-culture with *S. aureus* or *P. aeruginosa* compared to the corresponding mono-cultures measured by both RNA-sequencing and quantitative real-time PCR. *gyrA* and *recN* were used as control genes, respectively. log_2_ fold change and standard error of log_2_ fold change were calculated from measurements carried out in biological triplicate. These numerical values are used in Figure S11.

S12 File. Gene ontology enrichment analysis of differentially expressed *S. aureus* genes after 3 h of co-culture with *P. aeruginosa* compared to mono-culture. Identifier, description, number of genes contained, enrichment score, normalized enrichment score, p-value, adjusted p-value, q-value, rank, leading edge and identifier of each gene are reported for each significantly enriched biological process. These numerical values are used in Figure 2D.

S13 File. Gene ontology enrichment analysis of differentially expressed *S. aureus* genes after 10 h of co-culture with *P. aeruginosa* compared to mono-culture. Identifier, description, number of genes contained, enrichment score, normalized enrichment score, p-value, adjusted p-value, q-value, rank, leading edge and identifier of each gene are reported for each significantly enriched biological process. These numerical values are used in Figure 2E.

S14 File. Differential expression of *S. aureus* genes after 10 h of mono-culture compared to after 3 h of mono-culture. Transcript base mean, log_2_ fold change, standard error of log_2_ fold change, p-value, adjusted p-value, gene identifier, gene and gene product are reported for each *S. aureus* gene. These numerical values are used in Figure 2F and S12.

S15 File. Differential expression of *S. aureus* genes after 10 h of co-culture with *P. aeruginosa* compared to after 3 h of co-culture. Transcript base mean, log_2_ fold change, standard error of log_2_ fold change, p-value, adjusted p-value, gene identifier, gene and gene product are reported for each *S. aureus* gene. These numerical values are used in Figure 2F and S12.

S16 File. Data A: numerical values of the dynamics of the population size of *S. aureus* harvested from 3 h or 10 h old mono-cultures and regrown in LB medium for 5 h performed in biological triplicate and technical duplicate using well-mixed flasks for growth and LB agar plates for enumeration. Data B-D: numerical values of the dynamics of the population size of *S. aureus* harvested from 3 h or 10 h old mono-cultures and treated with ciprofloxacin at its MIC (against *S. aureus*), or 10× its MIC, or 25× its MIC in LB medium for 5 h performed in biological triplicate and technical duplicate using well-mixed flasks for treatment and LB agar plates for enumeration. Data E: numerical values of the dynamics of the population size of *S. aureus* harvested from 3 h or 10 h old co-cultures with *P. aeruginosa* and regrown in LB medium for 5 h performed in biological triplicate and technical duplicate using well-mixed flasks for growth and LB agar plates for enumeration. Data F-H: numerical values of the dynamics of the population size of *S. aureus* harvested from 3 h or 10 h old co-cultures and treated with ciprofloxacin at its MIC (against *S. aureus*), or 10× its MIC, or 25× its MIC in LB medium for 5 h performed in biological triplicate and technical duplicate using well-mixed flasks for treatment and LB agar plates for enumeration. Data I: numerical values of the dynamics of the population size of *P. aeruginosa* harvested from 3 h or 10 h old co-cultures and regrown in LB medium for 5 h performed in biological triplicate and technical duplicate using well-mixed flasks for growth and LB agar plates for enumeration. Data J-L: numerical values of the dynamics of the population size of *P. aeruginosa* harvested from 3 h or a 10 h old co-cultures and treated with ciprofloxacin at its MIC (against *S. aureus*), or 10× its MIC, or 25× its MIC in LB medium for 5 h performed in biological triplicate and technical duplicate using well-mixed flasks for treatment and LB agar plates for enumeration. Data M: numerical values of the dynamics of the population size of *P. aeruginosa* harvested from 3 h or 10 h old mono-cultures and regrown in LB medium for 5 h performed in biological triplicate and technical duplicate using well-mixed flasks for growth and LB agar plates for enumeration. Data N-P: numerical values of the dynamics of the population size of *P. aeruginosa* harvested from 3 h or 10 h old mono-cultures and treated with ciprofloxacin at its MIC (against *S. aureus*), or 10× its MIC, or 25× its MIC in LB medium for 5 h performed in biological triplicate and technical duplicate using well-mixed flasks for treatment and LB agar plates for enumeration. These numerical values are used in Figure 3AB and Figure S13.

S17 File. Data A: numerical values of the dynamics of *S. aureus* ATCC25923 population size in mono-culture exposed to ciprofloxacin at 0.5 µg mL^-1^ for 4 h and LB medium for 3 h performed in biological triplicate and technical quadruplicate using microfluidic chambers for growth and enumeration. Data B: corresponding number of dividing, non-dividing or lysing *S. aureus*. Data C: numerical values of the dynamics of *S. aureus* ATCC25923 population size in co-culture with *P. aeruginosa* PA14 exposed to ciprofloxacin at 0.5 µg mL^-1^ for 3 h and LB medium for 4 h performed in biological triplicate and technical quadruplicate using microfluidic chambers for growth and enumeration. Data D: corresponding number of dividing, non-dividing or lysing *S. aureus*. Data E: corresponding numerical values of the dynamics of *P. aeruginosa* PA14 population size in co-culture with *S. aureus*. Data F: numerical values of the dynamics of *P. aeruginosa* PA14 population size in mono-culture exposed to ciprofloxacin at 0.5 µg mL^-1^ for 4 h and LB medium for 3 h performed in biological triplicate and technical quadruplicate using microfluidic chambers for growth and enumeration. These numerical values are used in Figure 3DE and Figure S14.

S18 File. Data A: numerical values of the dynamics of the population size of *S. aureus* harvested from 3 h or 10 h old mono-cultures and regrown in LB medium for 5 h performed in biological triplicate and technical duplicate using well-mixed flasks for growth and LB agar plates for enumeration. Data B-D: numerical values of the dynamics of the population size of *S. aureus* harvested from 3 h or 10 h old mono-cultures and treated with vancomycin at its MIC (against *S. aureus*), or 10× its MIC, or 25× its MIC in LB medium for 5 h performed in biological triplicate and technical duplicate using well-mixed flasks for treatment and LB agar plates for enumeration. Data E: numerical values of the dynamics of the population size of *S. aureus* harvested from 3 h or 10 h old co-cultures and regrown in LB medium for 5 h performed in biological triplicate and technical duplicate using well-mixed flasks for growth and LB agar plates for enumeration. Data F-H: numerical values of the dynamics of the population size of *S. aureus* harvested from 3 h or 10 h old co-cultures and treated with vancomycin at its MIC, or 10× its MIC, or 25× its MIC in LB medium for 5 h performed in biological triplicate and technical duplicate using well-mixed flasks for treatment and LB agar plates for enumeration. Data I: numerical values of the dynamics of the population size of *P. aeruginosa* harvested from 3 h or 10 h old co-cultures and regrown in LB medium for 5 h performed in biological triplicate and technical duplicate using well-mixed flasks for growth and LB agar plates for enumeration. Data J-L: numerical values of the dynamics of the population size of *P. aeruginosa* harvested from 3 h or 10 h old co-cultures and treated with vancomycin at its MIC (against *S. aureus*), or 10× its MIC, or 25× its MIC in LB medium for 5 h performed in biological triplicate and technical duplicate using well-mixed flasks for treatment and LB agar plates for enumeration. These numerical values are used in Figure 4AB and Figure S15.

S19 File. Data A: Single-cell distributions of fluorescence detected in the FITC and the ECD filter for the following experimental conditions. M9 minimal medium only, 10 h old *S. aureus* ATCC 25923 mono-culture, 10 h old *S. aureus* SH1000 mCherry mono-culture, 10 h old *P. aeruginosa* PA14 mono-culture, 10 h old *S. aureus* ATCC 25923 mono-culture incubated in vancomycin-NBD at a concentration of 2 µg mL^-1^ for 1 h, 10 h old *S. aureus* SH1000 mCherry mono-culture incubated in vancomycin-NBD at a concentration of 2 µg mL^-1^ for 1 h, 10 h old *P. aeruginosa* PA14 mono-culture incubated in vancomycin-NBD at a concentration of 2 µg mL^-1^ for 1 h or 10 h old co-cultures of *S. aureus* SH1000 mCherry and *P. aeruginosa* PA14 incubated in vancomycin-NBD at a concentration of 2 µg mL^-1^ for 1 h. Data B: single-cell distributions of fluorescence detected in the FITC filter in individual *S. aureus* cells harvested from 10 h old mono- or co-culture, exposed to 2 µg mL^-1^ vancomycin-NBD for 20 min, 40 min or 60 min. In the co-culture experiments only data for cells expressing mCherry at levels above the ECD signal collected from M9 medium, i.e. only *S. aureus* cells, are presented. These numerical values are used in Figure 4CD, Figure S16 and S17.

S20 File. Data A: numerical values of the dynamics of *S. aureus* ATCC25923 population size in mono-culture exposed to vancomycin at 1 µg mL^-1^ for 4 h and LB medium for 3 h performed in biological triplicate and technical quadruplicate using microfluidic chambers for growth and enumeration. Data B: corresponding number of dividing, non-dividing or lysing *S. aureus*. Data C: numerical values of the dynamics of *S. aureus* ATCC25923 population size in co-culture with *P. aeruginosa* PA14 exposed to vancomycin at 1 µg mL^-1^ for 4 h and LB medium for 3 h performed in biological triplicate and technical quadruplicate using microfluidic chambers for growth and enumeration. Data D: corresponding number of dividing, non-dividing or lysing *S. aureus*. Data E: numerical values of the dynamics of *P. aeruginosa* PA14 population size in co-culture with *S. aureus* ATCC25923 exposed to vancomycin at 1 µg mL^-1^ for 4 h and LB medium for 3 h performed in biological triplicate and technical quadruplicate. All these numerical values are used in Figure 3FG and Figure S18. Data F: corresponding numerical values of the dependence of doubling time on the generation number for *S. aureus* in mono-culture or in co-culture with *P. aeruginosa* during and after treatment with vancomycin within microfluidic chambers in biological triplicate and technical quadruplicate. These numerical values are used in Figure S19.
